# Synaptic input-selective homeostasis safeguards developing cortical neurons toward setpoint activity

**DOI:** 10.64898/2026.08.01.729386

**Authors:** Zhao-Lin Cai, Colleen M. Longley, Dongwon Lee, Wu Chen, Mingshan Xue

## Abstract

Homeostatic plasticity stabilizes brain function by maintaining neuronal activity around a setpoint. Although homeostatic mechanisms are well characterized in mature circuits, it remains unknown how developing neurons, which progress from near silence at birth to sustained activity, establish and safeguard their setpoint activity levels against perturbations. Here we show that activity setpoints are intrinsic properties of neuronal identity actively achieved by cell-autonomous, targeted synaptic remodeling during development. Cortical layer 2/3 pyramidal cells were sparsely silenced from their birth by overexpressing the inward-rectifier potassium channel Kir2.1 to chronically suppress their excitability. These Kir2.1-expressing neurons, however, progressively overcame this perturbation during postnatal development and ultimately reached the activity levels comparable to neighboring control neurons. This recovery was accompanied by enhanced synaptic excitation and reduced inhibition that did not follow the classical global multiplicative synaptic scaling. Instead, excitation from infragranular layers, rather than layer 4 and layer 2/3, was selectively strengthened through increasing quantal amplitudes and unsilencing silent synapses that required the insertion of GluA2-containing AMPA receptors. Inhibition recruited by excitatory inputs was selectively reduced at specific pathways. These results identify a synaptic input-selective homeostatic program enabling cortical neurons to reach activity setpoints, revealing a new mechanism contributing to neurodevelopmental robustness.

## Introduction

The mammalian cerebral cortex achieves a remarkable balance between flexibility and stability. Dynamic neuronal activity enables flexible functions, as activity levels of neurons in the mature cortex span several orders of magnitude (*1*). Despite such a wide range of activity levels across neurons, the average firing rate of individual neurons over time tends to remain within a targeted range, or setpoint (*2–4*). This long-term stability of activity levels is believed to be crucial for maintaining cortical functions and result from multiple homeostatic mechanisms that regulates synaptic inputs and neuronal excitability in a compensatory manner to stabilize neuronal activity around the setpoints in response to perturbations (*5–8*). Not surprisingly, impaired homeostatic mechanisms are involved in the pathogeneses of neurological and psychiatric disorders (*9–12*).

Prior studies show that when the activity of a neuron is perturbed *in vitro* or *in vivo*, its synaptic inputs can be modified to help restore its activity to the previously established setpoint (*13–18*). Multiple homeostatic synaptic mechanisms have been identified. For example, the expression levels of postsynaptic receptors can be regulated by altering neuronal activity (*13*, *19–21*).

Synaptic scaling, the most common form of global receptor regulation, enhances or reduces all of a neuron’s synaptic inputs in a multiplicative manner, preserving the relative synaptic efficacies among different synapses (*22*). Receptor levels can also be locally regulated at specific synapses (*23–26*). Furthermore, neurotransmitter release and synapse numbers can be homeostatically adjusted in response to changes in activity (*14*, *18*, *27–33*).

While much is known about homeostatic synaptic plasticity in neurons where setpoint activity levels are already established, it is unclear how neurons acquire setpoint levels of activity and respond to activity perturbation in early development prior to establishing the setpoints.

Postmitotic cortical neurons gradually acquire the ability to generate action potentials, with the expressions of voltage-gated sodium and potassium channels starting from late embryonic stages in rodents (*34–36*). As developing neurons are gaining functional firing activity, they can encounter various perturbations ranging from internal genetic errors or developmental changes of neurons to external insults or experience-induced circuit modifications. Thus, it is possible that neurons ultimately settle on setpoint activity levels that are an emergent property of the circuits incorporating the impacts of perturbations. Alternatively, neurons may employ mechanisms counteracting perturbations to achieve targeted activity levels that are encoded as part of their identities. To explore this question, we sought to disrupt the activity of a small subset of neurons from their birth in an otherwise normal cortex and determine their responses to the ongoing perturbation.

## Results

### Developing cortical neurons establish setpoint activity despite persistent perturbation

To perturb neuronal activity *in vivo*, we used *in utero* electroporation to transfect plasmids overexpressing an inwardly rectifying potassium channel Kir2.1 into neural progenitor cells that give rise to a small subset (Extended Data Fig. 1a–c, 7.5 ± 0.8%, mean ± s.e.m., *n* = 13 slices from 5 mice) of layer 2/3 pyramidal cells in the mouse primary visual cortex (V1) (*37*). Whole-cell current clamp recordings in acute brain slices revealed a drastic reduction in the intrinsic excitability of Kir2.1-overexpressing cells (Kir2.1 neurons) throughout the development, as they had lower input resistances and larger rheobase currents than nearby untransfected layer 2/3 pyramidal cells (control neurons) (Extended Data Fig. 1d–h). The excitability of control neurons remained similar to that of neurons in non-electroporated control mice (Extended Data Fig. 2a– e). Thus, sparse Kir2.1 overexpression causes a cell-autonomous reduction in the excitability of layer 2/3 pyramidal cells.

We performed two-photon imaging-guided loose-patch recordings of spontaneous and visually evoked activity from Kir2.1 and nearby control neurons *in vivo* (Fig. 1a,b). Spikes were recorded in all control neurons regardless of age (Fig. 1c–g). In contrast, before postnatal day 13 (P13) all recorded Kir2.1 neurons (*n* = 11, 6 at P10 and 5 at P12) showed zero spikes. However, on P13, Kir2.1 neurons started to show neuronal activity, as spikes were recorded in 2 out of 6 Kir2.1 neurons (Fig. 1c, e–g). During postnatal week 3 (P15–21), Kir2.1 neurons became more active, as most of Kir2.1 neurons fired spikes spontaneously or in response to visual stimuli, although their spike rates remained significantly lower than those of control neurons (Fig. 1e–g).

**Figure 1.**
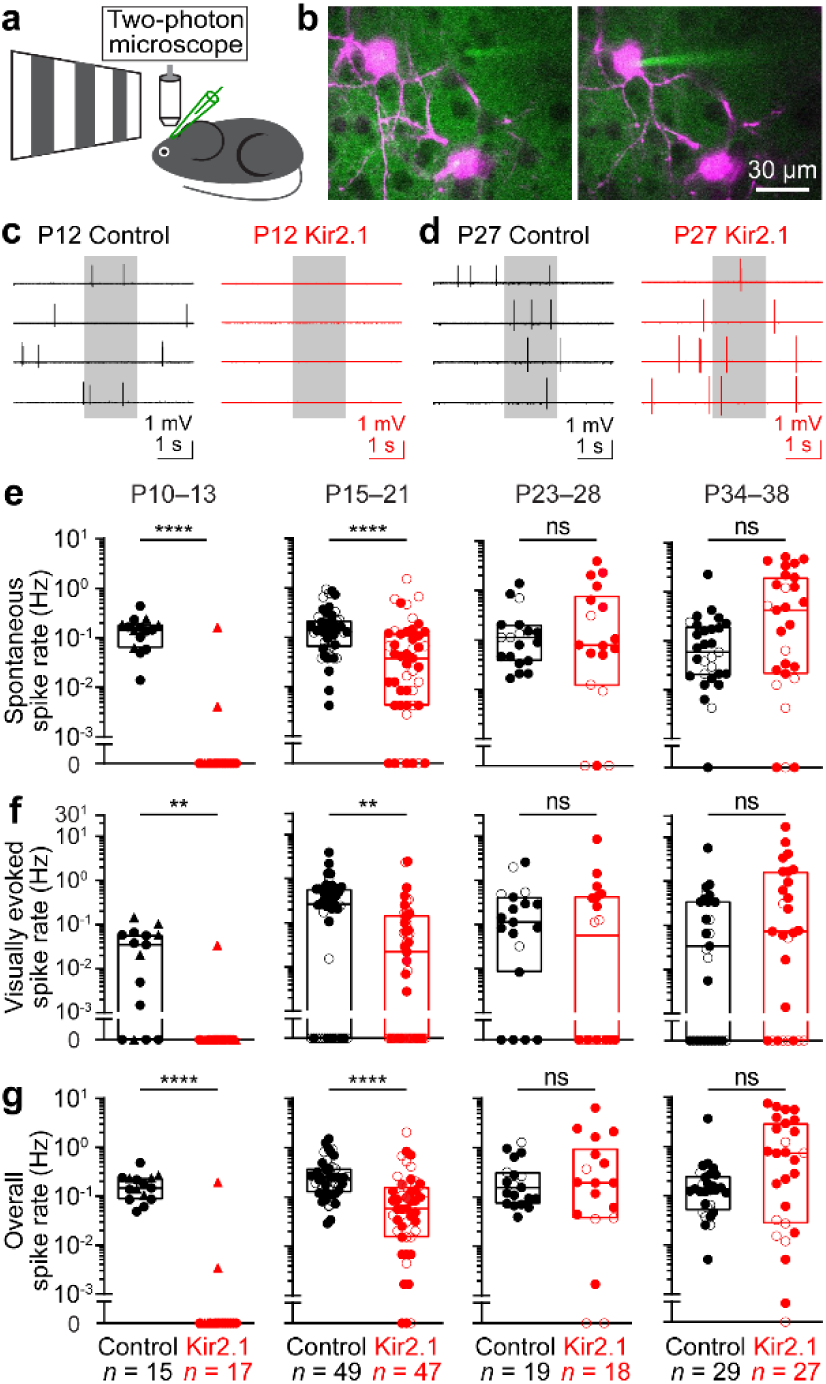
Cortical neurons overcome reduced excitability to reach setpoint activity levels during development. **(a)** Schematic of *in vivo* experiments. **(b)** A control neuron (left, dark shadow) and a Kir2.1 neuron (right, magenta) were sequentially recorded with Alexa Fluor 488-filled micropipettes (green). **(c)** Recordings from a control and a Kir2.1 neuron for spontaneous and visual-evoked spikes at postnatal day 12 (P12). Grey box, visual stimulation period. Note zero spikes in Kir2.1 neuron. (**d**) Similar to **c**, but at P27. Note similar spike rates in Kir2.1 neuron as control neuron. **(e**‒**g)** Summary data of spontaneous (e), visual-evoked (f) and overall (g) spike rates of control and Kir2.1 neurons at postnatal weeks 2 (P10‒13), 3 (P15‒21), 4 (P23‒ 28) and 5 (P34‒38). The center line of box plot indicates the median, and the box limits denote the 25‒75th percentiles. Each filled (male) or open (female) symbol represents a neuron. Kir2.1 neurons had no spikes at P10–12 (circle symbols) and started generating spikes from P13, as 2 out of 6 recorded cells had spikes (triangle symbols). Their spike rates remained significantly lower but were closer to control levels during postnatal week 3 and ultimately reached control levels in postnatal weeks 4 and 5. The numbers of recorded neurons (*n*) are indicated in the panels. ns, *P* > 0.05; **, *P* < 0.01; ****, *P* < 0.0001.

Subsequently, the activity of Kir2.1 neurons continued to increase and reached levels comparable to control neurons by postnatal weeks 4–5 (Fig. 1d,e–g), despite their severely reduced excitability (Extended Data Fig. 1f–h). These results indicate that cortical layer 2/3 pyramidal cells can gradually overcome the reduced excitability that persists since their birth and eventually achieve targeted activity levels *in vivo*.

### Nonuniform increase of synaptic excitation and decrease of inhibition

To determine the mechanisms by which Kir2.1 neurons overcame the reduced excitability to generate targeted levels of activity, we recorded miniature excitatory postsynaptic currents (mEPSCs), miniature inhibitory postsynaptic currents (mIPSCs), spontaneous EPSCs (sEPSCs), and spontaneous IPSCs (sIPSCs) in brain slices. Under the experimental conditions, these EPSCs are largely mediated by alpha-amino-3-hydroxy-5-methyl-4-isoxazolepropionic acid (AMPA) receptors and IPSCs by gamma-aminobutyric acid type A (GABAA) receptors. At postnatal week 3, the amplitudes, but not the frequencies, of mEPSCs were increased in Kir2.1 neurons, whereas the frequencies and amplitudes of mIPSCs were decreased (Fig. 2a–e). At postnatal week 5, the frequencies and amplitudes of mEPSCs were both increased in Kir2.1 neurons, but only the amplitudes, not the frequencies, of mIPSCs were decreased as compared to control neurons (Fig. 2h–l). Similar results were obtained for sEPSCs and sIPSCs except that at postnatal week 3, the frequencies of sEPSCs were also increased in Kir2.1 neurons (Extended Data Fig. 3). Importantly, the frequencies and amplitudes of sEPSCs and sIPSCs in control neurons remained similar to those in neurons of non-electroporated control mice (Extended Data Fig. 2f–j), indicating that the synaptic changes in Kir2.1 neurons occurs in a cell-autonomous manner.

**Figure 2.**
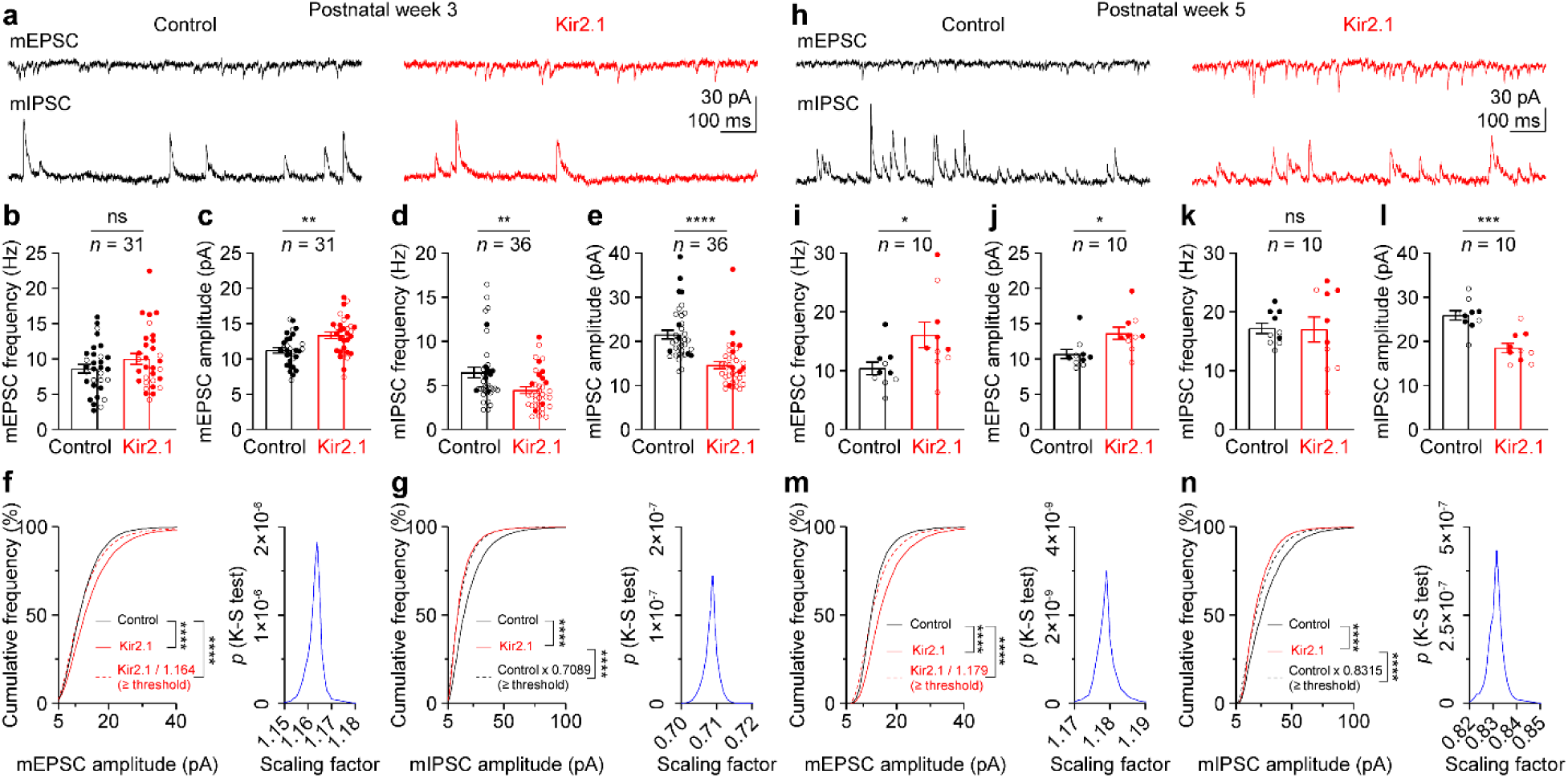
Nonuniform increase of mEPSCs and decrease of mIPSCs in Kir2.1 neurons. **(a)** mEPSCs and mIPSCs from a control and Kir2.1 neurons at postnatal week 3. **(b**‒**e)** Summary data of frequency and amplitude of mEPSCs (**b,c**) and mIPSCs (**d,e**) at postnatal week 3. Each filled (male) or open (female) symbol represents a neuron. Data are mean ± s.e.m. Note increased mEPSC amplitude and decreased mIPSC frequency and amplitude in Kir2.1 neurons. **(f)** Left, Cumulative distribution of mEPSC amplitudes from control and Kir2.1 neurons, with 500 randomly selected events per neuron. Upon dividing Kir2.1 mEPSC amplitudes by 1.164 and excluding subthreshold values (< 5 pA), the resulting scaled Kir2.1 distribution shows maximal overlap with the control distribution, yet remains significantly different. Right, the *P* values for the comparisons between control and scaled Kir2.1 distributions with different scaling factors. The *P* value is the largest with the scaling factor 1.164 used in the left panel. **(g)** As in **f**, but for distribution of mIPSC amplitudes. The best scaled control distribution is significantly different from the Kir2.1 distribution. **(h**‒**n)** As in **a**‒**g**, but for postnatal week 5. Note increased mEPSC frequency and amplitude and decreased mIPSC amplitude in Kir2.1 neurons. The best scaled distributions of Kir2.1 mEPSC amplitudes and control mIPSC amplitudes are significantly different from the control and Kir2.1 distributions, respectively. The numbers of pairs of recorded neurons (*n*) are indicated in the panels. ns, *P* > 0.05; *, *P* < 0.05; **, *P* < 0.01; ***, *P* < 0.001; ****, *P* < 0.0001.

These results indicate that both enhanced synaptic excitation and reduced synaptic inhibition contribute to promoting neuronal activity in Kir2.1 neurons.

To determine if the changes of synaptic strengths align with the multiplicative homeostatic synaptic scaling, we performed scaling analysis of the cumulative distributions of mEPSC and mIPSC amplitudes between Kir2.1 and control neurons by searching for a single scaling factor that best matched the two distributions (*38*). However, the best scaled distributions of mEPSC and mIPSC amplitudes still differed significantly from their corresponding counterparts (Fig. 2f,g,m,n). Similar results were obtained by the conventional test of multiplicative scaling (*13*)(Extended Data Fig. 4). Thus, these results demonstrate a nonuniform scaling of synaptic strengths at both excitatory and inhibitory synapses and suggest that different synaptic inputs were differentially adjusted in Kir2.1 neurons.

The nonuniform down-regulation of inhibition is consistent with the previous results that reducing the excitability of pyramidal cells selectively decreased their synaptic inhibition from parvalbumin-expressing, but not somatostatin-expressing, interneurons (*37*, *39*). However, the nonuniform up-regulation of excitation is unexpected, given that uniform synaptic scaling is considered the hallmark homeostatic excitatory synaptic plasticity mechanism in response to reduced neuronal activity (*8*, *22*). Layer 2/3 pyramidal cells in the V1 receive excitation from multiple sources including local inputs from different cortical layers and long-range inputs from the thalamus and other cortical areas (*40–45*). Thus, we sought to further investigate whether and how distinct excitatory inputs, along with the inhibition that they recruit, contribute to promoting neuronal activity in Kir2.1 neurons.

### Selective increase of synaptic excitation from local infragranular layers

We used adeno-associated viruses (AAV) or plasmids to selectively express excitatory channelrhodopsins (ChR2 or ReaChR) in different types of presynaptic neurons (see Methods) for photoactivating specific excitatory inputs while simultaneously recording pairs of Kir2.1 and nearby control neurons in brain slices. We first examined the ipsilateral local excitatory inputs from layer 2/3 and layer 4, the two main sources of excitation to layer 2/3 pyramidal cells (*40*, *41*, *46*). At postnatal weeks 3 and 5, layer 2/3-evoked EPSCs, disynaptic IPSCs, and the ratios of EPSC and IPSC amplitudes (E/I ratios) were similar between Kir2.1 and control neurons (Fig. 3a–3e). Layer 4-evoked EPSCs were also similar between Kir2.1 and control neurons at postnatal weeks 3 and 5, whereas disynaptic IPSCs were decreased in Kir2.1 neurons (Fig. 3f– 3i), resulting in an increase in layer 4-evoked E/I ratios in Kir2.1 neurons (Fig. 3j). The reduction of disynaptic IPSCs was due to the decreased inhibition from parvalbumin-expressing interneurons (*37*), but it is surprising that neither of the two major excitatory inputs to layer 2/3 pyramidal cells is homeostatically increased to promote activity in Kir2.1 neurons *in vivo*. Thus, we turned to other sources of excitation.

**Figure 3.**
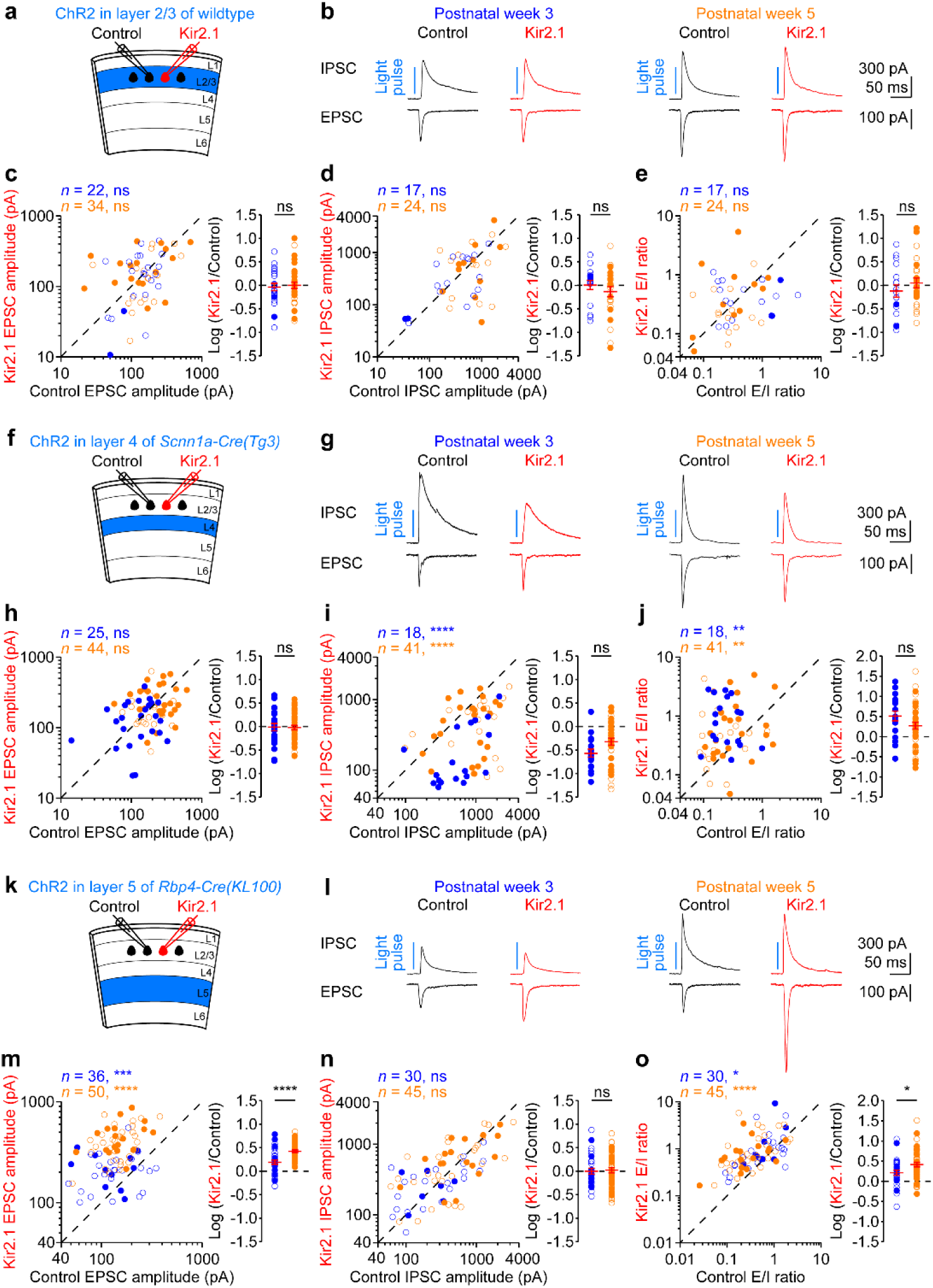
Local excitation from layer 5 but not layer 2/3 or layer 4 is increased in Kir2.1 neurons. **(a)** Schematic of slice experiments in **b** with Kir2.1 in a subset of layer 2/3 pyramidal cells and ChR2 in another subset of layer 2/3 pyramidal cells. **(b)** Monosynaptic EPSCs and disynaptic IPSCs from a pair of simultaneously recorded control and Kir2.1 neurons in response to photostimulation. **(c)** Left, summary data of layer 2/3-evoked EPSC amplitudes in control and Kir2.1 neurons at postnatal weeks 3 (blue) and 5 (orange). Each filled (male) or open (female) symbol represents a pair of control and Kir2.1 neurons. Right, logarithm of the ratios between EPSC amplitudes from pairs of Kir2.1 and control neurons. Red, mean ± s.e.m. **(d,e)** As in **c**, but for IPSC amplitudes and E/I ratios. Note similar EPSC and IPSC amplitudes and E/I ratios between control and Kir2.1 neurons. **(f**‒**j)** As in **a**‒**e**, but for ChR2 in layer 4 excitatory neurons. EPSC, IPSC amplitudes, and E/I ratio at postnatal week 3 are derived from previous work (*37*). Note similar EPSC amplitudes between Kir2.1 and control neurons, smaller IPSC amplitudes, and larger E/I ratios in Kir2.1 neurons. The decrease of IPSC amplitude and increase of E/I ratio in Kir2.1 neurons are similar between the two ages. **(k**‒**o)** As in **a**‒**e**, but for ChR2 in layer 5 excitatory neurons. Note similar IPSC amplitudes between Kir2.1 and control neurons, larger EPSC amplitudes, and larger E/I ratios in Kir2.1 neurons. The increase of EPSC amplitude and E/I ratio in Kir2.1 neurons is larger at postnatal week 5 than week 3. The numbers of pairs of recorded neurons (*n*) are indicated in the panels. ns, *P* > 0.05; *, *P* < 0.05; **, *P* < 0.01; ***, *P* < 0.001; ****, *P* < 0.0001.

Layer 2/3 pyramidal cells in the V1 also receive monosynaptic excitation from the thalamus, local infragranular layer 5 and layer 6, and long-range inputs from other cortical areas (*42–44*). We expressed ReaChR in the dorsal lateral geniculate nucleus (dLGN) of the thalamus and photostimulated their axons in layer 2/3. Evoked EPSCs were similar between Kir2.1 and control neurons, and disynaptic IPSCs were decreased in Kir2.1 neurons at postnatal weeks 3 and 5 (Extended Data Fig. 5a–d). Thus, thalamus-evoked E/I ratios were increased in Kir2.1 neurons (Extended Data Fig. 5e). The reduction of disynaptic IPSCs is likely due to the decreased inhibition from parvalbumin-expressing interneurons (*37*), as thalamic inputs preferentially target this population of cortical interneurons (*43*), but the thalamic inputs do not contribute to the increased synaptic excitation in Kir2.1 neurons.

Layer 5 neurons provide a significant fraction of the total presynaptic inputs onto layer 2/3 pyramidal neurons (*41*, *42*, *44*, *45*), although these inputs are poorly studied. Strikingly, layer 5- evoked EPSCs were 2 and 3 times larger in Kir2.1 neurons than control neurons at postnatal weeks 3 and 5, respectively (Fig. 3k–m), whereas disynaptic IPSCs were similar between the two groups (Fig. 3l,n). As a result, layer 5-evoked E/I ratios were increased in Kir2.1 neurons (Fig. 3o). The larger increase of EPSC amplitude and E/I ratio at postnatal week 5 than postnatal week 3 (Fig. 3m,o) aligns with the increasing activity of Kir2.1 neurons during this period (Fig. 1).

Layer 5 excitatory neurons in the V1 consist of two subpopulations: intratelencephalic (IT) neurons, which project to other cortical areas, and extratelencephalic (ET) neurons, which additionally target subcortical structures (*47*, *48*). We found that both layer 5 IT and ET neurons- evoked EPSCs were increased in Kir2.1 neurons (Extended Data Fig. 6).

Layer 6, another infragranular layer, provides minor excitatory inputs onto layer 2/3 (*41*, *42*, *44*, *45*). Nevertheless, we found that layer 6-evoked EPSCs were also larger in Kir2.1 neurons at postnatal weeks 3 and 5, and disynaptic IPSCs were smaller at postnatal week 3 but comparable to the control levels at postnatal week 5 (Extended Data Fig. 5f–i). Consequently, layer 6-evoked E/I ratios were increased in Kir2.1 neurons at both ages (Extended Data Fig. 5j). Overexpression of a non-conducting Kir2.1 mutant (Extended Data Fig. 1i–m) did not alter layer 5 or layer 6- evoked excitation (Extended Data Fig. 7a–j), confirming the specific effect of reducing neuronal excitability by Kir2.1. These results identify infragranular layers, particularly layer 5, as a source of the increased excitation in Kir2.1 neurons and reveal an unexpected input-specific synaptic potentiation that homeostatically compensates for the reduction of neuron excitability *in vivo*.

### Input-selective increase of corticocortical long-range synaptic excitation

Layer 2/3 pyramidal cells in the V1 receive long-range excitatory inputs from other cortical areas (*42*). Do these inputs remain unaltered in Kir2.1 neurons, just like the long-range thalamic inputs, or show the same input-selective enhancement as the local inputs? We used the long-range callosal projections to address this question because they originate from the contralateral cortical layer 2/3 or layer 5 excitatory neurons (*49*). By photostimulating the callosal axons in the electroporated V1, we found that contralateral layer 2/3-evoked EPSCs and disynaptic IPSCs were slightly decreased in Kir2.1 neurons at postnatal week 3, resulting in similar E/I ratios as control neurons. At postnatal week 5, contralateral layer 2/3-evoked EPSCs, disynaptic IPSCs, and E/I ratios were similar between Kir2.1 and control neurons (Fig. 4a–e). In contrast, contralateral layer 5-evoked EPSCs were increased in Kir2.1 neurons at both postnatal weeks 3 and 5, without altering disynaptic IPSCs (Fig. 4f–i). Consequently, contralateral layer 5-evoked E/I ratios were increased in Kir2.1 neurons (Fig. 4j). Overexpression of a non-conducting Kir2.1 mutant did not alter contralateral layer 5-evoked excitation (Extended Data Fig. 7k–o). These results demonstrate that the presynaptic neuronal type determines the input specificity of homeostatic synaptic potentiation for both local and long-range synapses.

**Figure 4.**
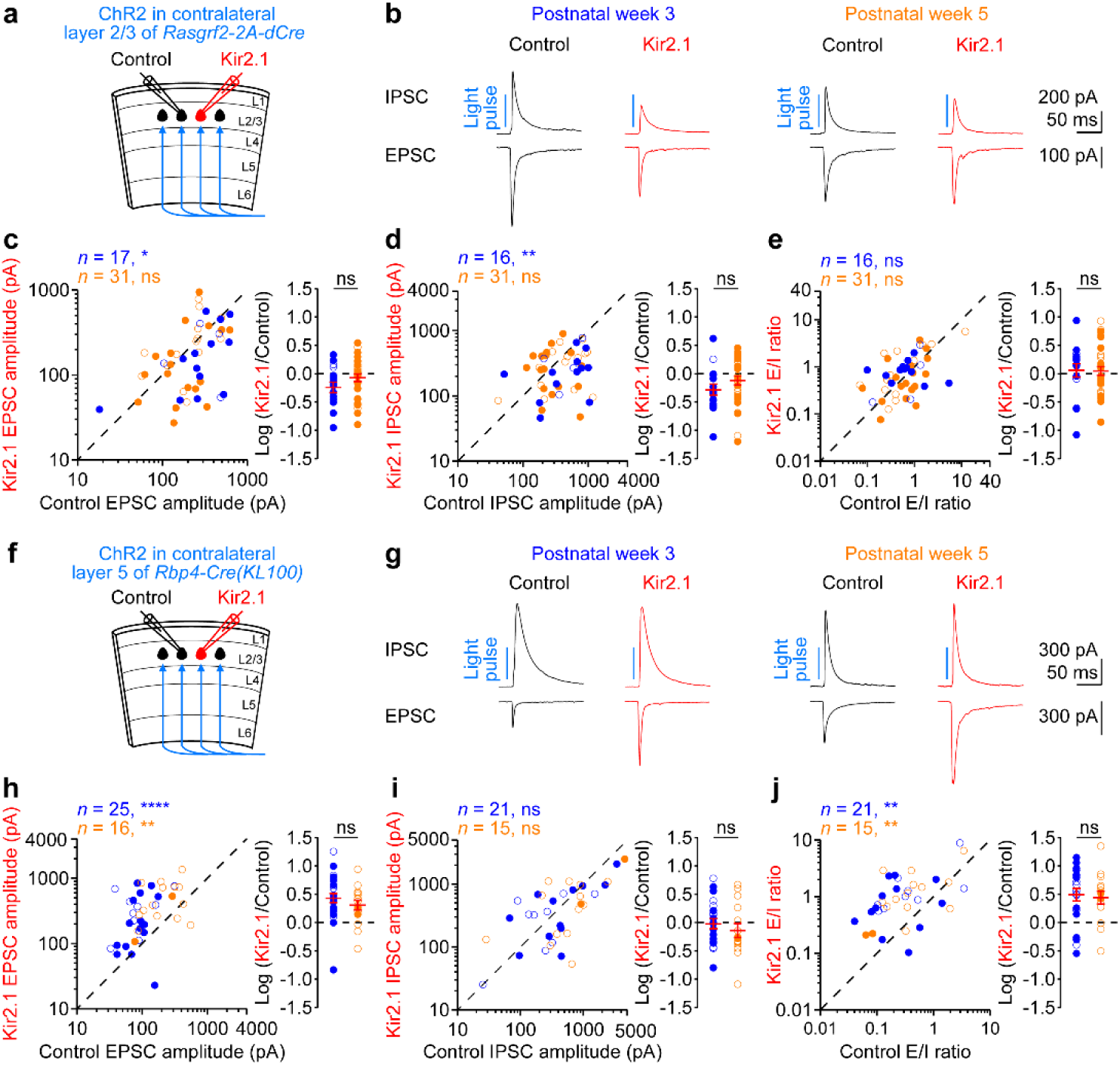
Excitation from contralateral layer 5 but not layer 2/3 is increased in Kir2.1 neurons. **(a)** Schematic of slice experiments in **b** with Kir2.1 in layer 2/3 pyramidal cells and ChR2 in the axons from contralateral layer 2/3 excitatory neurons. **(b)** Monosynaptic EPSCs and disynaptic IPSCs from a pair of simultaneously recorded control and Kir2.1 neurons in response to photostimulation. **(c)** Left, summary data of contralateral layer 2/3-evoked EPSC amplitudes in control and Kir2.1 neurons at postnatal weeks 3 (blue) and 5 (orange). Each filled (male) or open (female) symbol represents a pair of control and Kir2.1 neurons. Right, logarithm of the ratios between EPSC amplitudes from pairs of Kir2.1 and control neurons. Red, mean ± s.e.m. **(d**,**e)** As in **c**, but for IPSC amplitudes and E/I ratios. Note smaller EPSC and IPSC amplitudes in Kir2.1 neurons at postnatal week 3, similar EPSC and IPSC amplitudes between Kir2.1 and control neurons at postnatal week 5, and similar E/I ratios at both ages. **(f**‒**j)** As in **a**‒e, but for ChR2 in the axons from contralateral layer 5 excitatory neurons. Note similar IPSC amplitudes between Kir2.1 and control neurons, larger EPSC amplitudes, and larger E/I ratios in Kir2.1 neurons at both ages. The increase of EPSC amplitude and E/I ratio in Kir2.1 neurons are similar between the two ages. The numbers of pairs of recorded neurons (*n*) are indicated in the panels. Ns, *P* > 0.05; *, *P* < 0.05; **, *P* < 0.01; ****, *P* < 0.0001.

### Increased quantal amplitude at layer 5 excitatory inputs

The above results show that the potentiation of synaptic excitation is triggered by the reduction of neuronal excitability in postsynaptic Kir2.1 neurons, and its input-specificity is determined by the presynaptic neuronal type, but how is this input-selective synaptic potentiation expressed? Since both mEPSC amplitude and frequency are increased in Kir2.1 neurons, it is possible that the expression involves both presynaptic and postsynaptic mechanisms including quantal amplitude, presynaptic release probability, and the number of release sites or synapses. Thus, we used the local inputs from layer 5 excitatory neurons to determine these synaptic parameters at postnatal week 5.

To determine the quantal properties of these synapses, we first used an optogenetic method to isolate quantal EPSCs (qEPSCs) mediated by the glutamate release specifically from layer 5 excitatory neurons (*50*). We expressed ChR2 in layer 5 excitatory neurons and first recorded layer 5-evoked EPSCs in Kir2.1 and control neurons (Fig. 5a,b,d). We then recorded mEPSCs from the same neurons in the presence of voltage-gated sodium channel blocker, tetrodotoxin (TTX), but without voltage-gated potassium channel blockers. Under this condition, activation of ChR2 by a long ramping-down light stimulation did not evoke synchronous neurotransmitter release, but enhanced asynchronous synaptic vesicle exocytosis from layer 5 excitatory neurons, resulting in an increase in the frequency of mEPSCs (Fig. 5c). We mathematically subtracted the mEPSCs recorded during the baseline period (i.e., before light stimulation) from those recorded during light stimulation to obtain the frequency, average amplitude, and decay time constant of layer 5-evoked qEPSCs (Fig. 5c). The qEPSC amplitudes in Kir2.1 neurons were approximately 50% larger than those in control neurons (Fig. 5e), whereas the decay time constants were similar (Fig. 5h). This result indicates that the quantal releases from layer 5 neurons generate larger responses in Kir2.1 neurons, consistent with the increase in the amplitudes of total mEPSCs (Fig. 2j). We further measured the degree of blockade by a low-affinity competitive AMPA receptor antagonist, γ-DGG, on layer 5-evoked AMPA-EPSCs to differentiate between an increase in postsynaptic responsiveness and the amount of glutamate in presynaptic vesicles, as the more glutamate a vesicle releases, the lower the γ-DGG blockade is (*51*). We found that γ-DGG (1 mM) reduced EPSCs by 36% for both Kir2.1 and control neurons (Extended Data Fig. 8), supporting a postsynaptic mechanism for the increased quantal amplitudes in Kir2.1 neurons. However, the increased quantal amplitude was not sufficient to account for the increase in layer 5-evoked excitation (Fig. 5d‒f). Furthermore, the qEPSC frequencies in Kir2.1 neurons were more than twice those in control neurons (Fig. 5g). Together, these results suggest that in addition to an increase in quantal amplitude, other factors such as synapse number, presynaptic release probability, or both may contribute to the synaptic potentiation in Kir2.1 neurons.

**Figure 5.**
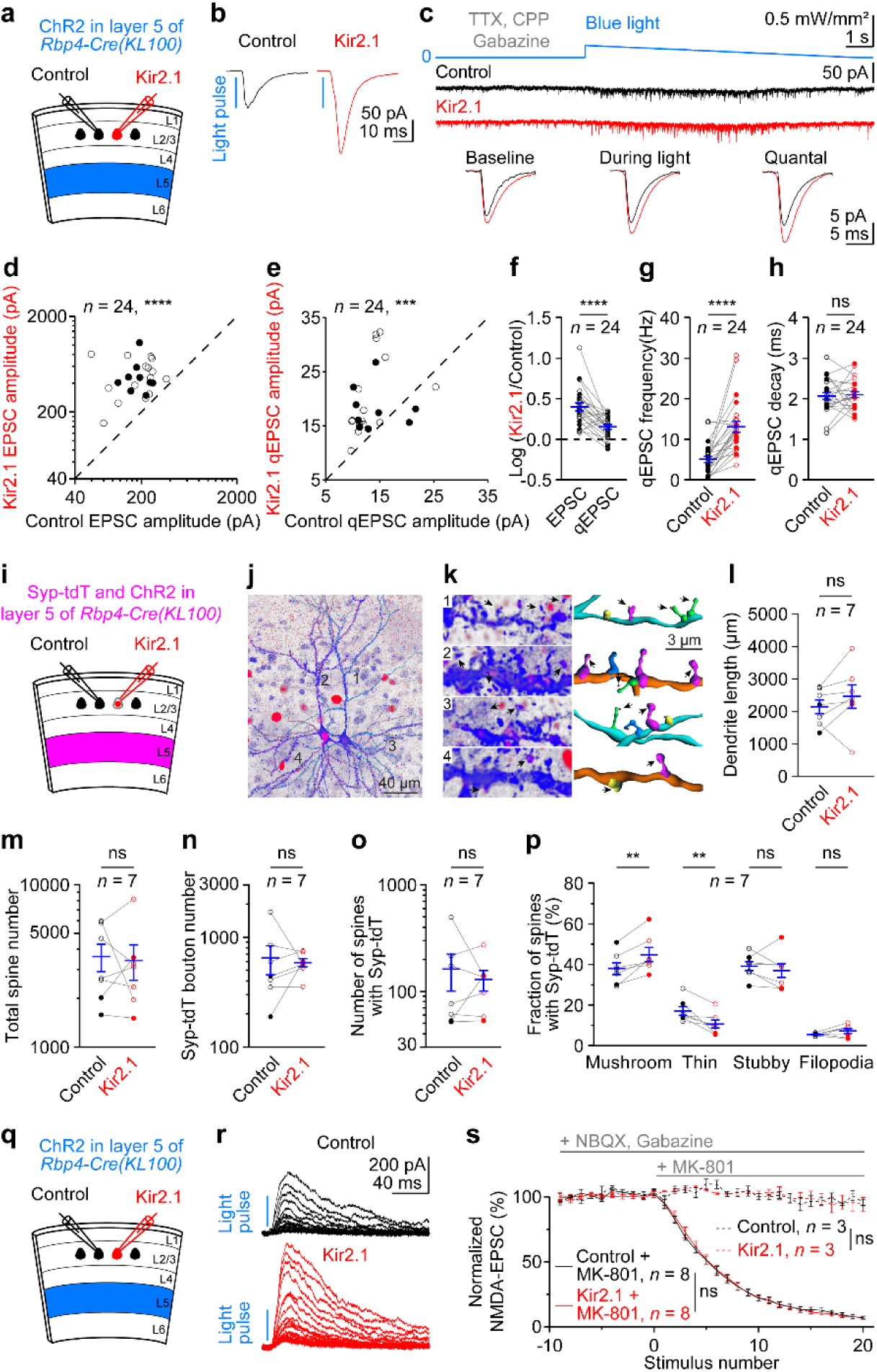
Postsynaptic mechanisms of increased excitation from layer 5 in Kir2.1 neurons. **(a)** Schematic of slice experiments in **b** and **c** with Kir2.1 in layer 2/3 pyramidal cells and ChR2 in layer 5 excitatory neurons at postnatal week 5. **(b)** EPSCs from a pair of simultaneously recorded control and Kir2.1 neurons in response to photostimulation. **(c)** AMPA receptor-mediated mEPSCs recorded in a pair of control and Kir2.1 neurons at baseline and during ramp-down light stimulation in the presence of TTX (voltage-gated sodium channel blocker), CPP (NMDA receptor antagonist) and Gabazine (GABAA receptor antagonist). The quantal EPSC (qEPSC) traces were computed based on the average mEPSC traces during the baseline period and light stimulation (see Methods). **(d**‒**h)** Summary data of layer 5-evoked EPSC amplitudes (**d**), qEPSC amplitudes (**e**), logarithm of the ratios between EPSC or qEPSC amplitudes (**f**), qEPSC frequencies (**g**), and decay time (**h**) from pairs of Kir2.1 and control neurons. Each or each pair of filled (male) or open (female) symbol(s) represents a pair of control and Kir2.1 neurons. Blue, mean ± s.e.m. Note smaller logarithm ratios of qEPSC amplitudes than those of EPSC amplitudes and higher qEPSC frequencies in Kir2.1 neurons. **(i)** Schematic of experiments in **j** and **k** with Kir2.1 in layer 2/3 pyramidal cells and Synaptophysin-tdTomato (Syp-tdT) and ChR2 in layer 5 excitatory neurons at postnatal week 5. **(j)** A pair of control and Kir2.1 (red, nuclear-localized dTomato) neurons filled with biocytin (blue). Reconstruction of the somata, dendrites, and spines from control and Kir2.1 neurons are shown in cyan and orange, respectively. **(k)** High-magnification images of 4 segments of dendrites from **j** (left) and their reconstructions (right). Segments 1 and 2 for apical dendrites and segments 3 and 4 for basal dendrites of control (cyan) and Kir2.1 (orange) neurons, respectively. Arrows, spines colocalized with Syp-tdT boutons (red, spine-Syp-tdT distance < 0.5 μm). Spines are classified as mushroom (magenta), thin (green), stubby (yellow), and filopodia (light blue). **(l**‒**p)** Summary data of dendrite lengths (**l**), total spine numbers (**m**), numbers of Syp-tdT boutons on dendrites and spines (**n**), numbers of spines colocalized with Syp-tdT (**o**), and fractions of different types of spines colocalized with Syp-tdT (**p**). Each pair of filled (male) or open (female) symbols represents a pair of control and Kir2.1 neurons. Blue, mean ± s.e.m. Note increased mushroom spines and decreased thin spines in Kir2.1 neurons. **(q)** Schematic of slice experiments in **r** with Kir2.1 in layer 2/3 pyramidal cells and ChR2 in layer 5 excitatory neurons at postnatal week 5. **(r)** NMDA-EPSCs from a pair of simultaneously recorded control and Kir2.1 neurons in response to repetitive photostimulations in the presence of NBQX (AMPA receptor antagonist), Gabazine, and MK-801 (open NMDA receptor blocker). **(s)** Summary data of NMDA-EPSC amplitudes without MK-801 (dashed lines) or with MK-801 following the baseline period (solid lines). Data are mean ± s.e.m. Note similar NMDA-EPSC blockade rates between control and Kir2.1 neurons. The numbers of pairs of recorded neurons (*n*) are indicated in the panels. ns, *P* > 0.05; **, *P* < 0.01; ****, *P* < 0.0001.

### Unaltered morphological synapse number and presynaptic release probability at layer 5 excitatory inputs

To determine the number of layer 5 excitatory synapses impinged onto Kir2.1 and control neurons, we expressed ChR2 and synaptic vesicle protein Synaptophysin fused with tdTomato (Syp-tdT) in layer 5 excitatory neurons for photostimulation and labeling presynaptic boutons, respectively. We recorded Kir2.1 and control neurons to confirm the increase of EPSCs in Kir2.1 neurons in these experiments (data not shown) and filled the recorded neurons with biocytin for visualizing their morphology. We reconstructed the somata, dendrites, and spines and identified Syp-tdT-labelled presynaptic boutons onto the dendrites and spines. The layer 5 excitatory synapses impinged onto Kir2.1 and control neurons were identified by the colocalization of Syp-tdT-labelled presynaptic boutons with postsynaptic spines (Fig. 5j,k). The total dendrite length, spine number and density, Syp-tdT-labelled bouton number and density, and layer 5 excitatory synapse number and density were unaltered in Kir2.1 neurons (Fig. 5l–o; Extended Data Fig. 9), indicating that the number of morphological synapses is an unlikely contributor to the synaptic potentiation in Kir2.1 neurons. Interestingly, among these synapses from layer 5 excitatory neurons we observed more mushroom-like spines and fewer thin spines in Kir2.1 neurons than control neurons (Fig. 5p), indicating a conversion from immature to mature synapses in Kir2.1 neurons.

To determine the presynaptic release probability, we repetitively photostimulated layer 5 excitatory neurons and recorded N-methyl-D-aspartic acid (NMDA) receptor-mediated EPSCs (NMDA-EPSCs) from Kir2.1 and control neurons in the presence of a use-dependent, open NMDA receptor blocker, MK-801. Thus, the rate of NMDA receptor blockade is proportional to presynaptic release probability (*52*, *53*). The blockade rates of NMDA-EPSCs were similar between Kir2.1 and control neurons (Fig. 5q–s), indicating that the presynaptic release probability of layer 5 excitatory inputs impinged onto Kir2.1 neurons is unaltered.

### Insertion of GluA2-containing AMPA receptors unsilences silent synapses at layer 5 excitatory inputs

The experiments above did not identify changes in morphological synapse number and presynaptic release probability, yet the frequencies of layer 5-evoked qEPSCs were elevated in Kir2.1 neurons. What mechanism would explain this seemingly discrepancy and account for the larger increase of layer 5-evoked EPSCs beyond the increased qEPSC amplitudes? When measuring presynaptic release probability, we noticed that layer 5-evoked NMDA-EPSCs were also increased in Kir2.1 neurons by about 50%, which is smaller than the increase of AMPA-EPSCs (Fig. 5r), indicating a postsynaptic mechanism that affects AMPA and NMDA receptors differently. Considering the amplitudes of AMPA qEPSCs increased by only 50%, while their frequencies more than doubled, one possibility is that a larger fraction of the excitatory synapses from layer 5 onto Kir2.1 neurons contain AMPA receptors. In other words, a greater number of silent synapses, which initially express only NMDA receptors, are unsilenced by AMPA receptor insertion during development in Kir2.1 neurons, a process known to associate with an increase in the AMPA-EPSC and NMDA-EPSC ratio (A/N ratio). To test this hypothesis, we measured layer 5-evoked AMPA-EPSCs and NMDA-EPSCs from the same cells and found that they were about 3 and 1.5 times larger in Kir2.1 neurons than control neurons, respectively (Fig. 6a–d), leading to a 2-fold increase of A/N ratio in Kir2.1 neurons (Fig. 6e). A larger fraction of mature spines in Kir2.1 neurons (Fig. 5p) is also consistent with the unsilencing of silent synapses, as mature spines contain AMPA receptors, whereas silent synapses are often immature (*54*). Thus, the number of AMPA receptor-containing functional synapses is increased in Kir2.1 neurons despite unaltered morphological synapse number.

**Figure 6.**
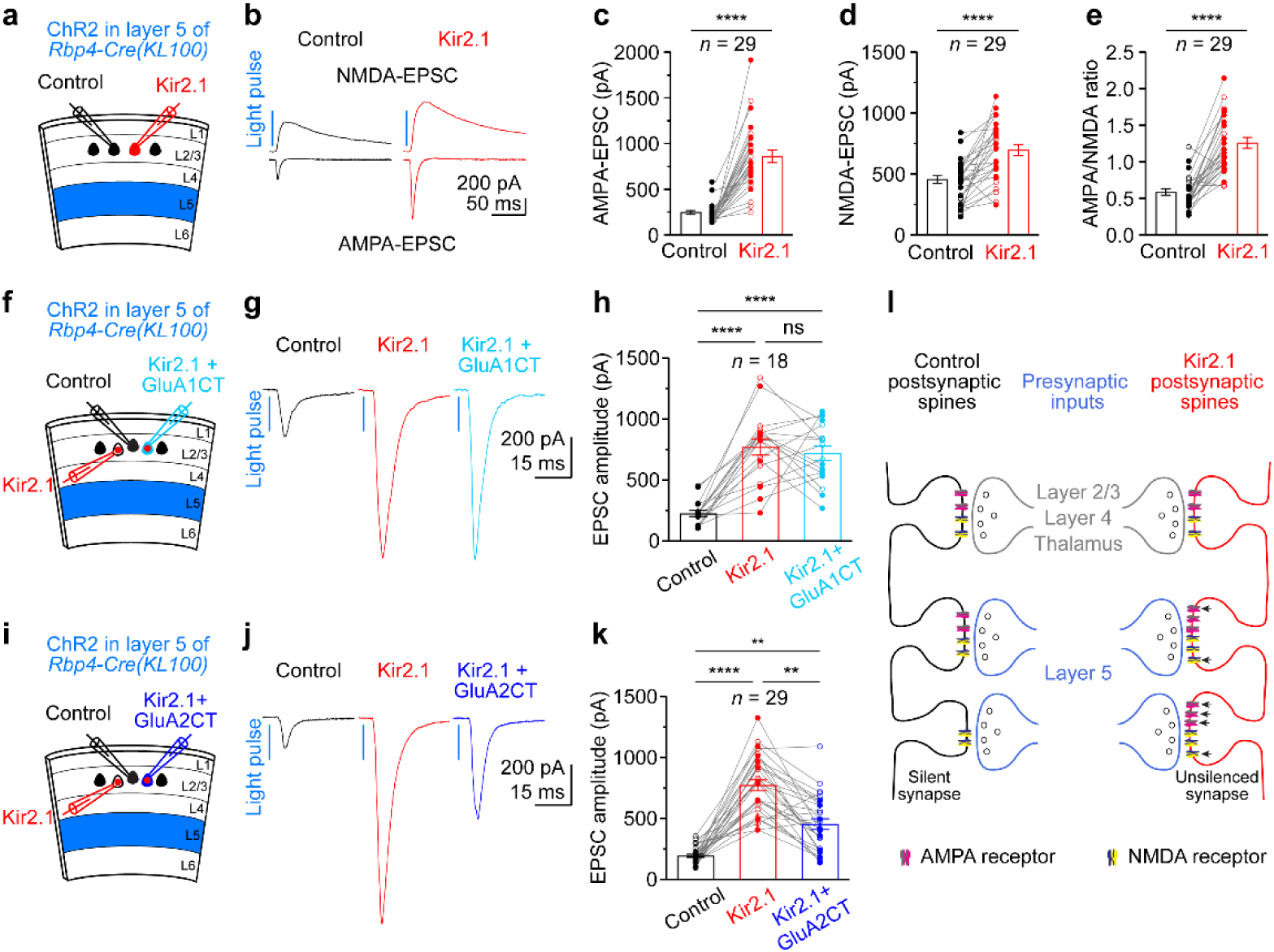
Unsilencing of silent synapses contributes to increased excitation from layer 5 in Kir2.1 neurons. **(a)** Schematic of slice experiments in **b** with Kir2.1 in layer 2/3 pyramidal cells and ChR2 in layer 5 excitatory neurons at postnatal week 5. **(b)** AMPA- and NMDA-EPSCs from a pair of simultaneously recorded control and Kir2.1 neurons in response to photostimulation. **(c**‒**e)** Summary data of AMPA-EPSC amplitudes (**c**), NMDA-EPSC amplitudes (**d**), and AMPA/NMDA ratios (**e**). Each pair of filled (male) or open (female) symbols represents a pair of control and Kir2.1 neurons. Bars, mean ± s.e.m. Note that not only AMPA- and NMDA-EPSC amplitudes are increased, AMPA/NMDA ratios are also enhanced in Kir2.1 neurons. **(f)** Schematic of slice experiments in **g** with Kir2.1 alone or co-expressed with GluA1CT in layer 2/3 pyramidal cells and ChR2 in layer 5 excitatory neurons at postnatal week 5. **(g)** EPSCs from a triplet of simultaneously recorded control, Kir2.1, and Kir2.1/GluA1CT neurons in response to photostimulation. **(h)** Summary data of EPSC amplitudes. Each triplet of filled (male) or open (female) symbols represents a triplet of control, Kir2.1, and Kir2.1/GluA1CT neurons. Bars, mean ± s.e.m. Note, GluA1CT does not alter the effect of Kir2.1. **(i**‒**k)** As in **f**‒**h**, but for GluA2CT. Note, GluA2CT reduces the effect of Kir2.1. **(l)** Model of postsynaptic mechanisms for input-selective homeostatic synaptic plasticity in Kir2.1 neurons that involve increasing postsynaptic NMDA and AMPA receptors and inserting AMPA receptors, at least part of which contain GluA2 subunits, into silent synapses specifically at the layer 5 excitatory inputs. Arrows indicate incorporated AMPA and NMDA receptors at previously active and silent synapses. The numbers of pairs or triplets of recorded neurons (*n*) are indicated in the panels. ns, *P* > 0.05; **, *P* < 0.01; ****, *P* < 0.0001.

To pinpoint which AMPA receptors are involved in the unsilencing of silent synapses, we overexpressed a C-terminal peptide from AMPA receptor subunit GluA1 (GluA1CT) or GluA2 (GluA2CT) to disrupt their insertion into synapses in Kir2.1 or control neurons (*55*). Co-expression of Kir2.1 with either GluA1CT or GluA2CT resulted in the same degree of reduction in neuronal excitability as Kir2.1 alone (Extended Data Fig. 10a–d). GluA1CT overexpression did not affect layer 5-evoked EPSCs in control (Extended Data Fig. 10e–g) or Kir2.1 neurons (Fig. 6f–h), indicating a minimal role of GluA1 in the potentiation of these synapses in Kir2.1 neurons (Extended Data Fig. 10h). In contrast, GluA2CT overexpression significantly reduced layer 5-evoked EPSCs in both control (Extended Data Fig. 10i–k) and Kir2.1 neurons (Fig. 6i– k). Importantly, GluA2CT-induced reduction of EPSCs was greater in Kir2.1 neurons than control neurons (Extended Data Fig. 10l), showing a crucial role of GluA2 subunit, but not GluA1, in the homeostatic potentiation of layer 5 excitatory inputs in Kir2.1 neurons. Thus, the input-selective potentiation of excitation in Kir2.1 neurons is mediated by postsynaptic mechanisms that not only increase both synaptic AMPA and NMDA receptors but also activate silent synapses through the insertion of GluA2-containing AMPA receptors (Fig. 6l), compensating for the reduced neuronal excitability during development to achieve the targeted setpoint activity levels.

## Discussion

Homeostatic plasticity has largely been conceptualized as a stabilizing mechanism that maintains neuronal activity around a pre-existing setpoint level once a circuit has formed (*5–8*). In contrast, our findings show that even when silenced by Kir2.1 overexpression from inception, developing cortical neurons “know” their activity setpoints and initiate a homeostatic mechanism to drive their activity toward the target levels despite persistent perturbations. This suggests that the activity “homeostat” is already operational from birth before the setpoint is first reached, and the genetic program defining a cortical neuron includes a target activity level that the developing neuron actively pursues when transitioning from the silent state at birth to an active circuit element. Thus, neuronal activity setpoints are encoded as fundamental properties of neuronal identity rather than mere outcomes of network equilibrium.

In response to perturbation of their neuronal excitability *in vivo*, cortical neurons execute a cell-autonomous homeostatic program that is strikingly selective for specific synaptic inputs. In contrast to the classical homeostatic multiplicative synaptic scaling that adjusts all inputs uniformly to preserve their relative weights (*22*), previous studies reported examples of synapse-specific homeostatic plasticity that redistribute synaptic weights across excitatory inputs (*32*, *56*, *57*). Kir2.1 neurons differentially weight specific excitatory afferent pathways by potentiating infragranular inputs, particularly those from layer 5, while maintaining major canonical inputs from layer 4 and layer 2/3 unchanged. Such selectivity of homeostatic mechanism may reflect differences in developmental timing, synapse maturity, or molecular compatibility between presynaptic and postsynaptic partners. Layer 5 excitatory neurons are major cortical output neurons that broadcast signals across cortical and subcortical targets (*58*, *59*). The preferential strengthening of their synapses onto layer 2/3 neurons suggests a critical role of this feedback pathway in regulating the activity of superficial layers (*60–62*). We speculate that this organization may provide a strategic advantage for maintaining network stability. When superficial circuits are weakened, deeper-layer inputs are upregulated to restore activity without perturbing the relative structure of canonical layer 4 feedforward and layer 2/3 recurrent pathways and compromising primary sensory information (*63*, *64*). Thus, our results suggest that homeostatic regulation can operate not only at the level of individual synapses but also through selective engagement of specific circuit motifs by distinguishing between presynaptic partners based on their identities. The observation that both local and long-range layer 5 inputs are preferentially strengthened further supports that presynaptic identity, rather than anatomical origin *per se*, is a key determinant of this homeostatic response.

Mechanistically, the potentiation of layer 5 excitatory inputs is solely mediated by postsynaptic modifications because none of the synapse number, presynaptic release probability, or glutamate in vesicles is altered, and the increased AMPA quantal amplitude and AMPA/NMDA ratio can fully account for the potentiation in Kir2.1 neurons. The increase in AMPA/NMDA ratio and the shift toward mature spines support a novel model in which homeostatic synaptic plasticity engages AMPA-silent synapses to become functionally active. As unsilencing silent synapse is a well-established mechanism in Hebbian long-term potentiation (*65*), our findings suggest an intriguing convergence of these two distinct plasticity mechanisms at the synaptic level. Previous studies found that activity blockade induced an upregulation of AMPA receptor subunit GluA1, GluA2, or both (*20*, *23*, *66–69*). Here we identify GluA2-containing AMPA receptor insertion as a key molecular effector of input-selective homeostatic plasticity, as disrupting GluA2 trafficking impairs the potentiation in Kir2.1 neurons, whereas interfering with GluA1 has little effect, although we cannot rule out the possibility that GluA1 may be transiently inserted during the development of this input-specific homeostatic compensation (*70*).

Apart from enhanced excitation, the disynaptic inhibition recruited by layer 4, layer 6, or thalamic inputs is reduced in Kir2.1 neurons. This is likely due to a reduction of parvalbumin-expressing interneurons-mediated inhibition, as it is selectively affected in Kir2.1 neurons (*37*). Together, increased excitation from layers 5 and 6 and decreased inhibition recruited by layer 4, layer 6, and thalamic inputs compensate for reduced intrinsic excitability in Kir2.1 neurons, allowing them to reach their setpoint level of activity.

The ability of cortical neurons to reach appropriate activity levels despite persistent perturbations highlights a robust developmental program that safeguards brain function. This robustness is likely critical given the variability inherent in genetic expression, developmental processes, and sensory experience. Our findings support a model in which neuronal identity includes a target activity setpoint that is actively enforced through homeostatic monitoring and selective synaptic adjustment, linking cellular identity to circuit-level stability during cortical development.

Disruptions in such mechanisms could contribute to circuit imbalances observed in neurodevelopmental disorders. Thus, our study provides a new conceptual and mechanistic foundation for understanding neurodevelopmental robustness and also raises several important questions for future investigation. First, the molecular mechanisms that enable developing neurons to sense deviations from their targeted activity setpoints and selectively potentiate specific excitatory inputs are unknown. Second, we reduced the intrinsic excitability of developing neurons from their birth. It remains to be determined when perturbations occur at different developmental stages, if and how neurons achieve activity setpoints. Third, whether an excitatory input-selective homeostatic mechanism is used by developing neurons to counteract enhanced excitability requires further investigation. Finally, although our experiments establish a cell-autonomous mechanism, how such processes interact with network-level homeostatic regulation remains to be determined.

## Acknowledgements

We thank Massimo Scanziani for support in whose laboratory preliminary experiments were performed, Hongmei Chen for assistance in plasmid preparation, Jessica Messier for help with AAV injections, Charles Gerfen, Nuo Li, and Hui-Chen Lu for sharing mouse lines, and Baohua Liu, Kevin Jiang, Massimo Scanziani, and Kyunghee Kim for comments on earlier versions of the manuscript. We acknowledge the use of ChatGPT and Gemini to assist in editing the language of the manuscript. The authors reviewed and edited the content for accuracy and take full responsibility for the final text. This work was supported in part by the National Institute of Mental Health (R01MH117089 to M.X. and F30MH118804 to C.M.L.), the Eunice Kennedy Shriver National Institute of Child Health and Human Development (P50HD103555 to Baylor College of Medicine Intellectual and Developmental Disabilities Research Center, Neurovisualization Core), and McKnight Foundation (a McKnight Scholar Award to M.X.). M.X. is a Caroline DeLuca Scholar.

## Methods

### Mice

Mice were housed in an Association for Assessment and Accreditation of Laboratory Animal Care International-certified animal facility on a 14-hour/10-hour light/dark cycle and provided with regular mouse chow and water ad libitum. All procedures to maintain and use mice were performed in strict accordance with the recommendations in the Guide for the Care and Use of Laboratory Animals of the National Institutes of Health and were approved by the Institutional Animal Care and Use Committee at Baylor College of Medicine (protocol AN- 6544). Experiments were performed during the light cycle. Hemizygous transgenic and heterozygous knock-in mice of both sexes were used in experiments. Female ICR mice were purchased from Baylor College of Medicine Center for Comparative Medicine or Charles River Laboratories. C57BL/6J, *Scnn1a-Cre(Tg3)*, and *Rasgrf2-2A-dCre* mice were obtained from the Jackson Laboratory (stock numbers 000664, 009613 and 022864, respectively), *Ntsr1- Cre(GN220)* and *Rbp4-Cre(KL100)* mice from the Mutant Mouse Resource and Research Center (stock numbers 017266-UCD and 031125-UCD, respectively), *Tlx3-Cre(PL56)* and *Chrna2- Cre(OE25)* mice from Dr. Charles Gerfen at the National Institute of Mental Health and Dr. Nuo Li at Baylor College of Medicine, and *Rora-ires-Cre* mice from Dr. Dennis O’Leary at Salk Institute for Biological Research and Dr. Hui-Chen Lu at Indiana University Bloomington. To induce recombination in *Rasgrf2-2A-dCre* mice, antibiotic trimethoprim (TMP) was intraperitoneally injected at 100–150 mg/kg body weight per day (*71*) on P13 and P14 (for experiments in postnatal week 3) or P17 and P18 (for experiments in postnatal week 5).

### DNA constructs

Plasmids pCAG-Kir2.1-T2A-tdTomato (Addgene 60598) expressing Kir2.1 E224G Y242F (referred to as Kir2.1) and pCAG-Kir2.1Mut-T2A-tdTomato (Addgene 60644) expressing a non-conducting Kir2.1 (Kir2.1 E224G Y242F G144A Y145A G146A, referred to as Kir2.1Mut) were described previously (*37*). pCAG-Kir2.1-P2A-3×NLS-dTomato (Addgene 245282) was generated by replacing the T2A-tdTomato in pCAG-Kir2.1-T2A-tdTomato with P2A-3×NLS-dTomato. pCAG-Kir2.1-T2A-EGFP (Addgene 245283) was generated by inserting the Kir2.1-T2A from pCAG-Kir2.1-T2A-tdTomato into pCAG-EGFP (Addgene 11150). pAAV- EF1α-DIO-Kir2.1-T2A-tdTomato (Addgene 245284) was generated by replacing the hChR2(H134R)-mCherry in pAAV-EF1α-DIO-hChR2(H134R)-mCherry (Addgene 20297) with Kir2.1-T2A-tdTomato. To generate pCAG-loxP-hChR2(H134R)-EYFP-loxP (Addgene 245285), a loxP-MCS (multiple cloning sites)-loxP cassette was first synthesized *de novo* and cloned into plasmid pIDTSMART by IDT to generate pIDTSMART-loxP-MCS-loxP. The hChR2(H134R)-EYFP from pAAV-EF1α-DIO-hChR2(H134R)-EYFP (Addgene 20298) was inserted into pIDTSMART-loxP-MCS-loxP to generate pIDTSMART-loxP-hChR2(H134R)-EYFP-loxP. The EGFP in pCAG-EGFP was then replaced by loxP-hChR2(H134R)-EYFP-loxP to generate pCAG-loxP-hChR2(H134R)-EYFP-loxP. Plasmid pAAV-hSyn-DIO-Synaptophysin-tdTomato (Addgene 245286) was generated by replacing the EF1α and oChIEF(E163A/T199C)-P2A- dTomato in pAAV-EF1α-DIO-oChIEF(E163A/T199C)-P2A-dTomato (Addgene 51094) with hSyn from pAAV-hSyn-Con/Fon-EYFP-WPRE (Addgene 55650) and Synaptophysin-tdTomato from pROSA26-CAG-LSL-Synaptophysin-tdTomato (Addgene 34881), respectively. Plasmid pAAV-EF1α-FRT-FLEX-EGFP (Addgene 190770) was generated by replacing the mNaChBac- T2A-tdTomato of pAAV-EF1α-FRT-FLEX-mNaChBac-T2A-tdTomato (Addgene 60658) with the EGFP from pCAG-EGFP. The C-terminal peptides of mouse AMPA receptor subunit GluA1 and GluA2 (GluA1CT: EFCYKSRSESKRMKGFCLIPQQSINEAIRTSTLPRNSGAGASGGSGSGENGRVVSQDFPKSMQSIPCMSHSSGMPLGATGL, GluA2CT: EFCYKSRAEAKRMKVAKNAQNINPSSSQNSQNFATYKEGYNVYGIESVKI) (*72*) with a linker GSGSGSGSGS were generated by PCR from the mouse genomic DNA and inserted into pAAV-EF1α-FRT-FLEX-EGFP to create pAAV-EF1α-FRT-FLEX-EGFP-linker-GluA1CT (Addgene 245287) and pAAV-EF1α-FRT-FLEX-EGFP-linker-GluA2CT (Addgene 245288), respectively.

### *In utero* electroporation

Female ICR mice at the age of 6–8 weeks were crossed with male C57BL/6J, *Scnn1a-Cre(Tg3)*, *Rasgrf2-2A-dCre*, *Ntsr1-Cre(GN220)*, *Rbp4-Cre(KL100)*, *Tlx3- Cre(PL56)*, *Chrna2-Cre(OE25)*, or *RORa-ires-Cre* mice to obtain timed pregnancies. On embryonic days 15 or 15.5, the female mice underwent the *in utero* electroporation procedure for transfecting a subset of layer 2/3 pyramidal neurons in the primary visual cortex (V1) as described previously (*37*, *73*). Briefly, a DNA solution (1.5 μl) was injected into one lateral ventricle using a beveled glass micropipette (tip size: 80-μm outer diameter, 50-μm inner diameter). Five electronic pulses (voltage: 39 V, duration: 50 ms) were delivered at 1 Hz with a Tweezertrode (5-mm diameter) and a square-wave pulse generator (BTX Gemini X2 Generator). Plasmids pCAG-Kir2.1-T2A-tdTomato, pCAG-Kir2.1Mut-T2A-tdTomato, pCAG-Kir2.1-P2A- 3×NLS-dTomato, and pCAG-Kir2.1-T2A-EGFP were used at the final concentrations of 1.5–3.0 μg/μl. Plasmids pAAV-EF1α-DIO-Kir2.1-T2A-tdTomato, pCAG-loxP-hChR2(H134R)-EYFP-loxP, and pCAG-EGFP were mixed at the final concentrations of 2.5 μg/μl, 2.0 μg/μl, and 0.2 μg/μl, respectively. Plasmid pAAV-EF1α-FRT-FLEX-EGFP-linker-GluA1CT or pAAV-EF1α- FRT-FLEX-EGFP-linker-GluA2CT was mixed with pCAG-Kir2.1-P2A-3×NLS-dTomato at the final concentrations of 2.0 μg/μl and 2.4 μg/μl, respectively. Fast Green (Sigma-Aldrich, 0.01% final concentration) was added to the DNA solution. After birth, transfected pups were identified by the transcranial fluorescence of tdTomato, dTomato, or EGFP using a stereomicroscope (Leica MZ10F).

### AAV production and injection

Recombinant AAV vectors AAV9-EF1α-DIO-hChR2(H134R)- EYFP (AV-9-20298P, 7.30 × 10^12^ genome copies/ml), AAV1-EF1α-DIO-hChR2(H134R)-EYFP (AV-1-20298P, 6.82 × 10^12^ genome copies/ml), and AAV9-hSyn-HI-eGFP-Cre (AV-9-PV1848, 1.67 × 10^12^ genome copies/ml) were obtained from the Penn Vector Core. AAV9-hSyn-Flpo (3.93 × 10^11^ genome copies/ml), AAV9-EF1α-DIO-ReaChR-P2A-dTomato (7.00 × 10^12^ genome copies/ml), and AAV9-EF1α-DIO-hChR2(H134R)-P2A-EYFP (1.50 × 10^13^ genome copies/ml) were produced and described previously (*37*, *50*, *73*). AAV9-hSyn-DIO-Synaptophysin- tdTomato (8.26 × 10^11^ genome copies/ml) was produced by Baylor College of Medicine Gene Vector Core.

AAV vectors were injected into the V1 at P0–2 as described previously (*37*, *73*) using an UltraMicroPump injector (UMP3, World Precision Instruments). Briefly, a beveled glass micropipette (tip size: 50-μm outer diameter, 30-μm inner diameter) was used to penetrate the scalp and skull and inject AAV at various depths (targeting cortical layer 2/3: 450, 350, 250 μm below the scalp; layer 4: 550, 450, 350 μm below the scalp; layer 5 and layer 6: 650, 550, 450 μm below the scalp; thalamus: 2200, 2000, 1800 μm below the scalp) at one location (V1: 1.6 mm lateral and 0.3 mm caudal from the lambda; thalamus: 1.5 mm lateral and 0.5 mm caudal from the lambda). A total of 150 nl of virus solution with the final concentrations described above was injected over 60 seconds. After injection, the micropipette was kept in the parenchyma at the final (shallowest) depth for 30 s before being slowly withdrawn.

To express ChR2 in specific cortical excitatory neurons, AAV9-EF1α-DIO-hChR2(H134R)- EYFP, AAV1-EF1α-DIO-hChR2(H134R)-EYFP, or AAV9-EF1α-DIO-hChR2(H134R)-P2A-EYFP was injected into the V1 of different Cre mouse lines: *Scnn1a-Cre(Tg3)* for layer 4, *Rbp4- Cre(KL100)* for ipsilateral or contralateral layer5, *Tlx3-Cre(PL56)* for layer 5 IT, *Chrna2- Cre(OE25)* for layer 5 ET; *Ntsr1-Cre(GN220)* for layer 6, and *Rasgrf2-2A-dCre* for contralateral layer 2/3. AAV9-EF1α-DIO-ReaChR-P2A-dTomato was injected into the thalamus of *RORa- ires-Cre* to express ReaChR in the dLGN. For local layer 2/3 excitatory neurons, two plasmids pAAV-EF1α-DIO-Kir2.1-T2A-tdTomato and pCAG-loxP-hChR2(H134R)-EYFP-loxP were transfected into a subset of layer 2/3 pyramidal cells in the V1 via *in utero* electroporation. AAV9-hSyn-HI-eGFP-Cre was injected into the V1 at P0‒1 to induce expression of Kir2.1 and deletion of ChR2 in approximately one third of the electroporated cells, so that the photocurrent of ChR2 would not interfere with the measurements of synaptic currents from Kir2.1 neurons while the rest of the electroporated cells without Cre expressed ChR2 for photostimulation of layer 2/3 inputs.

### *In vivo* physiology

Mice between postnatal days 10 and 38 were anesthetized with an intraperitoneal injection of chlorprothixene (5 mg/kg) followed by urethane (1.2 g/kg). Oxygen was given at a flow rate of 1 l/min during the experiments, with isoflurane (<0.5%) supplemented if necessary. Body temperature was maintained at 37°C using a feedback-based DC temperature control system. Dexamethasone sodium phosphate (2 mg/kg) and Lactated Ringer’s Injection (3 ml/kg) were administered subcutaneously. Whiskers and eyelashes were trimmed, and a thin layer of silicone oil (kinematic viscosity 30,000 cSt, Sigma-Aldrich) was applied to the corneas to prevent drying. The scalp and periosteum were removed, and Vetbond tissue adhesive (3M) was applied to stabilize all sutures. A custom-made recording chamber with a 3-mm diameter hole in the center was attached to the skull over the V1 with Vetbond tissue adhesive and dental cement (Ortho-Jet BCA, Lang Dental). The recording chamber was then secured on a custom-made holder. A craniotomy (1.5–2 mm diameter, centered at 2.5 mm lateral to midline and 1 mm rostral to lambda suture) was performed with a 0.3-mm diameter round bur on a high-speed rotary micromotor. The dura was left intact, and the craniotomy was covered by a thin layer of 1.5% type III-A agarose in HEPES-ACSF containing 142 mM NaCl, 5 mM KCl, 10 HEPES, 1.3 mM MgCl_2_, 3.1 mM CaCl_2_, and 10 mM D-glucose (310 mosmol, pH 7.4). HEPES-ACSF was added to the recording chamber.

Targeted loose-patch recordings were performed under the guidance of a two-photon laser scanning microscope. Two-photon imaging was conducted with a water immersion objective (40×, 0.8 numerical aperture, Olympus) on a Moveable Objective Microscope (Sutter Instrument) coupled with a Ti:Sapphire laser (Chameleon Ultra II, Coherent) under the control of ScanImage 3.6 (Janelia Farm Research Campus, HHMI). The laser wavelength was tuned to 950 nm (laser power after the objective: 25–50 mW) for two-photon excitation of tdTomato and Alexa Fluor 488.

An Axon Multiclamp 700B amplifier was used for extracellular recording of spikes. A patch pipette containing HEPES-ACSF and 10–20 μM Alexa Fluor 488 hydrazide (Life Technology) was advanced along its axis towards neurons located between 150 and 250 μm below the dura with minimal lateral movements. A small positive pressure was applied to the patch pipette to avoid clogging of the tip and inject a small amount of fluorescent dye to stain the extracellular space. Non-fluorescent neurons were visualized as negative images. The pipette resistance was constantly monitored in voltage-clamp mode. The concurrence of the pipette tip contacting the neuron and an increase in pipette resistance indicated successful targeting, which was further confirmed post hoc. Upon the release of positive pressure, a small negative pressure was applied to form a loose seal (10–30 MΩ). The amplifier was then switched to the current-clamp mode with zero current injection to record voltage. Data were low-pass filtered at 10 kHz and acquired at 32 kHz with a NI-DAQ board (NI PCIe-6259, National Instruments) under the control of a custom-written program running in MATLAB (MathWorks). Within a local region (< 50 μm), neighboring tdTomato positive (Kir2.1) and tdTomato negative (control) neurons were sequentially recorded in a random order. The correct targeting of tdTomato positive neurons was confirmed at the end of the recording either by filling the neuron with the fluorescent dye contained in the pipette via break-in or by the presence of neuronal fluorescence in the recording pipette due to the negative pressure.

Visual stimuli were generated in MATLAB with Psychophysics Toolbox and displayed on a gamma-corrected liquid-crystal display monitor (30 cm × 47.5 cm, 60 Hz refresh rate, mean luminance 50 cd/m^2^). The monitor was approximately centered at the retinotopic location corresponding to the V1 recording site by monitoring single-unit or multi-unit activity in response to a moving bar on the screen. During recordings, full-field sinusoidal drifting gratings (temporal frequency 0.04 cycles/degree, 100% contrast) were presented randomly at 12 different directions from 0° to 330° for 1.5 s, preceded and followed by the presentation of a grey screen for 2 s and 1.5 s, respectively. The complete set of stimuli was repeated 8–16 times.

Data were analyzed offline using a custom-written program in MATLAB. Voltage signals were high-pass filtered (125 Hz). Spikes were first detected as events exceeding five times the standard deviation of the noise, and then visually verified. Spontaneous spike rate was calculated as the average spike rate during the 2-s time window before the presentation of a visual stimulus. Evoked spike rate was calculated as the average spike rate during the 1.5-s time window of visual stimulation. Overall spike rate was calculated as the average spike rate during the entire recording period.

### Brain slice electrophysiology

Mice were anesthetized by an intraperitoneal injection of ketamine and xylazine mix at 3 times the euthanasia dose (240 and 48 mg/kg body weight, respectively) and transcardially perfused with cold (0–4°C) slice cutting solution containing 80 mM NaCl, 2.5 mM KCl, 1.3 mM NaH_2_PO_4_, 26 mM NaHCO_3_, 4 mM MgCl_2_, 0.5 mM CaCl_2_, 20 mM D-glucose, 75 mM sucrose, and 0.5 mM sodium ascorbate (315–320 mosmol, pH 7.4, saturated with 95% O_2_/ 5% CO_2_). Brains were removed and sectioned in the cold cutting solution using a microtome with vibrating blade (Leica VT1200S) to obtain 300-μm coronal slices. Slices were incubated in a custom-made interface holding chamber saturated with 95% O_2_/ 5% CO_2_ at 34°C for 30 minutes and then at room temperature for 20 minutes to 8 hours until transferred to the recording chamber.

Recordings were performed on submerged slices in artificial cerebrospinal fluid (ACSF) containing 119 mM NaCl, 2.5 mM KCl, 1.3 mM NaH_2_PO_4_, 26 mM NaHCO_3_, 1.3 mM MgCl_2_, 2.5 mM CaCl_2_, 20 mM D-glucose, and 0.5 mM sodium ascorbate (302–310 mosmol, pH 7.4, saturated with 95% O_2_/ 5% CO_2_, perfused at 3 ml/min) at 30–32°C. For whole-cell recordings, a K^+^-based pipette solution containing 142 mM K^+^-gluconate, 10 mM HEPES, 1 mM EGTA, 2.5 mM MgCl_2_, 4 mM ATP-Mg, 0.3 mM GTP-Na, and 10 mM Na_2_-phosphocreatine (288–294 mosmol, pH 7.32-7.37) or a Cs^+^-based pipette solution containing 121 mM Cs^+^- methanesulphonate, 10 mM HEPES, 10 mM EGTA, 2.5 mM MgCl_2_, 4 mM ATP-Mg, 0.3 mM GTP-Na, and 10 mM Na_2_-phosphocreatine (290–300 mosmol, pH 7.30–7.34) was used. Membrane potentials were not corrected for liquid junction potential (experimentally measured as 12.5 mV for the K^+^-based pipette solution and 9.5 mV for Cs^+^-based pipette solution).

Neurons were visualized with video-assisted infrared differential interference contrast imaging, and fluorescent neurons were identified by epifluorescence imaging under a water immersion objective (40×, 0.8 numerical aperture) on an upright SliceScope Pro 1000 microscope (Scientifica) with an infrared CCD camera (IR-1000, DAGE-MTI). Data were low-pass filtered at 4 kHz and acquired at 10 kHz with an Axon Multiclamp 700B amplifier and an Axon Digidata 1550 Data Acquisition System under the control of Clampex 10.5 (Molecular Devices). Data were analyzed offline using AxoGraph X (AxoGraph Scientific).

Neuronal intrinsic excitability was examined in whole-cell current clamp mode with the K^+^- based pipette solution. The resting membrane potential was recorded within the first 1–2 minutes after break-in. The input resistance was measured after balancing bridge by injecting a 500-ms- long hyperpolarizing current pulse (60–120 pA) to generate a small membrane potential hyperpolarization (2–10 mV) from the resting membrane potential. Depolarizing currents were increased in 5- or 10-pA steps to identify rheobase current.

Synaptic currents were recorded in the whole-cell voltage champ mode with the Cs^+^-based patch pipette solution. Only recordings with series resistance below 20 MΩ were included. EPSCs and IPSCs were recorded at the reversal potential for IPSCs (-60 mV) and EPSCs (+10 mV), respectively. Spontaneous EPSCs and IPSCs (sEPSCs and sIPSCs) were recorded in ACSF, and miniature ESPCs and IPSCs (mEPSCs and mIPSCs) in the presence of voltage-gated sodium channel blocker TTX (0.5 μM, Tocris or Hello Bio). A scaled sliding template method (AxoGraph X) was used to detect sPSCs and mPSCs. Data were low-pass filtered at 2 kHz and a 10-ms template (3 ms baseline, -2 pA amplitude, 0.6 ms rise time, and 3 ms decay time) was used to detect sEPSCs and mEPSCs, whereas a 20-ms template (3 ms baseline, 2 pA amplitude, 0.6 ms rise time, and 10 ms decay time) was used for sIPSCs and mIPSCs. The detection threshold was set to 3 times of noise standard deviation for sEPSCs and mEPSCs, and 2.5 times of noise standard deviation for sIPSCs and mIPSCs. Events with an amplitude less than 5 pA were excluded.

For synaptic scaling analysis, 500 mEPSCs or mIPSCs were randomly selected from each neuron and used in the conventional (*13*) and an improved test (*38*) of multiplicative scaling. In the conventional test, individual Kir2.1 mEPSC or mIPSC amplitudes were ranked and plotted against the ranked control mEPSC or mIPSC amplitudes, respectively. The data were fitted by a linear regression *Y* = *k*X + *m*, where *X*, *Y*, *k*, and *m* are the control amplitude, Kir2.1 amplitude, scaling factor, and constant, respectively. The *k* and *m* obtained from the fitting were then used to scale the Kir2.1 distributions. The scaled Kir2.1 mEPSC or mIPSC distributions were compared to the control mEPSC or mIPSC distributions, respectively, by the Kolmogorov-Smirnov test. In the improved test, individual Kir2.1 mEPSC amplitudes or control mIPSC amplitudes were scaled by different factors according to *A_scaled_* = *kA*, where *A*, *A_scaled_*, and *k* are the amplitude, scaled amplitude, and scaling factor, respectively, and then the subthreshold *A_scaled_* data (< 5 pA) were excluded to generate a series of scaled distributions. The scaled Kir2.1 mEPSC and control mIPSC distributions were compared to the control mEPSC and Kir2.1 mIPSC distributions, respectively, by the Kolmogorov-Smirnov test. The best scaled distribution is the one that yields the largest *P* value.

For optogenetic stimulation, blue light (455 nm) emitted from a collimated light-emitting diode (LED) was used for activating ChR2 and red light (617 nm) for ReaChR. LEDs were driven by a 2-channel LED driver (Mightex Systems) under the control of an Axon Digidata 1550 Data Acquisition system and Clampex 10.5 software. Light was delivered through the reflected light fluorescence illuminator port and the 40× objective. For light pulse stimulation, pulse duration (0.5–5 ms) and intensity (1.61–7.21 mW/mm^2^) were adjusted for each recording to evoke small (to minimize voltage-clamp errors) but reliable monosynaptic EPSCs. Disynaptic IPSCs were evoked using the same light pulse setting as for the corresponding monosynaptic EPSCs. Light pulses were delivered at 20- or 30-second interstimulus intervals.

To isolate layer 5 excitatory neurons-mediated quantal EPSCs (qEPSCs), AMPA receptor currents were recorded at -70 mV membrane potential in the presence of TTX (1 μM), NMDA receptor blocker CPP (10 μM, Tocris or Hello Bio) and GABAA receptor blocker SR95531 (10 μM, Tocris or Hello Bio) in ACSF. Typically, 10–40 sweeps were recorded for each neuron with a 40-second interval between sweeps. For each sweep, mEPSCs were recorded during a 10- second baseline period and a 10-second blue light stimulation period. The light intensity (455 nm, 0.32–7.21 mW/mm2) was ramped down to reduce the tonic currents. mEPSCs were detected using the same method as described above. The qEPSC parameters and average traces were computed as *A_quantal_* = (A_light_f_light_ − A_baseline_f_baseline_)/(f_light_ − f_baseline_), where Aquantal, *A_light_*, and *A_baseline_* are the amplitude, decay time constant, or average trace of qEPSCs, mEPSCs during light stimulation period, and mEPSCs during baseline period, respectively; *f_light_* and *f_baseline_* are the frequency of mEPSCs during light stimulation period and baseline period, respectively.

To evaluate the amount of glutamate released by a presynaptic vesicle, AMPA-EPSCs were first recorded in the presence of CPP (10 μM) in ACSF and then in the presence of CPP (10 μM) and γ-DGG (1 mM, Hello Bio) in response to photostimulation of layer 5.

To determine the presynaptic release probability of layer 5-mediated excitation, NMDA receptor currents were recorded at +40 mV membrane potential in the presence of AMPA receptor blocker NBQX (10 μM, Tocris or Hello Bio) and SR95531 (10 μM), while layer 5 excitatory neurons were repetitively photostimulated. Interstimulus interval was set at 50 s, which minimized the rundown of NMDA receptor currents to be less than 5% with 30 stimuli. The responses to the first 10 stimuli were recorded as the baseline, followed by 20 stimuli in the presence of a use-dependent, open NMDA receptor blocker MK-801 (10 μM, Hello Bio). The rate of NMDA receptor current blockade was measured, which is proportional to presynaptic release probability (*52*, *53*).

To determine AMPA-EPSC and NMDA-EPSC ratio (A/N ratio), AMPA-EPSCs were obtained by subtracting the EPSCs recorded at -60 mV membrane potential in the presence of NBQX (10 μM) and SR95531 (10 μM) from the EPSCs in ACSF. NMDA-EPSCs were then recorded at +40 mV membrane potential in the presence of NBQX (10 μM) and SR95531 (10 μM).

### Immunohistochemistry, confocal microscopy, and neuronal morphology reconstruction

To estimate the fraction of layer 2/3 pyramidal cells expressing Kir2.1, after electrophysiological recordings brain slices (300 μm) were fixed overnight in 4% paraformaldehyde in phosphate buffered saline (PBS, pH 7.4), cryoprotected with 30% sucrose in PBS, and then frozen for cryo- sectioning into 50-μm-thick coronal slices using a HM 450 Sliding Microtome (Thermo Scientific). Slices were stained with the primary antibody, guinea pig anti-NeuN (EMD Millipore Corporation, catalog # ABN90, lot # 4878142, 1:1000) and then secondary antibody, goat anti- guinea pig IgG (H+L) conjugated with Alexa Flour 647 (Invitrogen, catalog # A21450, lot # 1903515, 1:1000). Images were acquired on a TCS SP8 X Confocal Microscope (Leica) using a 63× oil objective at the 1× zoom. The numbers of Kir2.1 neurons (i.e., tdTomato positive) and NeuN positive cells were counted in layer 2/3 of the V1 using Imaris 9.5.0 (Oxford Instruments). Assuming that layer 2/3 in the V1 consists of 85% excitatory and 15% inhibitory neurons, Kir2.1 transfection rates in pyramidal cells were estimated as the number of tdTomato positive cells divided by the number of 85% NeuN positive cells.

To quantify neuronal morphology and synapse number, layer 5 excitatory neurons were transduced with AAV9-EF1α-DIO-hChR2(H134R)-P2A-EYFP and AAV9-hSyn-DIO- Synaptophysin-tdTomato in *Rbp4-Cre(KL100)* mice. A subset of layer 2/3 pyramidal neurons was electroporated with pCAG-Kir2.1-P2A-3×NLS-dTomato. During whole cell recordings, the Kir2.1 and control neurons were filled with 0.1% biocytin (Tocris) included in the pipette internal solution. Recorded slices were fixed with 4% paraformaldehyde in PBS (pH 7.4) overnight, rinsed with PBS for 3 times (15 minutes each), permeabilized in 0.2% Triton X-100 (Sigma-Aldrich) in PBS (PBS-T) at room temperature for 1 hour, and incubated in streptavidin- Dylight 405 (Invitrogen, 1:1000 in PBS-T) at room temperature for 2 hours. The slices were then rinsed with PBS for 3 times (15 minutes each) and mounted in ProLong Diamond Antifade Mountant (Invitrogen). Tile scanned z-stack images of the biocytin-filled Kir2.1 and control neurons were acquired on an TCS SP8 X Confocal Microscope (Leica) using a 63× oil objective at the 4× zoom. Sequential scans were performed between frames for biocytin-Dylight 405 (excitation 405 nm, emission 430–500 nm) and tdTomato (excitation 558 nm, emission 570–650 nm) signals. A series of images were acquired to cover an area of 46 μm × 46 μm and depth of 30–45 μm with the z-stack interval of 0.3 μm. The images were stitched together to cover the entire somatodendritic regions of both Kir2.1 and control neurons.

The somatodendritic regions of Kir2.1 and control neurons were reconstructed based on the biocytin-Dylight 405 signal by the Filament Tracer semi-automatic tools in Imaris 9.5.0 (Oxford Instruments). Spines were detected along the dendrites with estimated diameter of 0.5 μm.

Presynaptic boutons marked by Synaptophysin-tdTomato originating from layer 5 excitatory neurons were detected using the automatic adding spots function with an estimated diameter of 0.5 μm. A Synatophysin-tdTomato spot colocalized with a dendritic spine (distance less than 0.5 μm) was considered as a synapse. The dendrite lengths and numbers of spines and synapses were measured for both Kir2.1 and control neurons. Based on the morphology, spines were classified into subtypes: mushroom (the ratio of head max width to neck mean width >1 and the head max width ≥ 0.4 μm), thin (the ratio of spine length to head mean width > 2 and the ratio of head max width to neck mean width > 1), filopodia (spine length > 2 μm and the ratio of head mean width to neck mean width ≤ 1), and stubby (the remaining spines)(*74–78*).

### Statistics

All reported sample numbers (*n*) represent biological replicates. Statistical analyses were performed with Prism 11 (GraphPad Software). The D’Agostino-Pearson, Anderson-Darling, Shapiro-Wilk, and Kolmogorov-Smirnov tests were used to determine if data were normally distributed. If all data within an experiment passed all four normality tests, statistical tests assuming a Gaussian distribution were used. Otherwise, non-Gaussian distribution tests were applied. All statistical tests were two-tailed with an alpha of 0.05. Sex effect was inspected by two-way or three-way ANOVA. The details of all statistical tests, numbers of replicates, and *P* values are reported in Supplementary table 1.

---

**Extended Data Figure 1.**
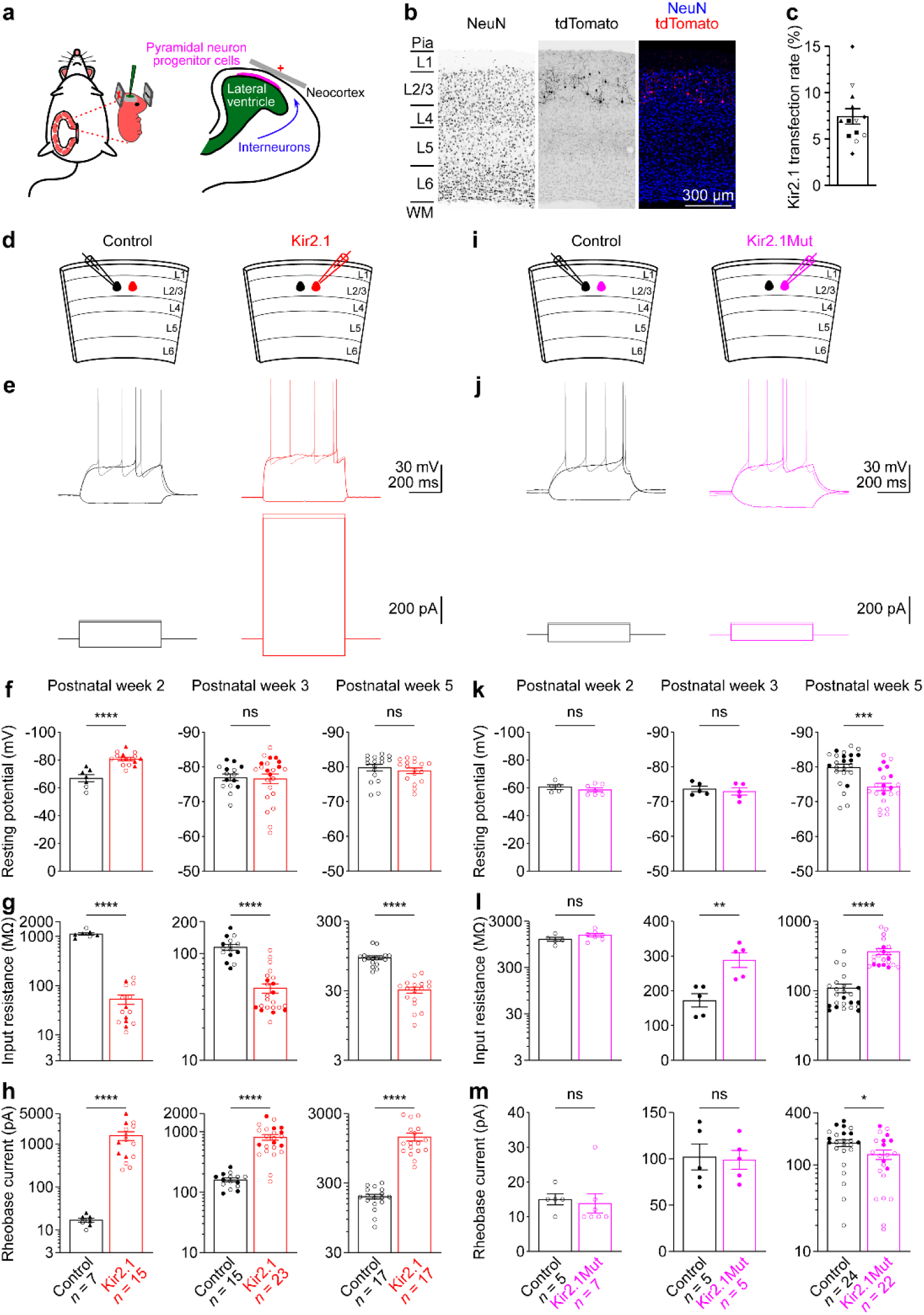
Neuronal intrinsic excitability in Kir2.1 or Kir2.1Mut neurons. **(a)** Schematic of *in utero* electroporation. **(b)** Representative fluorescence images of a V1 slice showing Kir2.1-T2A-tdTomato expression in layer 2/3. NeuN staining used to label neurons. (**c**) Summary data of fraction of layer 2/3 pyramidal neurons expressing Kir2.1-T2A-tdTomato. Each filled (male) or open (female) symbol represents a slice, and different symbols represent different mice. Data are mean ± s.e.m. *n* =13 slices from 5 mice (2–3 slices per mouse). **(d)** Schematics of slice experiments in **e** with Kir2.1 in a subset of layer 2/3 pyramidal cells. **(e)** Membrane potentials (upper panels) in response to current injections (lower panels) in control and Kir2.1 neurons. **(f**‒**h)** Summary data of resting membrane potentials (**f**), input resistances (**g**), and rheobase currents (**h**) in control and Kir2.1 neurons at postnatal weeks 2 (P8‒9), 3 (P17‒ 20), and 5 (P28‒35). Each filled (male) or open (female) circle symbol represents a neuron except for the data from postnatal week 2, where open circle symbols represent neurons from unknown-sex mice that were *in utero* electroporated with pCAG-Kir2.1-T2A-tdTomato or pCAG-Kir2.1Mut-T2A-tdTomato, whereas filled triangle symbols represent neurons from male mice that were *in utero* electroporated with pAAV-EF1α-DIO-Kir2.1-T2A-tdTomato and injected with AAV9-hSyn-HI-eGFP-Cre at P1 to induce Kir2.1 expression. Data are mean ± s.e.m. Note hyperpolarized resting membrane potentials at postnatal week 2, decreased resting input resistances, and increased rheobase currents at all ages in Kir2.1 neurons. **(i**‒**m)** As in **d**‒**h**, but for a non-conducting mutant Kir2.1 (Kir2.1Mut). Note increased input resistances at postnatal weeks 3 and 5, depolarized resting membrane potentials, and decreased rheobase currents only at postnatal week 5 in Kir2.1Mut neurons. The numbers of recorded neurons are indicated in the panels. ns, *P* > 0.05; *, *P* < 0.05; **, *P* < 0.01; ***, *P* < 0.0001; ****, *P* < 0.0001.

**Extended Data Figure 2.**
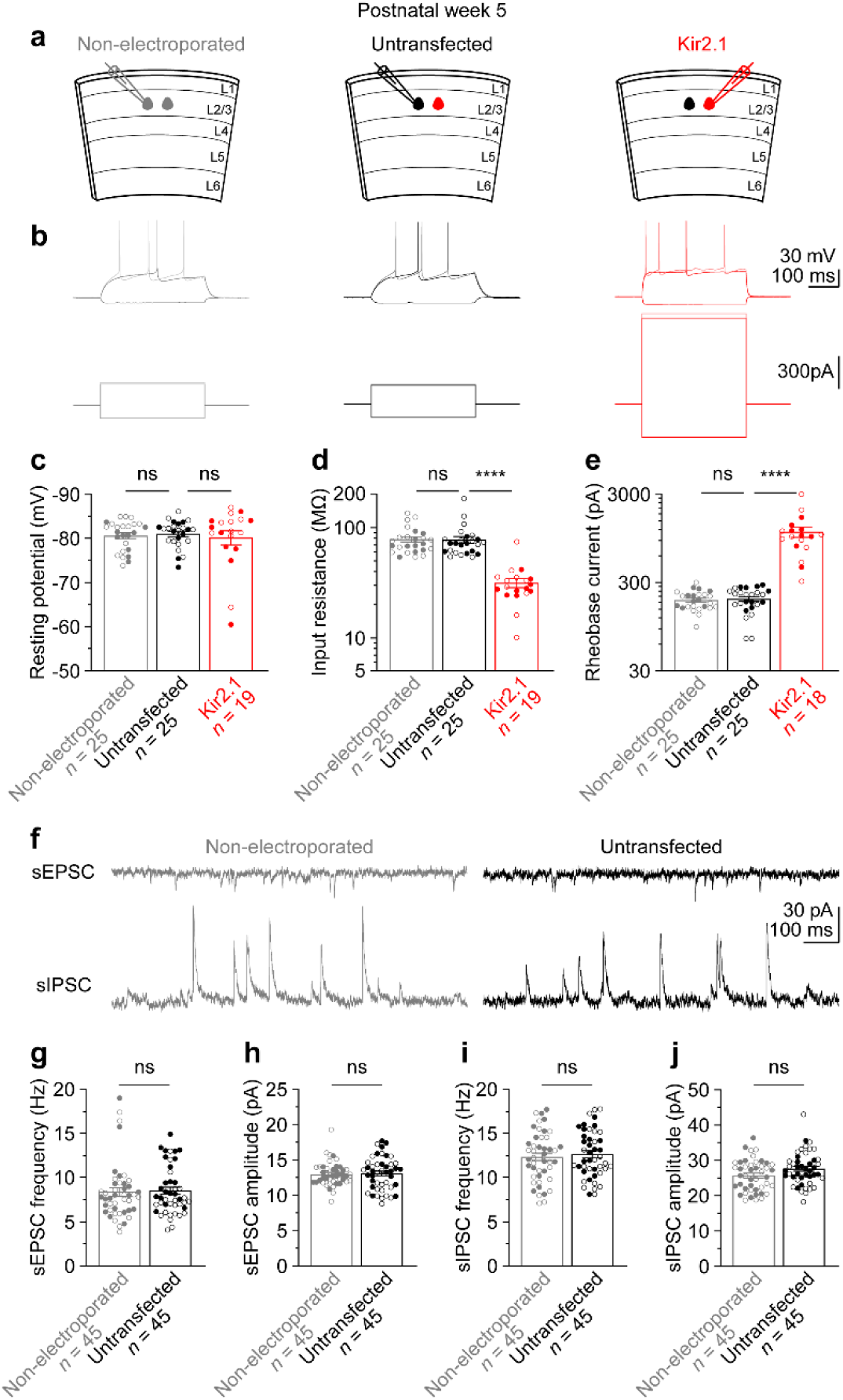
Kir2.1 overexpression in a subset of layer 2/3 pyramidal cells does not affect neuronal intrinsic excitability and synaptic inputs of untransfected pyramidal cells. **(a)** Schematics of slice experiments in **b** and **f** with no electroporation or Kir2.1 in a subset of layer 2/3 pyramidal cells at postnatal week 5. **(b)** Membrane potentials (upper panels) in response to current injections (lower panels) from a pyramidal cell in a non-electroporated littermate mouse (left, grey) and an untransfected control neuron (middle, black) and a Kir2.1 neuron (right, red) in an electroporated mouse. **(c**‒**e)** Summary data of resting membrane potentials (**c**), input resistances (**d**), and rheobase currents (**e**). Each filled (male) or open (female) symbol represents a neuron. Data are mean ± s.e.m. Note similar resting membrane potentials, input resistances, and rheobase currents between non-electroporated and untransfected neurons. **(f)** Spontaneous EPSCs and IPSCs from a non-electroporated (left, grey) and an untransfected neuron (right, black). **(g**‒**j)** Summary data of sEPSC frequencies (**g**), amplitudes (**h**), sIPSC frequencies (**i**), and amplitudes (**j**). Note similar sEPSCs and sIPSCs between non-electroporated and untransfected neurons. Each filled (male) or open (female) symbol represents a neuron. Data are mean ± s.e.m. The numbers of recorded neurons are indicated in the panels. ns, *P* > 0.05; ****, *P* < 0.0001.

**Extended Data Figure 3.**
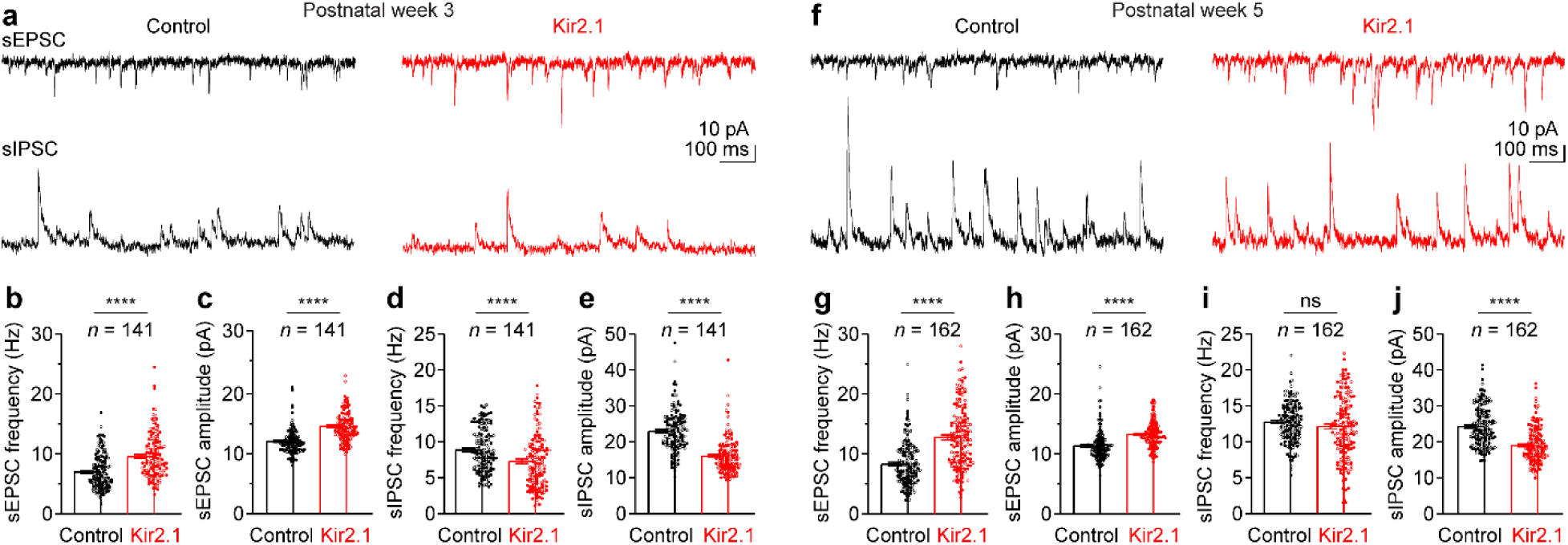
Increased sEPSCs and decreased sIPSCs in Kir2.1 neurons. **(a)** sEPSCs and sIPSCs from a recorded control and Kir2.1 neurons at postnatal week 3. **(b**‒**e)** Summary data of frequency and amplitude of sEPSCs (**b,c**) and sIPSCs (**d**,**e**). Each filled (male) or open (female) symbol represents a recorded neuron. Data are mean ± s.e.m. Note increased sEPSC frequencies and amplitudes of sEPSCs and decreased sIPSC frequencies and amplitudes in Kir2.1 neurons. **(f**‒**j)** As in **a**‒**e**, but for postnatal week 5. Note increased sEPSC frequencies and amplitudes and decreased sIPSCs amplitudes in Kir2.1 neurons. The numbers of pairs of recorded neurons (*n*) are indicated in the panels. Ns, *P* > 0.05; ****, *P* < 0.0001.

**Extended Data Figure 4.**
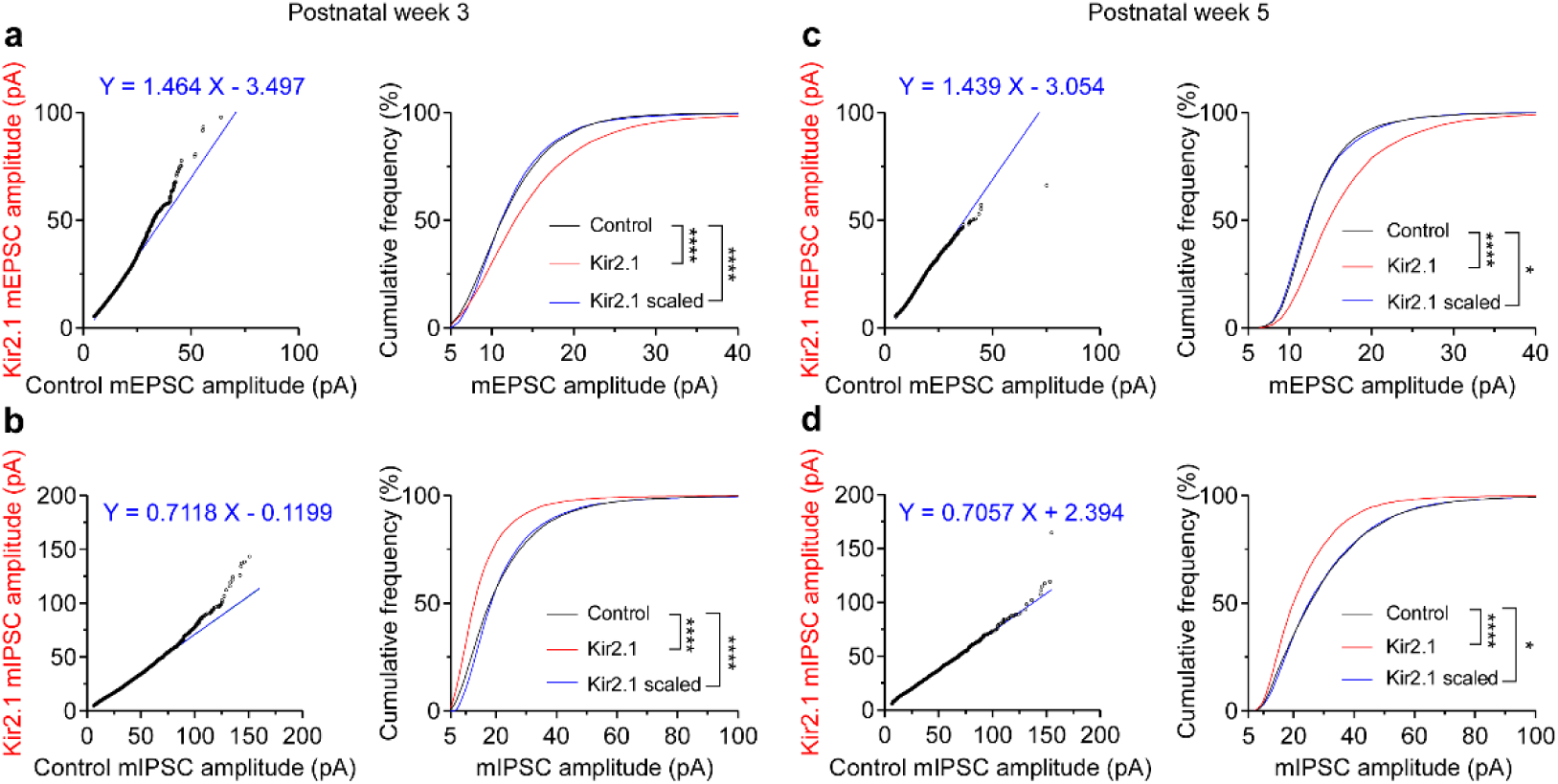
Scaling analysis of mEPSCs and mIPSCs with best fit equations. **(a)** Left, ranked mEPSC amplitudes were plotted for 31 pairs of control and Kir2.1 neurons at postnatal week 3, with 500 randomly selected events per neuron. Each data point represents an individual mEPSC amplitude. The relationship was fitted with a linear regression to obtain the best-fit equation indicated in the panel. Right, cumulative distributions of mEPSC amplitudes. Kir2.1 mEPSC amplitudes were transformed by the best-fit equation to obtain the scaled Kir2.1 distribution, which is significantly different from the control distribution. **(b)** As in **a**, but for mIPSC amplitudes from 36 pairs of control and Kir2.1 neurons. The scaled Kir2.1 distribution is significantly different from the control distribution. **(c,d)** As in **a,b**, but for 10 pairs of control and Kir2.1 neurons at postnatal week 5. The scaled Kir2.1 mEPSC and mIPSC amplitude distributions are both significantly different from the control distributions. *, *P* < 0.05; ****, *P* < 0.0001.

**Extended Data Figure 5.**
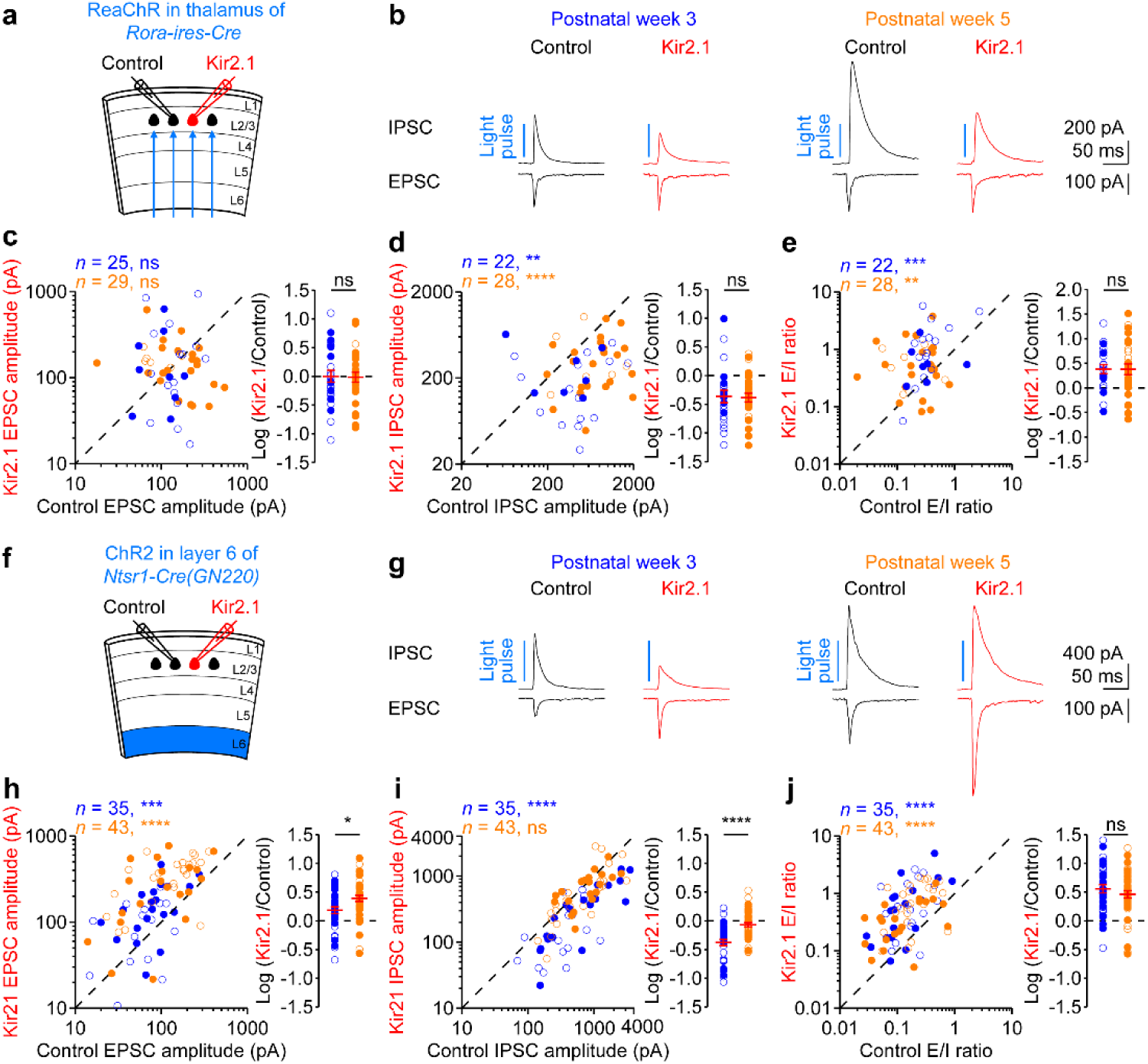
Excitation from layer 6 but not thalamus is increased in Kir2.1 neurons. **(a)** Schematic of slice experiments in **b** with Kir2.1 in layer 2/3 pyramidal cells and ReaChR in the axons from thalamic excitatory neurons. **(b)** Monosynaptic EPSCs and disynaptic IPSCs from a pair of simultaneously recorded control and Kir2.1 neurons in response to photostimulation. **(c)** Left, summary data of thalamus-evoked EPSC amplitudes in control and Kir2.1 neurons at postnatal weeks 3 (blue) and 5 (orange). Each filled (male) or open (female) symbol represents a pair of control and Kir2.1 neurons. Right, logarithm of the ratios between EPSC amplitudes from pairs of Kir2.1 and control neurons. Red, mean ± s.e.m. **(d**,**e)** As in **c**, but for IPSC amplitudes and E/I ratios. Note similar EPSC amplitudes between control and Kir2.1 neurons, smaller IPSC amplitudes, and larger E/I ratios in Kir2.1 neurons. The decrease of IPSC amplitude and increase of E/I ratio in Kir2.1 neurons are similar between the two ages. **(f**‒**j)** As in **a**‒**e**, but for ChR2 in layer 6 excitatory neurons. Note larger EPSC amplitudes at both ages, smaller IPSC amplitudes at postnatal week 3, and larger E/I ratios at both ages in Kir2.1 neurons. The increase of EPSC amplitude in Kir2.1 neurons is larger at postnatal week 5 than week 3, but the increase of E/I ratio is similar between the two ages. The numbers of pairs of recorded neurons (*n*) are indicated in the panels. ns, *P* > 0.05; **, *P* < 0.01; ***, *P* < 0.001; ****, *P* < 0.0001.

**Extended Data Figure 6.**
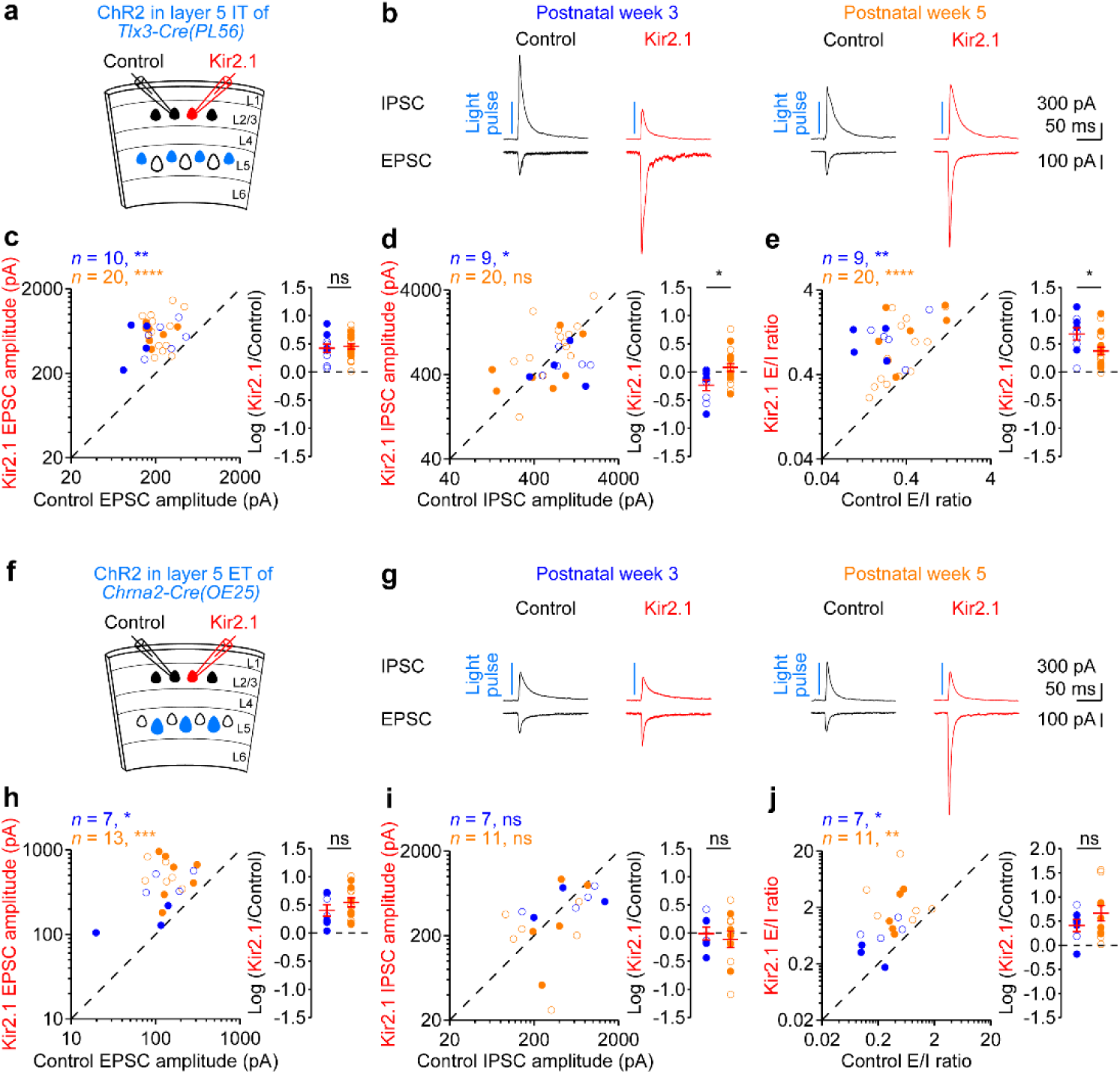
Increased excitation from both layer 5 IT and ET neurons onto Kir2.1 neurons. **(a)** Schematic of slice experiments in **b** with Kir2.1 in layer 2/3 pyramidal cells and ChR2 in layer 5 IT excitatory neurons. **(b)** Monosynaptic EPSCs and disynaptic IPSCs from a pair of simultaneously recorded control and Kir2.1 neurons in response to photostimulation. **(c)** Left, summary data of layer 5 IT-evoked EPSC amplitudes in control and Kir2.1 neurons at postnatal weeks 3 (blue) and 5 (orange). Each filled (male) or open (female) symbol represents a pair of control and Kir2.1 neurons. Right, logarithm of the ratios between EPSC amplitudes from pairs of Kir2.1 and control neurons. Red, mean ± s.e.m. **(d,e)** As in **c**, but for IPSC amplitudes and E/I ratios. Note larger EPSC amplitudes at both ages, smaller IPSC amplitudes at postnatal week 3, and larger E/I ratios at both ages in Kir2.1 neurons. The increase of EPSC amplitude in Kir2.1 neurons is similar between the two ages, but the increase of E/I ratio is smaller at postnatal week 5 than week 3. **(f**‒**j)** As in **a**‒**e**, but for ChR2 in layer 5 ET excitatory neurons. Note similar IPSC amplitudes between Kir2.1 and control neurons, larger EPSC amplitudes, and larger E/I ratios in Kir2.1 neurons at both ages. The increase of EPSC amplitude and E/I ratio in Kir2.1 neurons is similar between the two ages. The numbers of pairs of recorded neurons (*n*) are indicated in the panels. ns, *P* > 0.05; *, *P* < 0.05; **, *P* < 0.01; ***, *P* < 0.001; ****, *P* < 0.0001.

**Extended Data Figure 7.**
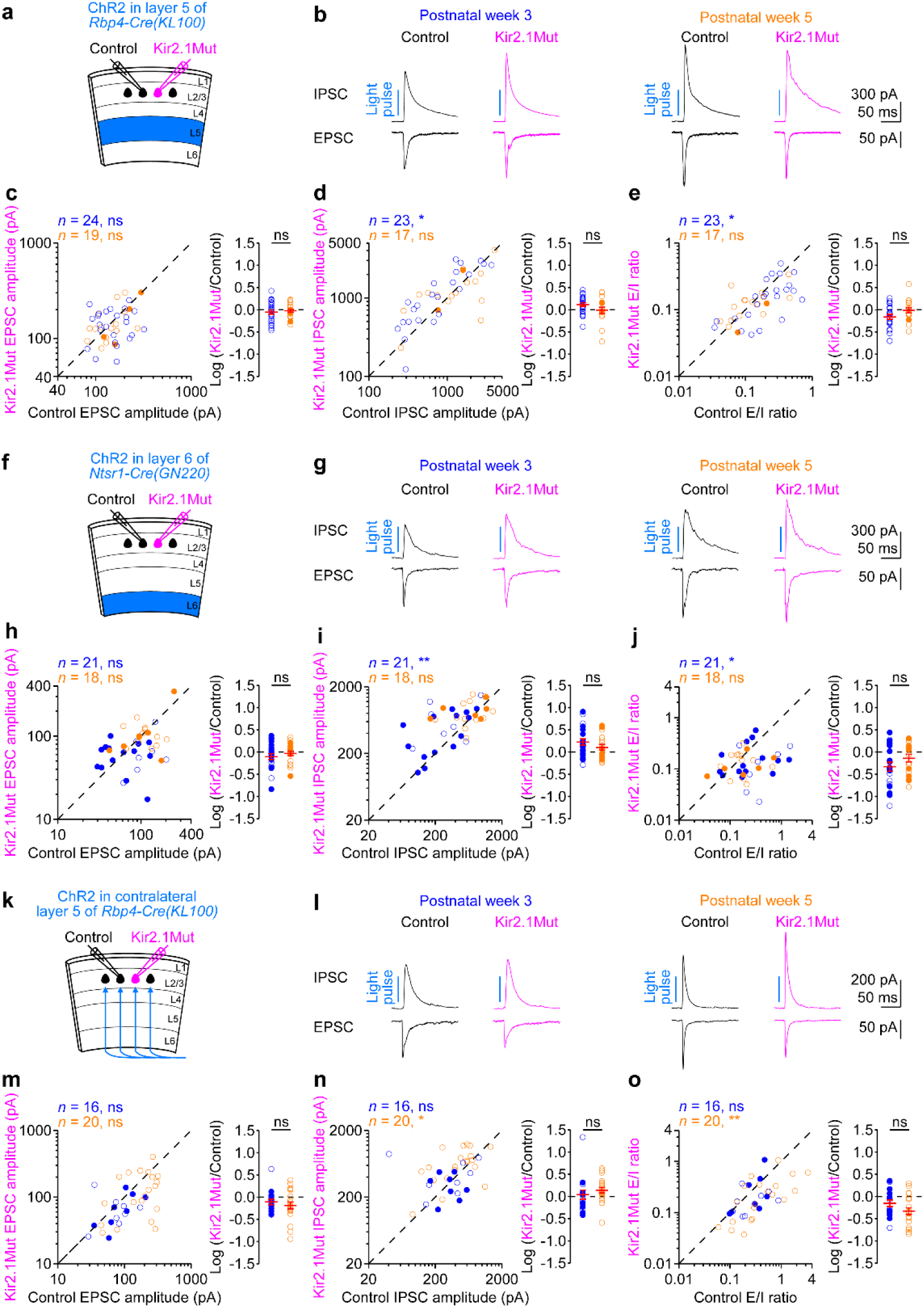
Kir2.1Mut overexpression in layer 2/3 pyramidal cells does not affect excitation from layer 5, layer 6, and contralateral layer 5. **(a)** Schematic of slice experiments in **b** with Kir2.1Mut in layer 2/3 pyramidal cells and ChR2 in layer 5 excitatory neurons. **(b)** Monosynaptic EPSCs and disynaptic IPSCs from a pair of simultaneously recorded control and Kir2.1Mut neurons in response to photostimulation. **(c)** Left, summary data of layer 5-evoked EPSC amplitudes in control and Kir2.1Mut neurons at postnatal weeks 3 (blue) and 5 (orange). Each filled (male) or open (female) symbol represents a pair of control and Kir2.1Mut neurons. Right, logarithm of the ratios between EPSC amplitudes from pairs of Kir2.1Mut and control neurons. Red, mean ± s.e.m. **(d**,**e)** As in **c**, but for IPSC amplitudes and E/I ratios. Note similar EPSC amplitudes between control and Kir2.1Mut neurons at both ages, larger IPSC amplitudes, and smaller E/I ratios in Kir2.1Mut neurons only at postnatal week 3. **(f**‒**j)** As in **a**‒**e**, but for slice experiments with ChR2 in layer 6 excitatory neurons. Note similar EPSC amplitudes between control and Kir2.1Mut neurons at both ages, larger IPSC amplitudes, and smaller E/I ratios in Kir2.1Mut neurons only at postnatal week 3. **(k**‒**o)** As in **a**‒**e**, but for slice experiments with ChR2 in the axons from contralateral layer 5 excitatory neurons. Note similar EPSC amplitudes between control and Kir2.1Mut neurons at both ages, large IPSC amplitudes, and smaller E/I ratios in Kir2.1Mut neurons only at postnatal week 5. The numbers of pairs of recorded neurons (*n*) are indicated in the panels. ns, *P* > 0.05; *, *P* < 0.05; **, *P* < 0.01.

**Extended Data Figure 8.**
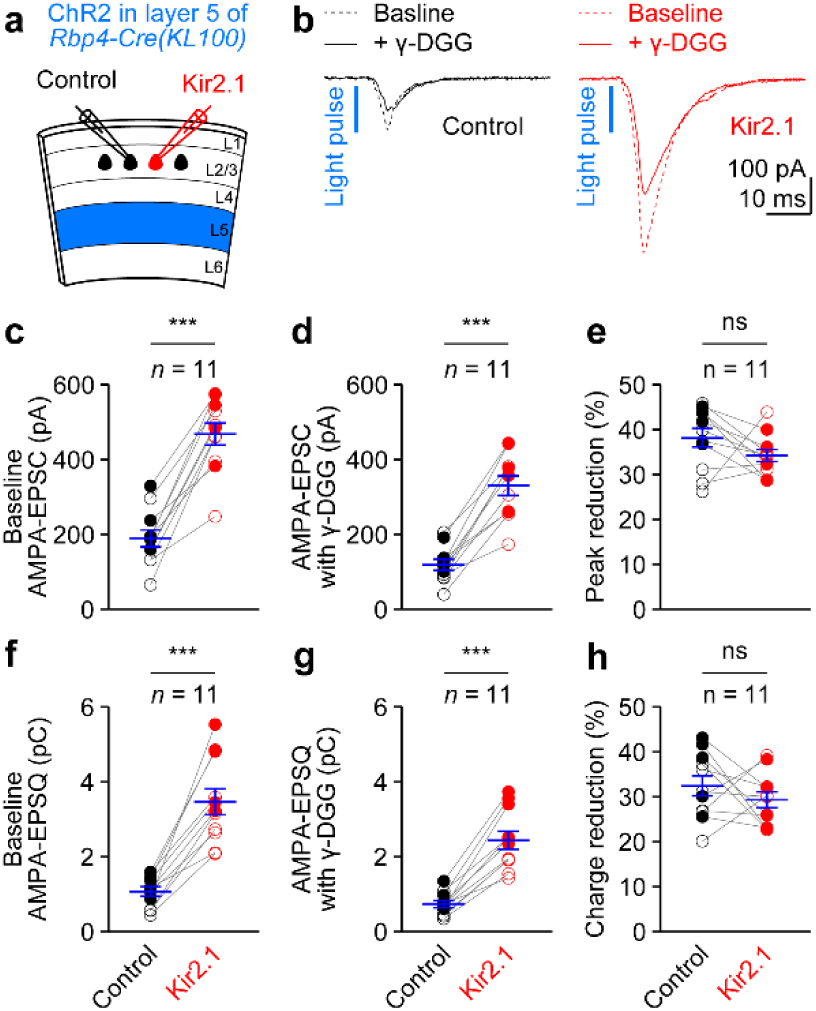
Unaltered amount of glutamate released from presynaptic vesicles. **(a)** Schematic of slice experiments in **b** with Kir2.1 in layer 2/3 pyramidal cells and ChR2 in layer 5 excitatory neurons at postnatal week 5. (**b**) AMPA-EPSCs before (dashed traces) and after 1 mM γ-DGG application (solid traces) in ACSF containing CPP (NMDA receptor antagonist) from a pair of simultaneously recorded control and Kir2.1 neurons in response to photostimulation. **(c**‒**h)** Summary data of layer 5-evoked AMPA-EPSC amplitudes and charges before (**c**,**f**) and after γ-DGG application (**d**,**g**), and degree of reduction by γ-DGG (**e**,**h**) from pairs of Kir2.1 and control neurons. Each pair of filled (male) or open (female) symbols represents a pair of control and Kir2.1 neurons. Blue, mean ± s.e.m. Note similar degree of reduction of AMPA-EPSC amplitudes and charges by γ-DGG between Kir2.1 and control neurons. The numbers of pairs of recorded neurons (*n*) are indicated in the panels. ns, *P* > 0.05; ***, *P* < 0.001.

**Extended Data Figure 9.**
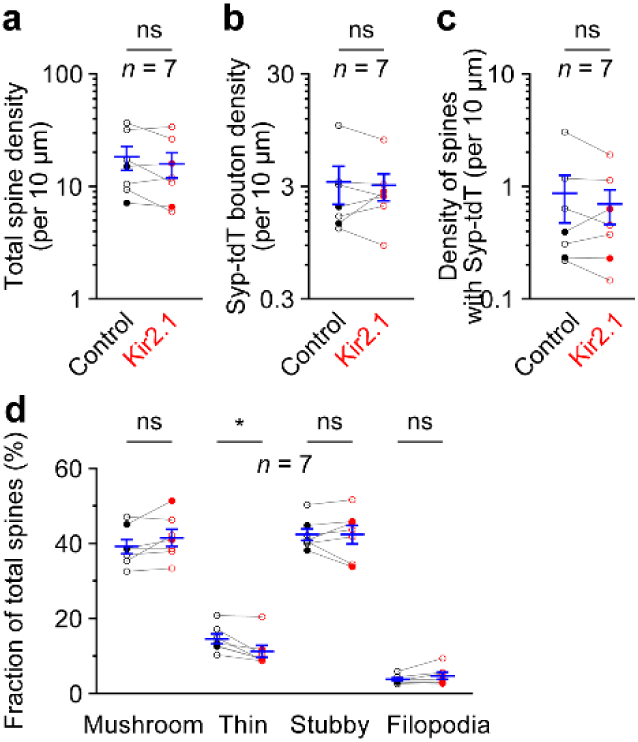
Densities of total spines, Syp-tdT boutons, spines with Syp-tdT, and fraction of total spines. **(a**‒**d)** The densities of total spines (**a**), Syp-tdT boutons on both dendrites and spines (**b**), spines colocalized with Syp-tdT (**c**), and fractions of different types of spines (**d**) from pairs of control and Kir2.1 neurons at postnatal week 5. Each pair of filled (male) or open (female) symbols represents a pair of control and Kir2.1 neurons. Blue, mean ± s.e.m. Note decreased thin spines in Kir2.1 neurons. The numbers of pairs of neurons (*n*) are indicated in the panels. ns, *P* > 0.05; *, *P* < 0.05.

**Extended Data Figure 10.**
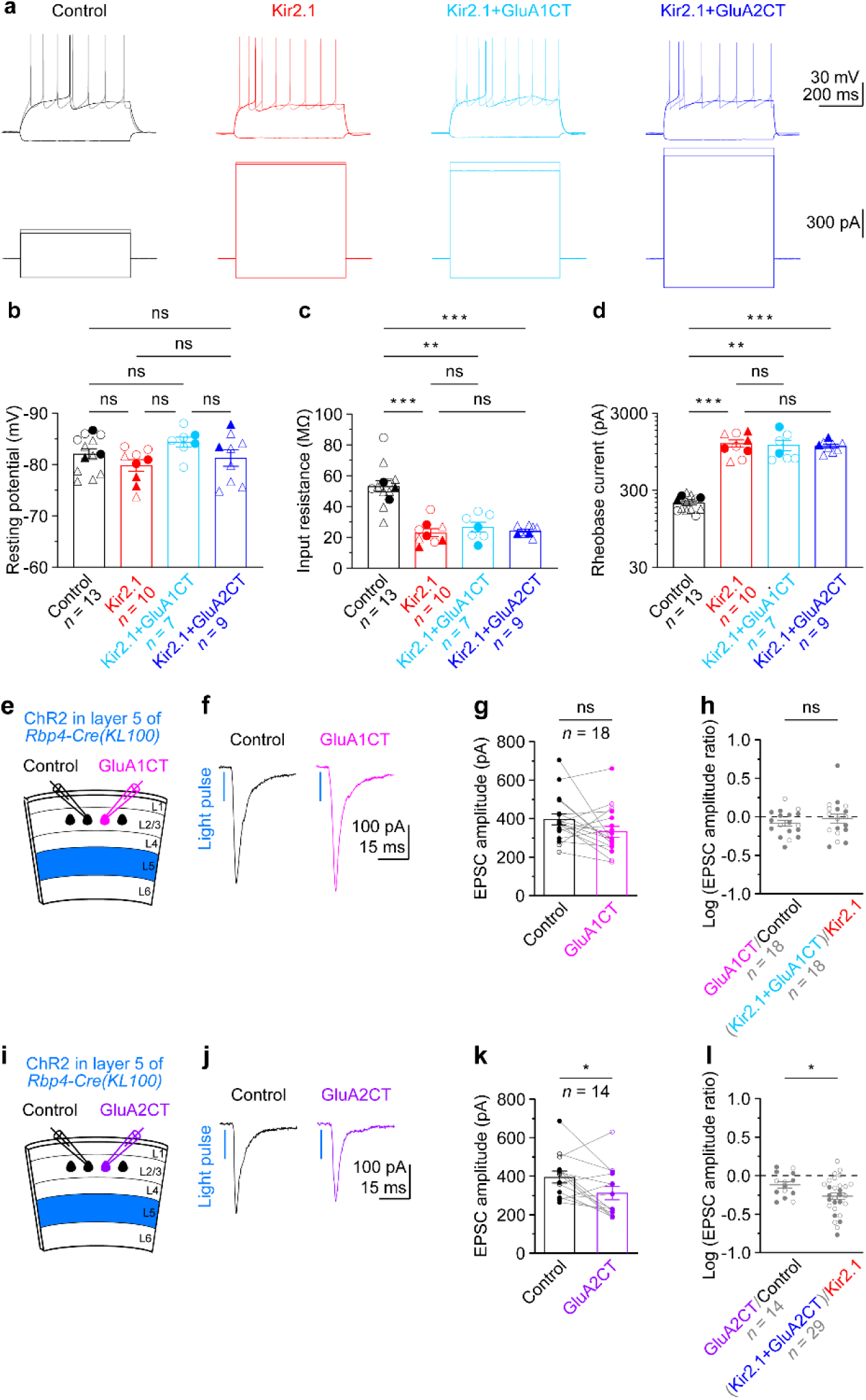
The impacts of GluA1CT or GluA2CT on neuronal intrinsic excitability in Kir2.1 neurons and layer 5 excitatory inputs in control neurons. **(a)** Membrane potentials (upper panels) in response to current injections (lower panels) from a control, Kir2.1, Kir2.1 and GluA1CT co-expressing (Kir2.1+GluA1CT), and Kir2.1 and GluA2CT co-expressing (Kir2.1+GluA2CT) neurons. **(b**‒**d)** Summary data of resting membrane potentials (**b**), input resistances (**c**), and rheobase currents (**d**) in control, Kir2.1, Kir2.1+GluA1CT, and Kir2.1+GluA2CT neurons at postnatal week 5. Each filled (male) or open (female) symbol represents a neuron. Circle symbols are from mice electroporated with Kir2.1 and GluA1CT, and triangle symbols from mice with Kir2.1 and GluA2CT. Note similar resting membrane potentials in all four groups. Input resistances are reduced, whereas rheobase currents are increased to the same levels in Kir2.1, Kir2.1+GluA1CT, and Kir2.1+GluA2CT neurons as compared to control neurons. **(e)** Schematic of slice experiments in **f** with GluA1CT in layer 2/3 pyramidal cells and ChR2 in layer 5 excitatory neurons. **(f)** Monosynaptic EPSCs from a pair of simultaneously recorded control and GluA1CT neurons in response to photostimulation. **(g)** Summary data of EPSC amplitudes in control and GluA1CT neurons. Each pair of filled (male) or open (female) symbol represents a pair of control and GluA1CT neurons. Bars, mean ± s.e.m. Note similar EPSC amplitudes between control and GluA1CT neurons. **(h)** Logarithm of the ratios between EPSC amplitudes from pairs of GluA1CT and control neurons or Kir2.1/GluA1CT and Kir2.1 neurons (data from Figure 6h). Each filled (male) or open (female) symbol represents a pair of neurons. Lines, mean ± s.e.m. Note similar logarithmic ratios between both groups. **(i**‒**l)** As in **e**‒**h**, but for GluA2CT. Note mild reduction of EPSC amplitudes in GluA2CT neurons compared to control neurons and more pronounced reduction in Kir2.1/GluA2CT neurons compared to Kir2.1 neurons (data from Figure 6k). The numbers of recorded neurons (*n*) are indicated in the panels. ns, *P* > 0.05; *, *P* < 0.05; **, *P* < 0.01; ***, *P* < 0.001.

