## Supplemental Table 1 for "Synaptic input-selective homeostasis safeguards developing cortical neurons toward setpoint activity"

Supplemental table 1 (PW, Postnatal Week)

| Figure | Experiment | Parameter | Statistical test | n (neuron) of N (mouse) or n (event, noted) | P value | Sex Effect (P value) | Sex Difference (P value) | |
|---|---|---|---|---|---|---|---|---|
| | | | | | | | Male | Female |
| Fig. 1e | In vivo spike recordings: Control vs Kir2.1 | Spontaneous spike rate, PW2 (P10-13) | Mann Whitney test | 15 of 3 vs 17 of 3 | <0.0001 | | | |
| | | Spontaneous spike rate, PW3 (P15-21) | Mann Whitney test | 49 of 7 vs 47 of 7 | <0.0001 | 0.0296 | 0.0467 | 0.8311 |
| | | Spontaneous spike rate, PW4 (P23-28) | Mann Whitney test | 19 of 4 vs 18 of 4 | 0.9223 | 0.2271 | | |
| | | Spontaneous spike rate, PW5 (P34-38) | Mann Whitney test | 29 of 5 vs 27 of 5 | 0.0605 | 0.0498 | 0.0005 | 0.7464 |
| Fig. 1f | | Visual-evoked spike rate, PW2 (P10-13) | Mann Whitney test | 15 of 3 vs 17 of 3 | 0.0072 | | | |
| | | Visual-evoked spike rate, PW3 (P15-21) | Mann Whitney test | 49 of 7 vs 47 of 7 | 0.0037 | 0.075 | | |
| | | Visual-evoked spike rate, PW4 (P23-28) | Mann Whitney test | 19 of 4 vs 18 of 4 | 0.7525 | 0.8616 | | |
| | | Visual-evoked spike rate, PW5 (P34-38) | Mann Whitney test | 29 of 5 vs 27 of 5 | 0.2129 | 0.1765 | | |
| Fig. 1g | | Overall spike rate, PW2 (P10-13) | Mann Whitney test | 15 of 3 vs 17 of 3 | <0.0001 | | | |
| | | Overall spike rate, PW3 (P15-21) | Mann Whitney test | 49 of 7 vs 47 of 7 | <0.0001 | 0.4598 | | |
| | | Overall spike rate, PW4 (P23-28) | Mann Whitney test | 19 of 4 vs 18 of 4 | 0.8746 | 0.3725 | | |
| | | Overall spike rate, PW5 (P34-38) | Mann Whitney test | 29 of 5 vs 27 of 5 | 0.0551 | 0.0343 | 0.0004 | 0.8164 |
| Fig. 2b | mEPSC (PW3): Control vs Kir2.1 | Frequency | Mann Whitney test | 31 of 5 vs 31 of 5 | 0.2933 | 0.0676 | | |
| Fig. 2c | | Amplitude | Unpaired t test with Welch's correction | 31 of 5 vs 31 of 5 | 0.0012 | 0.5001 | | |
| Fig. 2f | | Cumulative distribution of amplitudes (scaled with optimal factor) | Kolmogorov-Smirnov test | Events: 15500 (Control) vs 15500 (Kir2.1) vs 15130 (Kir2.1/1.164) | <0.0001 (Control vs Kir2.1), <0.0001 (Control vs Kir2.1/1.164) | | | |
| Ext Fig.4a | | Cumulative distribution of amplitudes (scaled with best equation) | Kolmogorov-Smirnov test | Events: 15500 (Control) vs 15500 (Kir2.1) vs 15500 (Kir2.1 scaled) | <0.0001 (Control vs Kir2.1), <0.0001 (Control vs Kir2.1 scaled) | | | |
| Fig. 2d | mIPSC (PW3): Control vs Kir2.1 | Frequency | Mann Whitney test | 36 of 5 vs 36 of 5 | 0.0056 | 0.2122 | | |
| Fig. 2e | | Amplitude | Mann Whitney test | 36 of 5 vs 36 of 5 | <0.0001 | 0.0002 | 0.0025 | <0.0001 |
| Fig. 2g | | Cumulative distribution of amplitudes (scaled with optimal factor) | Kolmogorov-Smirnov test | Events: 18000 (Control) vs 18000 (Kir2.1) vs 17322 (Control x 0.7089) | <0.0001 (Control vs Kir2.1), <0.0001 (Kir2.1 vs Control x 0.7089) | | | |
| Ext Fig.4b | | Cumulative distribution of amplitudes (scaled with best equation) | Kolmogorov-Smirnov test | Events: 18000 (Control) vs 18000 (Kir2.1) vs 18000 (Kir2.1 scaled) | <0.0001 (Control vs Kir2.1), <0.0001 (Control vs Kir2.1 scaled) | | | |
| Fig. 2i | mEPSC (PW5): Control vs Kir2.1 | Frequency | Unpaired t test with Welch's correction | 10 of 3 vs 10 of 3 | 0.0358 | 0.09 | | |
| Fig. 2j | | Amplitude | Mann Whitney test | 10 of 3 vs 10 of 3 | 0.0115 | 0.0807 | | |
| Fig. 2m | | Cumulative distribution of amplitudes (scaled with optimal factor) | Kolmogorov-Smirnov test | Events: 5000 (Control) vs 5000 (Kir2.1) vs 4998 (Kir2.1/1.179) | <0.0001 (Control vs Kir2.1), <0.0001 (Control vs Kir2.1/1.179) | | | |
| Ext Fig.4c | | Cumulative distribution of amplitudes (scaled with best equation) | Kolmogorov-Smirnov test | Events: 5000 (Control) vs 5000 (Kir2.1) vs 5000 (Kir2.1 scaled) | <0.0001 (Control vs Kir2.1), 0.0136 (Control vs Kir2.1 scaled) | | | |
| Fig. 2k | mIPSC (PW5): Control vs Kir2.1 | Frequency | Mann Whitney test | 10 of 3 vs 10 of 3 | >0.9999 | 0.0041 | 0.5081 | 0.4107 |
| Fig. 2l | | Amplitude | Mann Whitney test | 10 of 3 vs 10 of 3 | 0.0003 | 0.9675 | | |
| Fig. 2n | | Cumulative distribution of amplitudes (scaled with optimal factor) | Kolmogorov-Smirnov test | Events: 5000 (Control) vs 5000 (Kir2.1) vs 5000 (Control x 0.8315) | <0.0001 (Control vs Kir2.1), <0.0001 (Kir2.1 vs Control x 0.8315) | | | |
| Ext Fig.4d | | Cumulative distribution of amplitudes (scaled with best equation) | Kolmogorov-Smirnov test | Events: 5000 (Control) vs 5000 (Kir2.1) vs 5000 (Kir2.1 scaled) | <0.0001 (Control vs Kir2.1), 0.0495 (Control vs Kir2.1 scaled) | | | |
| Fig. 3c | Layer 2/3 input-evoked: Control vs Kir2.1 | EPSC amplitude (PW3) | Wilcoxon matched-paired signed rank test | 22 of 6 vs 22 of 6 | 0.8237 | 0.0414 | 0.6508 | 0.3069 |
| | | EPSC amplitude (PW5) | Wilcoxon matched-paired signed rank test | 34 of 10 vs 34 of 10 | 0.9328 | 0.4777 | | |
| | | EPSC log ratio (PW3 vs PW5) | Unpaired t test | 22 of 6 vs 34 of 10 | 0.7538 | 0.44 | | |
| Fig. 3d | | Disynaptic IPSC amplitude (PW3) | Wilcoxon matched-paired signed rank test | 17 of 6 vs 17 of 6 | 0.4307 | 0.0771 | | |
| | | Disynaptic IPSC amplitude (PW5) | Wilcoxon matched-paired signed rank test | 24 of 8 vs 24 of 8 | 0.684 | 0.8513 | | |
| | | IPSC log ratio (PW3 vs PW5) | Mann Whitney test | 17 of 6 vs 24 of 8 | 0.4087 | 0.7042 | | |
| Fig. 3e | | E/I ratio (PW3) | Wilcoxon matched-paired signed rank test | 17 of 6 vs 17 of 6 | 0.1901 | 0.2125 | | |
| | | E/I ratio (PW5) | Wilcoxon matched-paired signed rank test | 24 of 8 vs 24 of 8 | 0.9218 | 0.1343 | | |
| | | E/I ratio log ratio (PW3 vs PW5) | Unpaired t test | 17 of 6 vs 24 of 8 | 0.3188 | 0.458 | | |
| Fig. 3h | Layer 4 input-evoked: Control vs Kir2.1 | EPSC amplitude (PW3) | Wilcoxon matched-paired signed rank test | 25 of 4 vs 25 of 4 | 0.8119 | | | |
| | | EPSC amplitude (PW5) | Wilcoxon matched-paired signed rank test | 44 of 11 vs 44 of 11 | 0.8217 | 0.0065 | 0.9667 | 0.8946 |
| | | EPSC log ratio (PW3 vs PW5) | Unpaired t test | 25 of 4 vs 44 of 11 | 0.917 | | | |
| Fig. 3i | | Disynaptic IPSC amplitude (PW3) | Wilcoxon matched-paired signed rank test | 18 of 4 vs 18 of 4 | <0.0001 | | | |
| | | Disynaptic IPSC amplitude (PW5) | Wilcoxon matched-paired signed rank test | 41 of 11 vs 41 of 11 | <0.0001 | 0.3345 | | |
| | | IPSC log ratio (PW3 vs PW5) | Unpaired t test | 18 of 4 vs 41 of 11 | 0.0506 | | | |
| Fig. 3j | | E/I ratio (PW3) | Wilcoxon matched-paired signed rank test | 18 of 4 vs 18 of 4 | 0.0019 | | | |
| | | E/I ratio (PW5) | Wilcoxon matched-paired signed rank test | 41 of 11 vs 41 of 11 | 0.0034 | 0.2346 | | |
| | | E/I ratio log ratio (PW3 vs PW5) | Unpaired t test | 18 of 4 vs 41 of 11 | 0.5136 | | | |

| | | | | | | | | |
|---|---|---|---|---|---|---|---|---|
| Fig. 3m | Layer 5 input-evoked: Control vs Kir2.1 | EPSC amplitude (PW3) | Wilcoxon matched-paired signed rank test | 36 of 6 vs 36 of 6 | 0.0006 | 0.9415 | | |
| | | EPSC amplitude (PW5) | Wilcoxon matched-paired signed rank test | 50 of 23 vs 50 of 23 | <0.0001 | 0.289 | | |
| | | EPSC log ratio (PW3 vs PW5) | Unpaired t test with Welch's correction | 36 of 6 vs 50 of 23 | <0.0001 | 0.3547 | | |
| Fig. 3n | | Disynaptic IPSC amplitude (PW3) | Wilcoxon matched-paired signed rank test | 30 of 6 vs 30 of 6 | 0.3818 | 0.212 | | |
| | | Disynaptic IPSC amplitude (PW5) | Wilcoxon matched-paired signed rank test | 45 of 22 vs 45 of 22 | 0.911 | 0.5828 | | |
| | | IPSC log ratio (PW3 vs PW5) | Unpaired t test | 30 of 6 vs 45 of 22 | 0.7828 | 0.9238 | | |
| Fig. 3o | | E/I ratio (PW3) | Wilcoxon matched-paired signed rank test | 30 of 6 vs 30 of 6 | 0.0137 | 0.3373 | | |
| | | E/I ratio (PW5) | Wilcoxon matched-paired signed rank test | 45 of 22 vs 45 of 22 | <0.0001 | 0.2711 | | |
| | | E/I ratio log ratio (PW3 vs PW5) | Unpaired t test | 30 of 6 vs 45 of 22 | 0.0246 | 0.6053 | | |
| Fig. 4c | Contralateral layer 2/3 input-evoked: Control vs Kir2.1 | EPSC amplitude (PW3) | Wilcoxon matched-paired signed rank test | 17 of 6 vs 17 of 6 | 0.0202 | 0.7351 | | |
| | | EPSC amplitude (PW5) | Wilcoxon matched-paired signed rank test | 31 of 9 vs 31 of 9 | 0.6919 | 0.5025 | | |
| | | EPSC log ratio (PW3 vs PW5) | Unpaired t test | 17 of 6 vs 31 of 9 | 0.1416 | 0.7797 | | |
| Fig. 4d | | Disynaptic IPSC amplitude (PW3) | Wilcoxon matched-paired signed rank test | 16 of 6 vs 16 of 6 | 0.0027 | 0.6232 | | |
| | | Disynaptic IPSC amplitude (PW5) | Wilcoxon matched-paired signed rank test | 31 of 9 vs 31 of 9 | 0.2556 | 0.1292 | | |
| | | IPSC log ratio (PW3 vs PW5) | Mann Whitney test | 16 of 6 vs 31 of 9 | 0.109 | 0.6646 | | |
| Fig. 4e | | E/I ratio (PW3) | Wilcoxon matched-paired signed rank test | 16 of 6 vs 16 of 6 | 0.2744 | 0.7555 | | |
| | | E/I ratio (PW5) | Wilcoxon matched-paired signed rank test | 31 of 9 vs 31 of 9 | 0.4214 | 0.3572 | | |
| | | E/I ratio log ratio (PW3 vs PW5) | Mann Whitney test | 16 of 6 vs 31 of 9 | 0.5118 | 0.58 | | |
| Fig. 4h | Contralateral layer 5 input-evoked: Control vs Kir2.1 | EPSC amplitude (PW3) | Wilcoxon matched-paired signed rank test | 25 of 10 vs 25 of 10 | <0.0001 | 0.5939 | | |
| | | EPSC amplitude (PW5) | Wilcoxon matched-paired signed rank test | 16 of 7 vs 16 of 7 | 0.0052 | 0.3658 | | |
| | | EPSC log ratio (PW3 vs PW5) | Mann Whitney test | 25 of 1o vs 16 of 7 | 0.4348 | 0.3566 | | |
| Fig. 4i | | Disynaptic IPSC amplitude (PW3) | Wilcoxon matched-paired signed rank test | 21 of 10 vs 21 of 10 | 0.5621 | 0.24 | | |
| | | Disynaptic IPSC amplitude (PW5) | Wilcoxon matched-paired signed rank test | 15 of 7 vs 15 of 7 | 0.1514 | 0.0143 | 0.0155 | 0.3851 |
| | | IPSC log ratio (PW3 vs PW5) | Unpaired t test | 21 of 10 vs 15 of 7 | 0.4255 | 0.4649 | | |
| Fig. 4j | | E/I ratio (PW3) | Wilcoxon matched-paired signed rank test | 21 of 10 vs 21 of 10 | 0.0033 | 0.1897 | | |
| | | E/I ratio (PW5) | Wilcoxon matched-paired signed rank test | 15 of 7 vs 15 of 7 | 0.0084 | 0.2357 | | |
| | | E/I ratio log ratio (PW3 vs PW5) | Mann Whitney test | 21 of 10 vs 15 of 7 | 0.4851 | 0.9756 | | |
| Fig. 5d | Quantal AMPA-EPSCs: Control vs Kir2.1 | Total evoked EPSC amplitude | Wilcoxon matched-pairs signed rank test | 24 of 12 vs 24 of 12 | <0.0001 | 0.9149 | | |
| Fig. 5e | | Quantal AMPA-EPSC amplitude | Wilcoxon matched-pairs signed rank test | 24 of 12 vs 24 of 12 | 0.0001 | 0.7588 | | |
| Fig. 5f | | Log ratio of EPSC and qEPSC | Wilcoxon matched-pairs signed rank test | 24 of 12 vs 24 of 12 | <0.0001 | 0.6376 | | |
| Fig. 5g | | Quantal AMPA-EPSC frequency | Wilcoxon matched-pairs signed rank test | 24 of 12 vs 24 of 12 | <0.0001 | 0.6137 | | |
| Fig. 5h | | Quantal AMPA-EPSC decay constant | Wilcoxon matched-pairs signed rank test | 24 of 12 vs 24 of 12 | 0.5838 | 0.5558 | | |
| Fig. 5l | Layer 5 input-mediated synapse labelling and reconstruction: Control vs Kir2.1 | Dendrite length | Wilcoxon matched-pairs signed rank test | 7 of 4 vs 7 of 4 | 0.2188 | >0.9999 | | |
| Fig. 5m | | Total spine conuts | Wilcoxon matched-pairs signed rank test | 7 of 4 vs 7 of 4 | >0.9999 | 0.226 | | |
| Fig. 5n | | Syp puncta on dendrtites and spines | Wilcoxon matched-pairs signed rank test | 7 of 4 vs 7 of 4 | 0.4688 | 0.3506 | | |
| Fig. 5o | | Syp-spine counts | Wilcoxon matched-pairs signed rank test | 7 of 4 vs 7 of 4 | 0.9375 | 0.2833 | | |
| Fig. 5p | | Syp-spine classification | Two-way RM ANOVA, Šídák's multiple comparisons test | 7 of 4 vs 7 of 4 | 0.0029 (Mushroom), 0.0064 (Thin), 0.6262 (Stubby), 0.8199 (Filopodia) | 0.523 | | |
| Ext Fig. 9a | | Total spine density | Wilcoxon matched-pairs signed rank test | 7 of 4 vs 7 of 4 | 0.4588 | 0.4069 | | |
| Ext Fig. 9b | | Syp puncta density | Wilcoxon matched-pairs signed rank test | 7 of 4 vs 7 of 4 | 0.8125 | 0.5668 | | |
| Ext Fig. 9c | | Syp-spine density | Wilcoxon matched-pairs signed rank test | 7 of 4 vs 7 of 4 | 0.4688 | 0.4569 | | |
| Ext Fig. 9d | | Overall spine classification | Two-way RM ANOVA, Šídák's multiple comparisons test | 7 of 4 vs 7 of 4 | 0.1403 (Mushroom), 0.0198 (Thin), >0.9999 (Stubby), 0.8474 (Filopodia) | >0.9999 | | |
| Fig. 5s | Layer 5 presynaptic release probability: Control vs Kir2.1 | NMDA-EPSC block rate with MK-801 | Two-way RM ANOVA, , Šídák's multiple comparisons test | 8 of 3 vs 8 of 3 | 0.9653 | 0.3926 | | |
| Fig. 6c | Layer 5 input-mediated AMPA/NMDA ratio: Control vs Kir2.1 | AMPA-EPSC amplitude | Wilcoxon matched-pairs signed rank test | 29 of 10 vs 29 of 10 | <0.0001 | 0.2465 | | |
| Fig. 6d | | NMDA-EPSC amplitude | Wilcoxon matched-pairs signed rank test | 29 of 10 vs 29 of 10 | <0.0001 | 0.0184 | <0.0001 | 0.0088 |
| Fig. 6e | | A/N ratio | Wilcoxon matched-pairs signed rank test | 29 of 10 vs 29 of 10 | <0.0001 | 0.349 | | |
| Fig. 6h | GluA1CT perturbation | EPSC amplitude | One-way ANOVA Friedman test, Dunn's multiple comparisons test | 18 of 10 vs 18 of 10 | <0.0001 (Control vs Kir2.1), >0.9999 (Kir2.1 vs Kir2.1+GluA1CT), <0.0001 (Control vs Kir2.1+GluA1CT) | 0.2172 | | |
| Fig. 6k | GluA2CT perturbation | EPSC amplitude | One-way ANOVA Friedman test, Dunn's multiple comparisons test | 29 of 20 vs 29 of 20 | <0.0001 (Control vs Kir2.1), 0.0031 (Kir2.1 vs Kir2.1+GluA2CT), 0.0031 (Control vs Kir2.1+GluA2CT) | 0.6458 | | |

| | | | | | | | | |
|---|---|---|---|---|---|---|---|---|
| Ext Fig. 1f | Intrinsic excitability: Control vs Kir2.1 | Resting potential | Mann Whitney test | PW2: 7 of 2 vs 15 of 2; PW3: 15 of 3 vs 23 of 3; PW5: 17 of 3 vs 17 of 3 | <0.0001 (PW2), 0.7013 (PW3), 0.2451 (PW5) | 0.0220 (PW3) | 0.4631 (PW3) | 0.7044 (PW3) |
| Ext Fig. 1g | | Input resistance | Mann Whitney test | PW2: 7 of 2 vs 15 of 2; PW3: 15 of 3 vs 23 of 3; PW5: 17 of 3 vs 17 of 3 | <0.0001 (PW2), <0.0001 (PW3), <0.0001 (PW5) | 0.4203 (PW3) | | |
| Ext Fig. 1h | | Rheobase current | Mann Whitney test | PW2: 7 of 2 vs 15 of 2; PW3: 15 of 3 vs 23 of 3; PW5: 17 of 3 vs 17 of 3 | <0.0001 (PW2), <0.0001 (PW3), <0.0001 (PW5) | 0.2493 (PW3) | | |
| Ext Fig. 1k | Intrinsic excitability: Control vs Kir2.1Mut | Resting potential | Mann Whitney test | PW2: 5 of 1 vs 7 of 1; PW3: 5 of 1 vs 5 of 1; PW5: 24 of 5 vs 22 of 5 | 0.5303 (PW2), 0.5476 (PW3), 0.0004 (PW5) | 0.0290 (PW5) | 0.0685 (PW5) | 0.0011 (PW5) |
| Ext Fig. 1l | | Input resistance | Mann Whitney test | PW2: 5 of 1 vs 7 of 1; PW3: 5 of 1 vs 5 of 1; PW5: 24 of 5 vs 22 of 5 | 0.2020 (PW2), 0.0079 (PW3), <0.0001 (PW5) | 0.0354 (PW5) | 0.0025 (PW5) | <0.0001 (PW5) |
| Ext Fig. 1m | | Rheobase current | Mann Whitney test | PW2: 5 of 1 vs 7 of 1; PW3: 5 of 1 vs 5 of 1; PW5: 24 of 5 vs 22 of 5 | 0.2361 (PW2), >0.9999 (PW3), 0.0488 (PW5) | 0.0007 (PW5) | 0.1003 (PW5) | 0.1240 (PW5) |
| Ext Fig. 2c | Intrinsic excitability: Non-electroporated vs Untransfected vs Kir2.1 | Resting potential | One way ANOVA Kruskal-Wallis test, Dunn's multiple comparisons test | 25 of 7 vs 25 of 7 vs 19 of 7 | >0.9999 (Non-electroporated vs Untransfected), >0.9999 (Untransfected vs Kir2.1) | 0.2067 | | |
| Ext Fig. 2d | | Input resistance | One way ANOVA Kruskal-Wallis test, Dunn's multiple comparisons test | 25 of 7 vs 25 of 7 vs 19 of 7 | >0.9999 (Non-electroporated vs Untransfected), <0.0001 (Untransfected vs Kir2.1) | 0.0666 | | |
| Ext Fig. 2e | | Rheobase current | One way ANOVA Kruskal-Wallis test, Dunn's multiple comparisons test | 25 of 7 vs 25 of 7 vs 19 of 7 | >0.9999 (Non-electroporated vs Untransfected), <0.0001 (Untransfected vs Kir2.1) | 0.9087 | | |
| Ext Fig. 2g | Spontaneous EPSCs and IPSCs: Non-electroporated vs Untransfected | sEPSC frequency | Mann Whitney test | 45 of 7 vs 45 of 7 | 0.734 | 0.0144 | 0.1198 | 0.267 |
| Ext Fig. 2h | | sEPSC amplitude | Mann Whitney test | 45 of 7 vs 45 of 7 | 0.756 | 0.3727 | | |
| Ext Fig. 2i | | sIPSC frequency | Mann Whitney test | 45 of 7 vs 45 of 7 | 0.081 | 0.1247 | | |
| Ext Fig. 2j | | sIPSC amplitude | Mann Whitney test | 45 of 7 vs 45 of 7 | 0.1977 | 0.7532 | | |
| Ext Fig. 3b | Spontaneous EPSCs: Control vs Kir2.1 | sEPSC frequency (PW3) | Mann Whitney test | 141 of 33 vs 141 vs 33 | <0.0001 | 0.508 | | |
| Ext Fig. 3c | | sEPSC amplitude (PW3) | Mann Whitney test | 141 of 33 vs 141 vs 33 | <0.0001 | 0.5843 | | |
| Ext Fig. 3d | | sEPSC frequency (PW5) | Mann Whitney test | 162 of 39 vs 162 of 39 | <0.0001 | 0.0067 | <0.0001 | <0.0001 |
| Ext Fig. 3e | | sEPSC amplitude (PW5) | Mann Whitney test | 162 of 39 vs 162 of 39 | <0.0001 | 0.0134 | <0.0001 | <0.0001 |
| Ext Fig. 3g | Spontaneous IPSCs: Control vs Kir2.1 | sIPSC frequency (PW3) | Mann Whitney test | 141 of 33 vs 141 vs 33 | <0.0001 | 0.2016 | | |
| Ext Fig. 3h | | sIPSC amplitude (PW3) | Mann Whitney test | 141 of 33 vs 141 vs 33 | <0.0001 | 0.7773 | | |
| Ext Fig. 3i | | sIPSC frequency (PW5) | Unpaired t test with Welch's correction | 162 of 39 vs 162 of 39 | 0.1241 | 0.1867 | | |
| Ext Fig. 3j | | sIPSC amplitude (PW5) | Mann Whitney test | 162 of 39 vs 162 of 39 | <0.0001 | 0.8044 | | |
| Ext Fig. 5c | Thalamic input-evoked: Control vs Kir2.1 | EPSC amplitude (PW3) | Wilcoxon matched-paired signed rank test | 25 of 6 vs 25 of 6 | 0.751 | 0.3596 | | |
| | | EPSC amplitude (PW5) | Wilcoxon matched-paired signed rank test | 29 of 7 vs 29 of 7 | 0.8314 | 0.1209 | | |
| | | EPSC log ratio (PW3 vs PW5) | Unpaired t test | 25 of 6 vs 29 of 7 | 0.913 | 0.4592 | | |
| Ext Fig. 5d | | Disynaptic IPSC amplitude (PW3) | Wilcoxon matched-paired signed rank test | 22 of 6 vs 22 of 6 | 0.0011 | 0.743 | | |
| | | Disynaptic IPSC amplitude (PW5) | Wilcoxon matched-paired signed rank test | 28 of 7 vs 28 of 7 | <0.0001 | 0.8044 | | |
| | | IPSC log ratio (PW3 vs PW5) | Unpaired t test | 22 of 6 vs 28 of 7 | 0.8769 | 0.5735 | | |
| Ext Fig. 5e | | E/I ratio (PW3) | Wilcoxon matched-paired signed rank test | 22 of 6 vs 22 of 6 | 0.0004 | 0.23 | | |
| | | E/I ratio (PW5) | Wilcoxon matched-paired signed rank test | 28 of 7 vs 28 of 7 | 0.0012 | 0.6249 | | |
| | | E/I ratio log ratio (PW3 vs PW5) | Unpaired t test | 22 of 6 vs 28 of 7 | 0.988 | 0.2606 | | |
| Ext Fig. 5h | Layer 6 input-evoked: Control vs Kir2.1 | EPSC amplitude (PW3) | Wilcoxon matched-paired signed rank test | 35 of 13 vs 35 of 13 | 0.0001 | 0.0981 | | |
| | | EPSC amplitude (PW5) | Wilcoxon matched-paired signed rank test | 43 of 14 vs 43 of 14 | <0.0001 | 0.2523 | | |
| | | EPSC log ratio (PW3 vs PW5) | Unpaired t test | 35 of 13 vs 43 of 14 | 0.0107 | 0.7612 | | |
| Ext Fig. 5i | | Disynaptic IPSC amplitude (PW3) | Wilcoxon matched-paired signed rank test | 35 of 13 vs 35 of 13 | <0.0001 | 0.0373 | 0.0001 | 0.0907 |
| | | Disynaptic IPSC amplitude (PW5) | Wilcoxon matched-paired signed rank test | 43 of 14 vs 43 of 14 | 0.1922 | 0.9764 | | |
| | | IPSC log ratio (PW3 vs PW5) | Mann Whitney test | 35 of 13 vs 43 of 14 | <0.0001 | 0.4309 | | |
| Ext Fig. 5j | | E/I ratio (PW3) | Wilcoxon matched-paired signed rank test | 35 of 13 vs 35 of 13 | <0.0001 | 0.9631 | | |
| | | E/I ratio (PW5) | Wilcoxon matched-paired signed rank test | 43 of 14 vs 43 of 14 | <0.0001 | 0.0141 | 0.0288 | <0.0001 |
| | | E/I ratio log ratio (PW3 vs PW5) | Unpaired t test | 35 of 13 vs 43 of 14 | 0.5589 | 0.7773 | | |
| Ext Fig. 6c | Layer 5 IT input-evoked: Control vs Kir2.1 | EPSC amplitude (PW3) | Wilcoxon matched-paired signed rank test | 10 of 2 vs 10 of 2 | 0.002 | 0.3022 | | |
| | | EPSC amplitude (PW5) | Wilcoxon matched-paired signed rank test | 20 of 5 vs 20 of 5 | <0.0001 | 0.4414 | | |
| | | EPSC log ratio (PW3 vs PW5) | Unpaired t test | 10 of 2 vs 20 of 5 | 0.729 | 0.0011 | 0.8246 | 0.8233 |
| Ext Fig. 6d | | Disynaptic IPSC amplitude (PW3) | Wilcoxon matched-paired signed rank test | 9 of 2 vs 9 of 2 | 0.0273 | 0.7195 | | |
| | | Disynaptic IPSC amplitude (PW5) | Wilcoxon matched-paired signed rank test | 20 of 5 vs 20 of 5 | 0.2162 | 0.229 | | |
| | | IPSC log ratio (PW3 vs PW5) | Unpaired t test | 9 of 2 vs 20 of 5 | 0.0127 | 0.9623 | | |

| | | | | | | | | |
|---|---|---|---|---|---|---|---|---|
| Ext Fig. 6e | | E/I ratio (PW3) | Wilcoxon matched-paired signed rank test | 9 of 2 vs 9 of 2 | 0.0039 | 0.3957 | | |
| | | E/I ratio (PW5) | Wilcoxon matched-paired signed rank test | 20 of 5 vs 20 of 5 | <0.0001 | 0.2527 | | |
| | | E/I ratio log ratio (PW3 vs PW5) | Unpaired t test | 9 of 2 vs 20 of 5 | 0.0205 | 0.0286 | 0.0623 | 0.0291 |
| Ext Fig. 6h | Layer 5 ET input-evoked: Control vs Kir2.1 | EPSC amplitude (PW3) | Wilcoxon matched-paired signed rank test | 7 of 2 vs 7 of 2 | 0.0156 | 0.0932 | | |
| | | EPSC amplitude (PW5) | Wilcoxon matched-paired signed rank test | 13 of 3 vs 13 of 3 | 0.0002 | 0.5874 | | |
| | | EPSC log ratio (PW3 vs PW5) | Mann Whitney test | 7 of 2 vs 13 of 3 | 0.3114 | 0.1762 | | |
| Ext Fig. 6i | | Disynaptic IPSC amplitude (PW3) | Wilcoxon matched-paired signed rank test | 7 of 2 vs 7 of 2 | 0.5781 | 0.7957 | | |
| | | Disynaptic IPSC amplitude (PW5) | Wilcoxon matched-paired signed rank test | 11 of 3 vs 11 of 3 | 0.6377 | 0.34 | | |
| | | IPSC log ratio (PW3 vs PW5) | Mann Whitney test | 7 of 2 vs 11 of 3 | 0.7242 | 0.8796 | | |
| Ext Fig. 6j | | E/I ratio (PW3) | Wilcoxon matched-paired signed rank test | 7 of 2 vs 7 of 2 | 0.0312 | 0.0374 | 0.162 | 0.0097 |
| | | E/I ratio (PW5) | Wilcoxon matched-paired signed rank test | 11 of 3 vs 11 of 3 | 0.001 | 0.3492 | | |
| | | E/I ratio log ratio (PW3 vs PW5) | Mann Whitney test | 7 of 2 vs 11 of 3 | 0.4252 | 0.5194 | | |
| Ext Fig. 7c | Layer5 input-evoked: Control vs Kir2.1Mut | EPSC amplitude (PW3) | Wilcoxon matched-paired signed rank test | 24 of 4 vs 24 of 4 | 0.4732 | | | |
| | | EPSC amplitude (PW5) | Wilcoxon matched-paired signed rank test | 19 of 5 vs 19 of 5 | 0.6226 | 0.2217 | | |
| | | EPSC log ratio (PW3 vs PW5) | Unpaired t test | 24 of 4 vs 19 of 5 | 0.6322 | | | |
| Ext Fig. 7d | | Disynaptic IPSC amplitude (PW3) | Wilcoxon matched-paired signed rank test | 23 of 4 vs 23 of 4 | 0.0112 | | | |
| | | Disynaptic IPSC amplitude (PW5) | Wilcoxon matched-paired signed rank test | 17 of 5 vs 17 of 5 | 0.7819 | 0.8447 | | |
| | | IPSC log ratio (PW3 vs PW5) | Unpaired t test | 23 of 4 vs 17 of 5 | 0.1012 | | | |
| Ext Fig. 7e | | E/I ratio (PW3) | Wilcoxon matched-paired signed rank test | 23 of 4 vs 23 of 4 | 0.0179 | | | |
| | | E/I ratio (PW5) | Wilcoxon matched-paired signed rank test | 17 of 5 vs 17 of 5 | 0.7119 | 0.658 | | |
| | | E/I ratio log ratio (PW3 vs PW5) | Unpaired t test | 23 of 4 vs 17 of 5 | 0.0898 | | | |
| Ext Fig. 7h | Layer 6 input-evoked: Control vs Kir2.1Mut | EPSC amplitude (PW3) | Wilcoxon matched-paired signed rank test | 21 of 3 vs 21 of 3 | 0.1111 | 0.0315 | 0.6037 | 0.0012 |
| | | EPSC amplitude (PW5) | Wilcoxon matched-paired signed rank test | 18 of 4 vs 18 of 4 | 0.9661 | 0.5716 | | |
| | | EPSC log ratio (PW3 vs PW5) | Unpaired t test | 21 of 3 vs 18 of 4 | 0.4087 | 0.1724 | | |
| Ext Fig. 7i | | Disynaptic IPSC amplitude (PW3) | Wilcoxon matched-paired signed rank test | 21 of 3 vs 21 of 3 | 0.0033 | 0.3436 | | |
| | | Disynaptic IPSC amplitude (PW5) | Wilcoxon matched-paired signed rank test | 18 of 4 vs 18 of 4 | 0.2121 | 0.8255 | | |
| | | IPSC log ratio (PW3 vs PW5) | Unpaired t test | 21 of 3 vs 18 of 4 | 0.2317 | 0.863 | | |
| Ext Fig. 7j | | E/I ratio (PW3) | Wilcoxon matched-paired signed rank test | 21 of 3 vs 21 of 3 | 0.0239 | 0.5715 | | |
| | | E/I ratio (PW5) | Wilcoxon matched-paired signed rank test | 18 of 4 vs 18 of 4 | 0.3038 | 0.6739 | | |
| | | E/I ratio log ratio (PW3 vs PW5) | Unpaired t test | 21 of 3 vs 18 of 4 | 0.1654 | 0.3112 | | |
| Ext Fig. 7m | Contralateral layer 5 input-evoked: Control vs Kir2.1Mut | EPSC amplitude (PW3) | Wilcoxon matched-paired signed rank test | 16 of 2 vs 16 of 2 | 0.0739 | 0.6238 | | |
| | | EPSC amplitude (PW5) | Wilcoxon matched-paired signed rank test | 20 of 4 vs 20 of 4 | 0.0759 | | | |
| | | EPSC log ratio (PW3 vs PW5) | Mann Whitney test | 16 of 2 vs 20 of 4 | 0.7176 | | | |
| Ext Fig. 7n | | Disynaptic IPSC amplitude (PW3) | Wilcoxon matched-paired signed rank test | 16 of 2 vs 16 of 2 | 0.782 | 0.1823 | | |
| | | Disynaptic IPSC amplitude (PW5) | Wilcoxon matched-paired signed rank test | 20 of 4 vs 20 of 4 | 0.0266 | | | |
| | | IPSC log ratio (PW3 vs PW5) | Mann Whitney test | 16 of 2 vs 20 of 4 | 0.0949 | | | |
| Ext Fig. 7o | | E/I ratio (PW3) | Wilcoxon matched-paired signed rank test | 16 of 2 vs 16 of 2 | 0.1439 | 0.4967 | | |
| | | E/I ratio (PW5) | Wilcoxon matched-paired signed rank test | 20 of 4 vs 20 of 4 | 0.0042 | | | |
| | | E/I ratio log ratio (PW3 vs PW5) | Unpaired t test | 16 of 2 vs 20 of 4 | 0.1103 | | | |
| Ext Fig. 8c | Layer 5 presynaptic vesicle content: Control vs Kir2.1 | Baseline AMPA-EPSC amplitude | Wilcoxon matched-paired signed rank test | 11 of 5 vs 11 of 5 | 0.001 | 0.1149 | | |
| Ext Fig. 8d | | AMPA-EPSC amplitude with γ-DGG | Wilcoxon matched-paired signed rank test | 11 of 5 vs 11 of 5 | 0.001 | 0.1111 | | |
| Ext Fig. 8e | | Peak current reduction percentage | Wilcoxon matched-paired signed rank test | 11 of 5 vs 11 of 5 | 0.123 | 0.2434 | | |
| Ext Fig. 8f | | Baseline AMPA-EPSQ | Wilcoxon matched-paired signed rank test | 11 of 5 vs 11 of 5 | 0.001 | 0.0054 | <0.0001 | 0.0004 |
| Ext Fig. 8g | | AMPA-EPSQ with γ-DGG | Wilcoxon matched-paired signed rank test | 11 of 5 vs 11 of 5 | 0.001 | 0.0072 | <0.0001 | <0.0001 |
| Ext Fig. 8h | | Charge reduction percentage | Wilcoxon matched-paired signed rank test | 11 of 5 vs 11 of 5 | 0.2402 | 0.3917 | | |
| Ext Fig. 10b | Intrinsic excitability of Kir2.1 co-expression with GluA1CT or GluA2CT | Resting potential | One way ANOVA Kruskal-Wallis test, Dunn's multiple comparisons test | 13 of 6 (Control) vs 10 of 5 (Kir2.1) vs 7 of 2 (Kir2.1+GluA1CT) vs 9 of 4 (Kir2.1+GluA2CT) | 0.1844 (Control vs Kir2.1), 0.7312 (Control vs Kir2.1+GluA1CT), >0.9999 (Kir2.1+GluA2CT) | 0.4034 | | |
| Ext Fig. 10c | | Input resistance | One way ANOVA Kruskal-Wallis test, Dunn's multiple comparisons test | | 0.0002 (Control vs Kir2.1), 0.0057 (Control vs Kir2.1+GluA1CT), 0.0006 (Kir2.1+GluA2CT), >0.9999 (Kir2.1 vs Kir2.1+GluA1CT), >0.9999 (Kir2.1 vs Kir2.1+GluA2CT) | 0.34 | | |
| Ext Fig. 10d | | Rheobase current | One way ANOVA Kruskal-Wallis test, Dunn's multiple comparisons test | | 0.0002 (Control vs Kir2.1), 0.0022 (Control vs Kir2.1+GluA1CT), 0.0002 (Kir2.1+GluA2CT), >0.9999 (Kir2.1 vs Kir2.1+GluA1CT), >0.9999 (Kir2.1 vs Kir2.1+GluA2CT) | 0.1622 | | |

| | | | | | | |
|---|---|---|---|---|---|---|
| Ext Fig. 10g | The effect of GluA1CT on control and Kir2.1 neurons | EPSC amplitude (Control vs GluA1CT) | Wilcoxon matched-paired signed rank test | 18 of 3 vs 18 of 3 | 0.0898 | 0.228 |
| Ext Fig. 10h | | Log ratio (GluA1CT in Control vs Kir2.1) | Mann Whitney test | 18 of 3 vs 18 of 10 | 0.4619 | 0.7628 |
| Ext Fig. 10k | The effect of GluA2CT on control and Kir2.1 neurons | EPSC amplitude (Control vs GluA2CT) | Wilcoxon matched-paired signed rank test | 14 of 3 vs 14 of 3 | 0.0419 | 0.2136 |
| Ext Fig. 10l | | Log ratio (GluA2CT in Control vs Kir2.1) | Unpaired t test with Welch's correction | 14 of 3 vs 29 of 20 | 0.0196 | 0.245 |
