## Supplementary figures and images for "Synaptic input-selective homeostasis safeguards developing cortical neurons toward setpoint activity"

### Extended data F1_Intrinsic excitability (Kir2.1 or Kir2.1Mut) +Transfection rate 20260529.tif

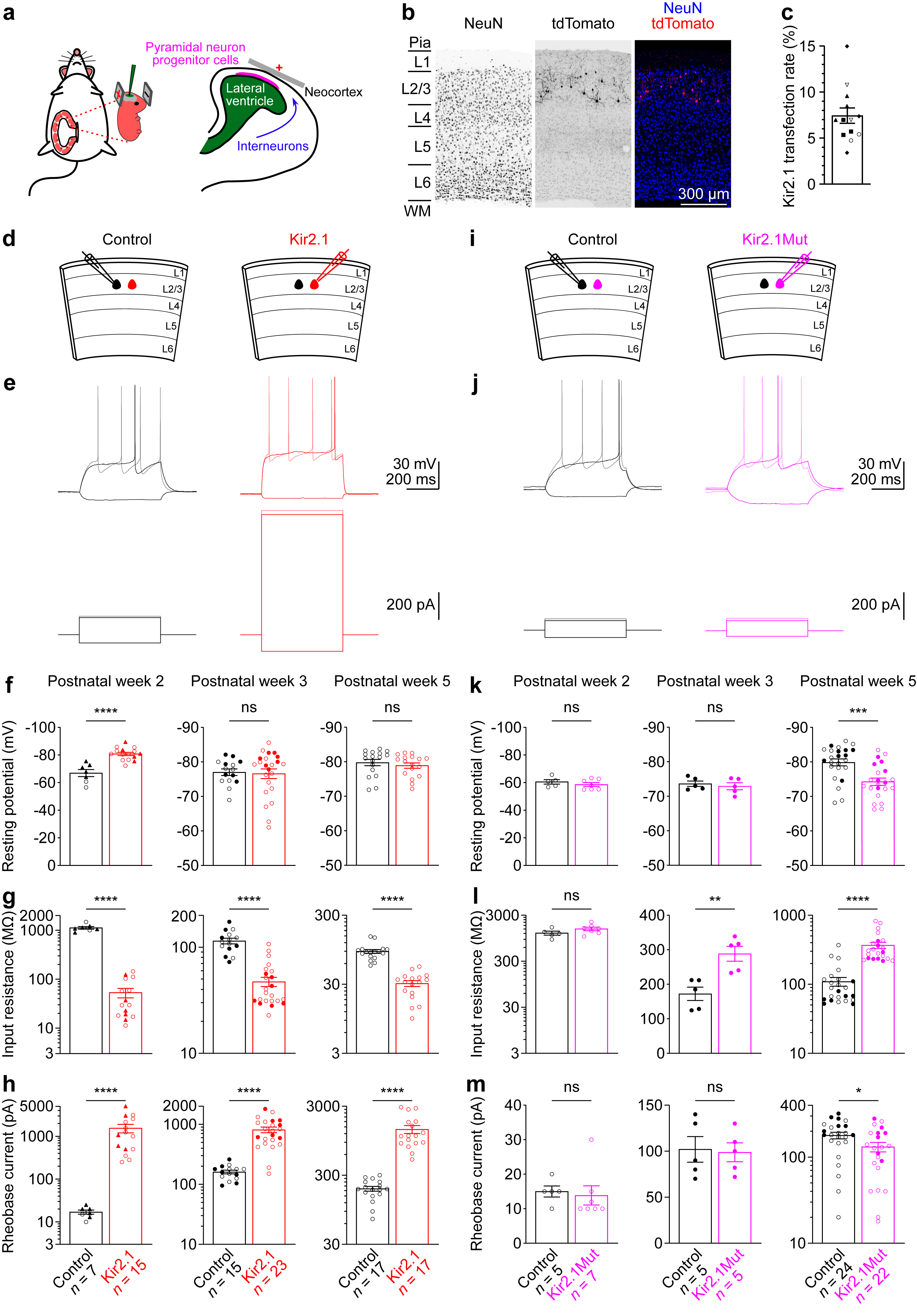

### Extended data F2_Blank control vs Untransfected intrinsic excitability + sEPSC_sIPSC P28-35 20260427.tif

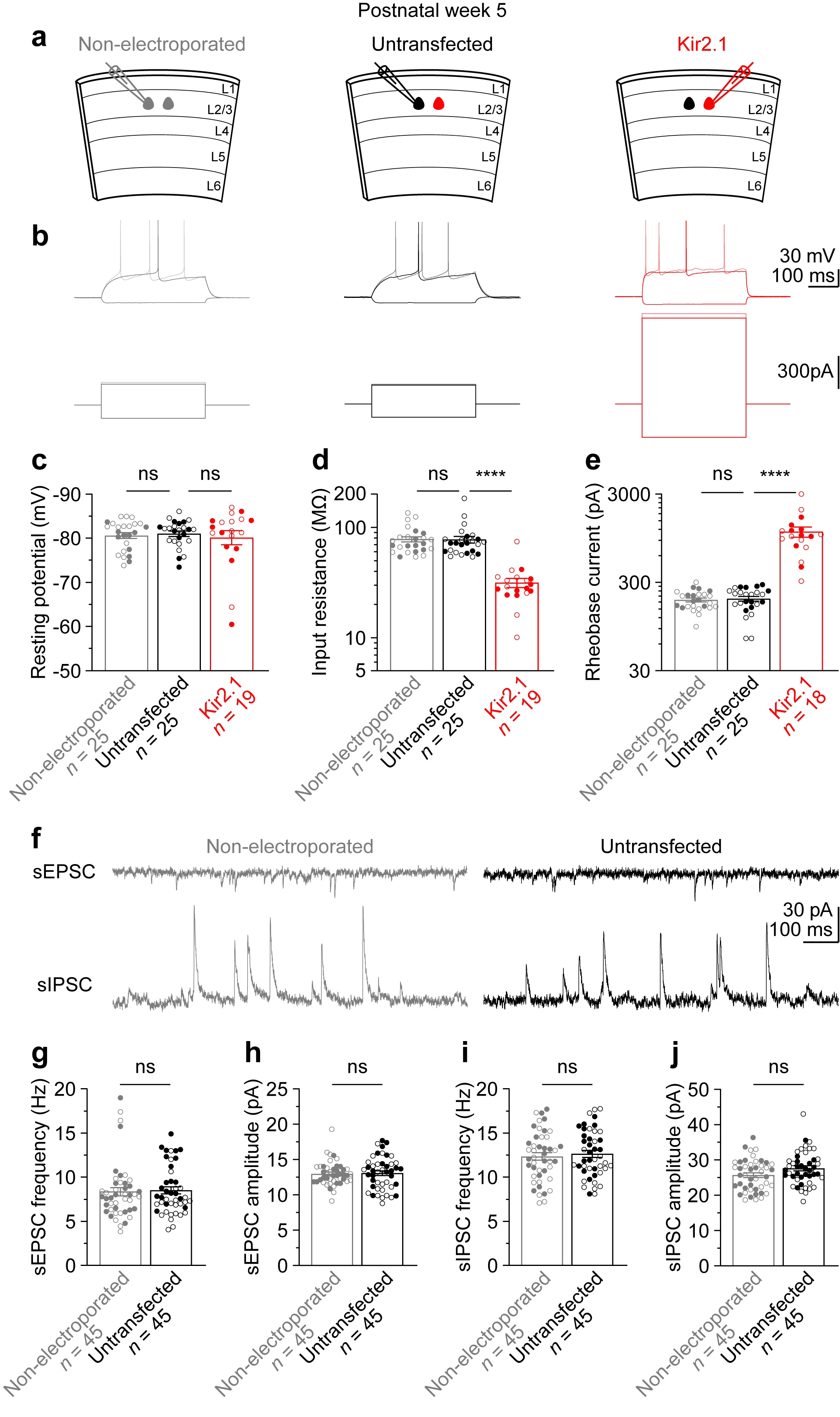

### Extended data F3_sEPSC+sIPSC 20260415.tif

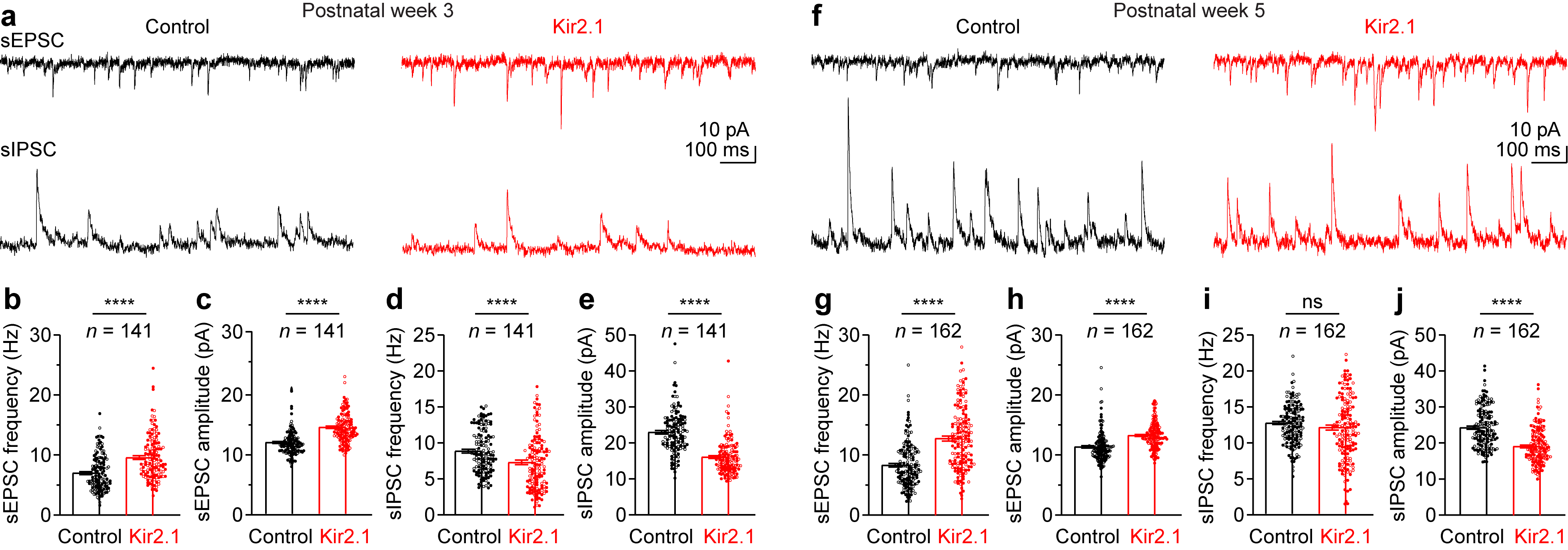

### Extended data F4_Non-scaling of mEPSCs and mIPSCs (best fit equation) 20260424.tif

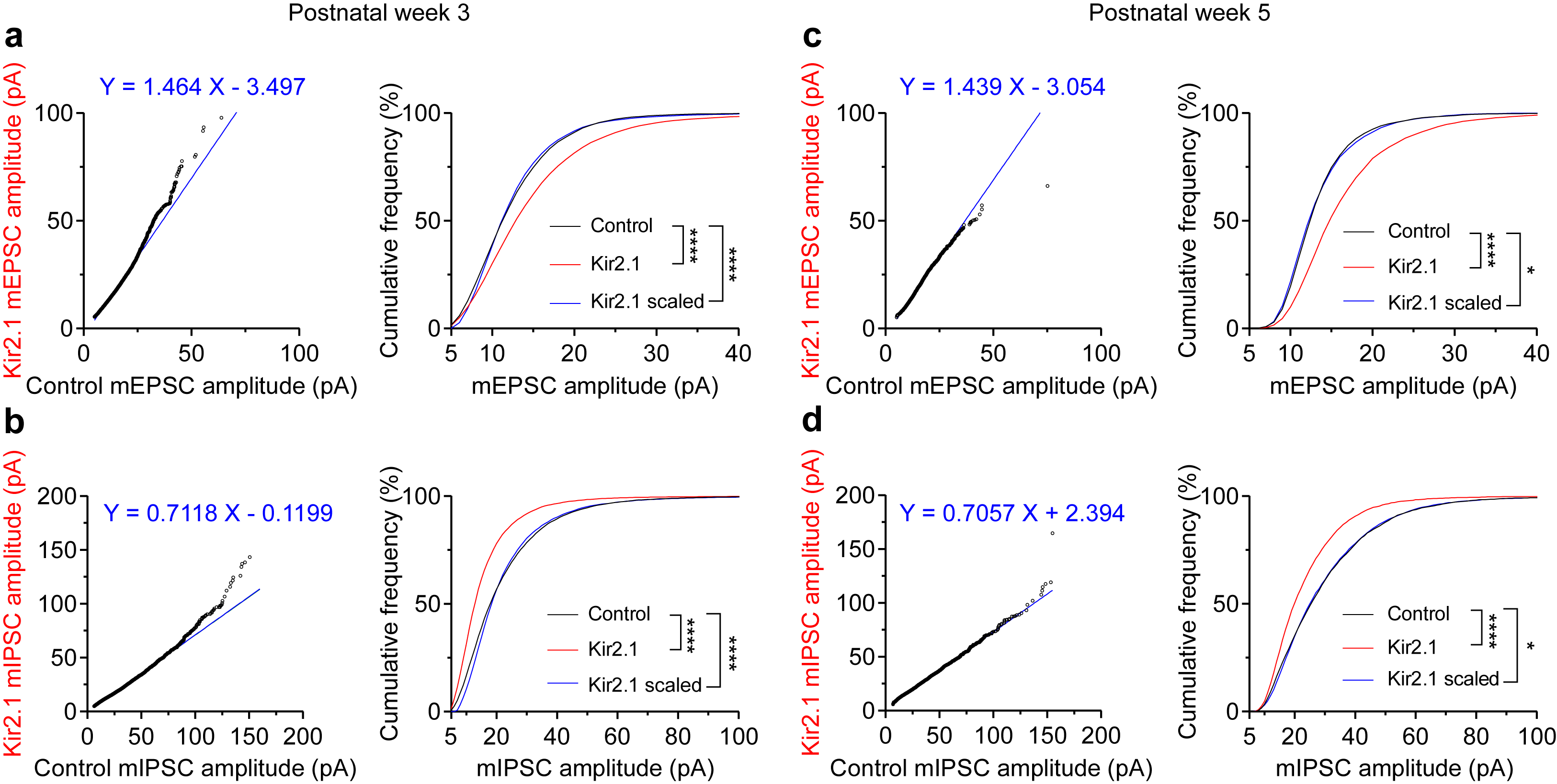

### Extended data F5_Thalamic and Layer 6 input EPSC+IPSC+EI ratio 20260602.tif

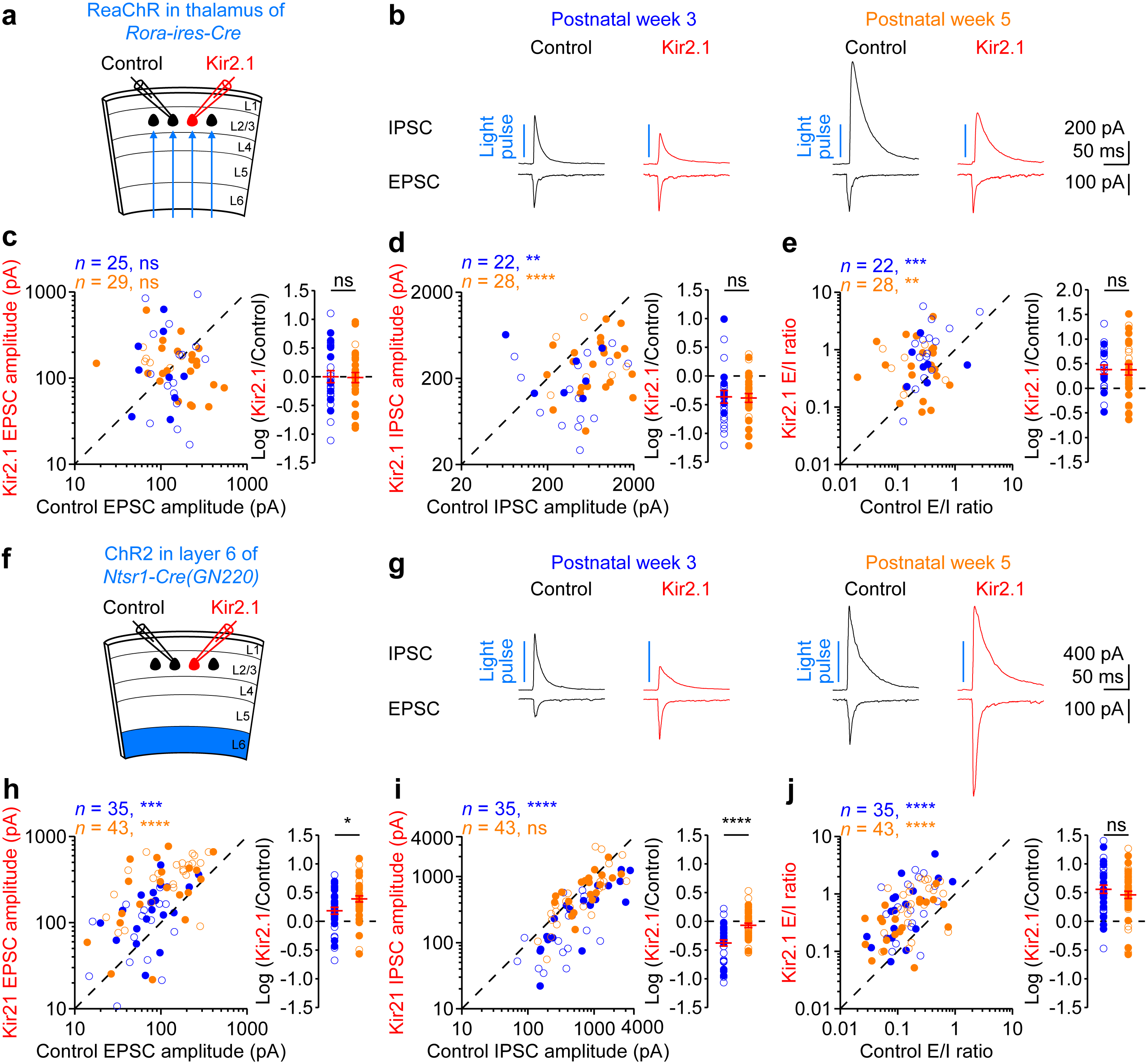

### Extended data F6_Layer 5 IT ET inputs 20260630.tif

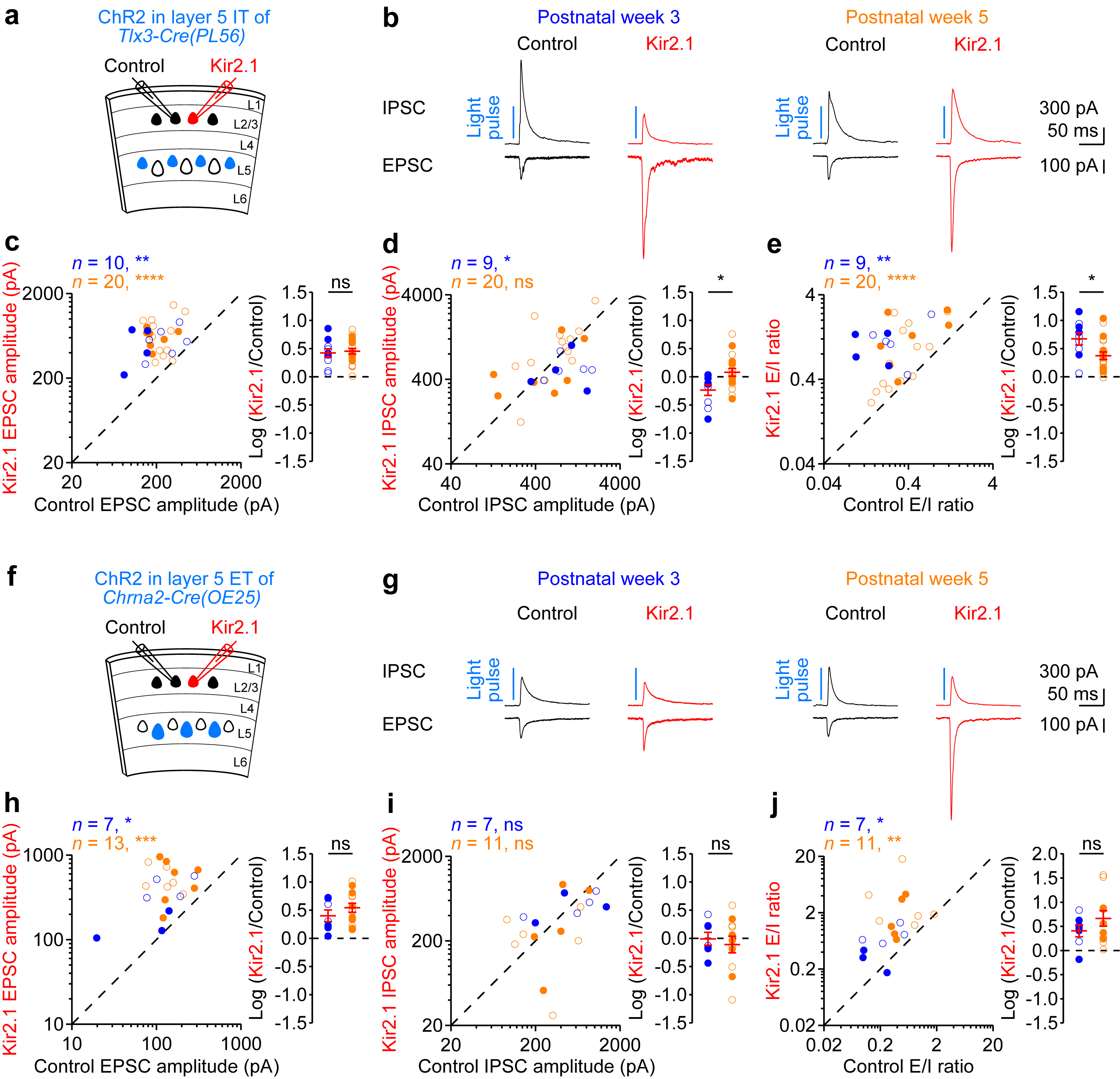

### Extended data F7_Layer5_6_contralateral L5 inputs to Kir2.1Mut EPSC 20260602.tif

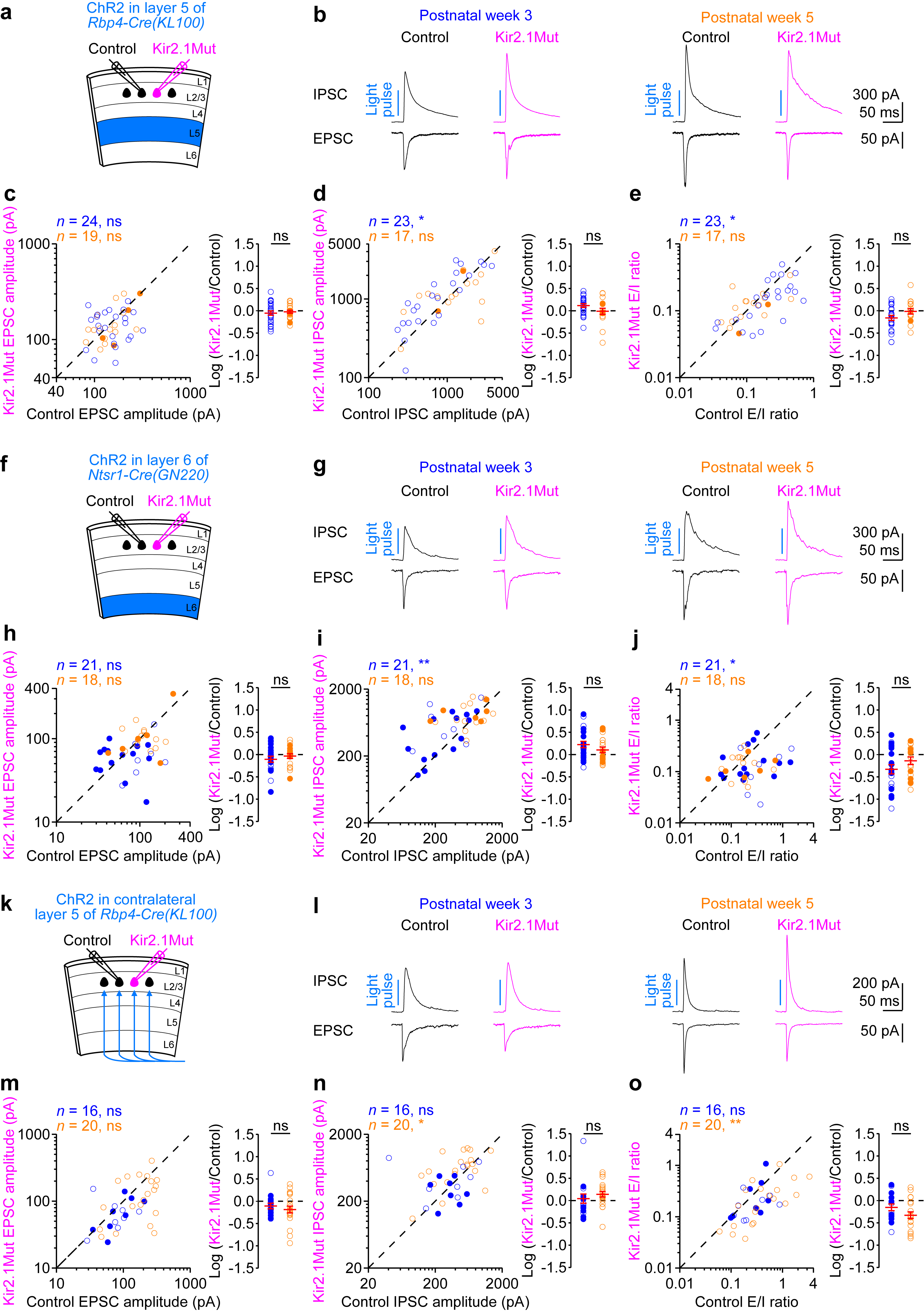

### Extended data F8 DGG_20260424.tif

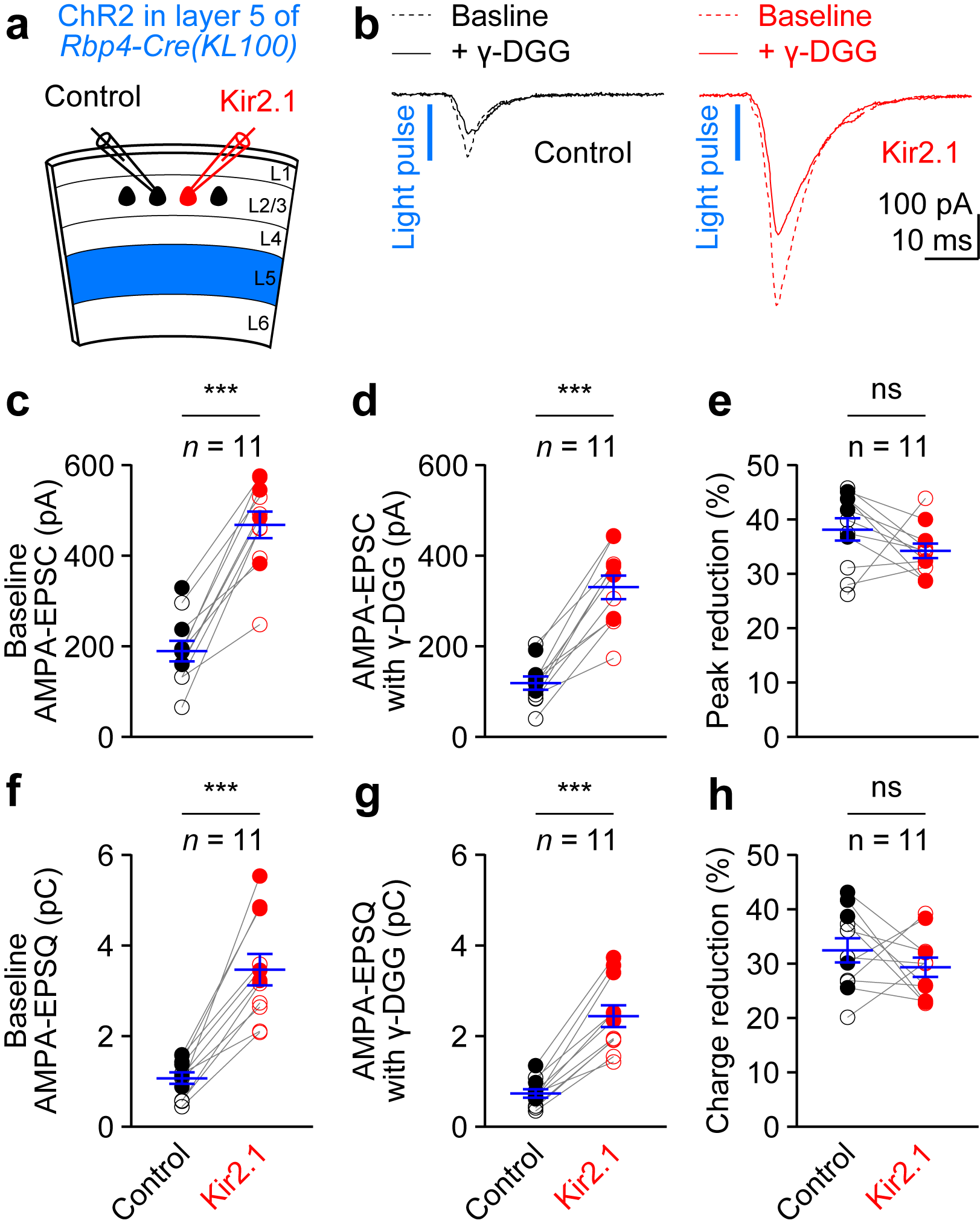

### Extended data F9 Quantification on all dendritic spines_20260424.tif

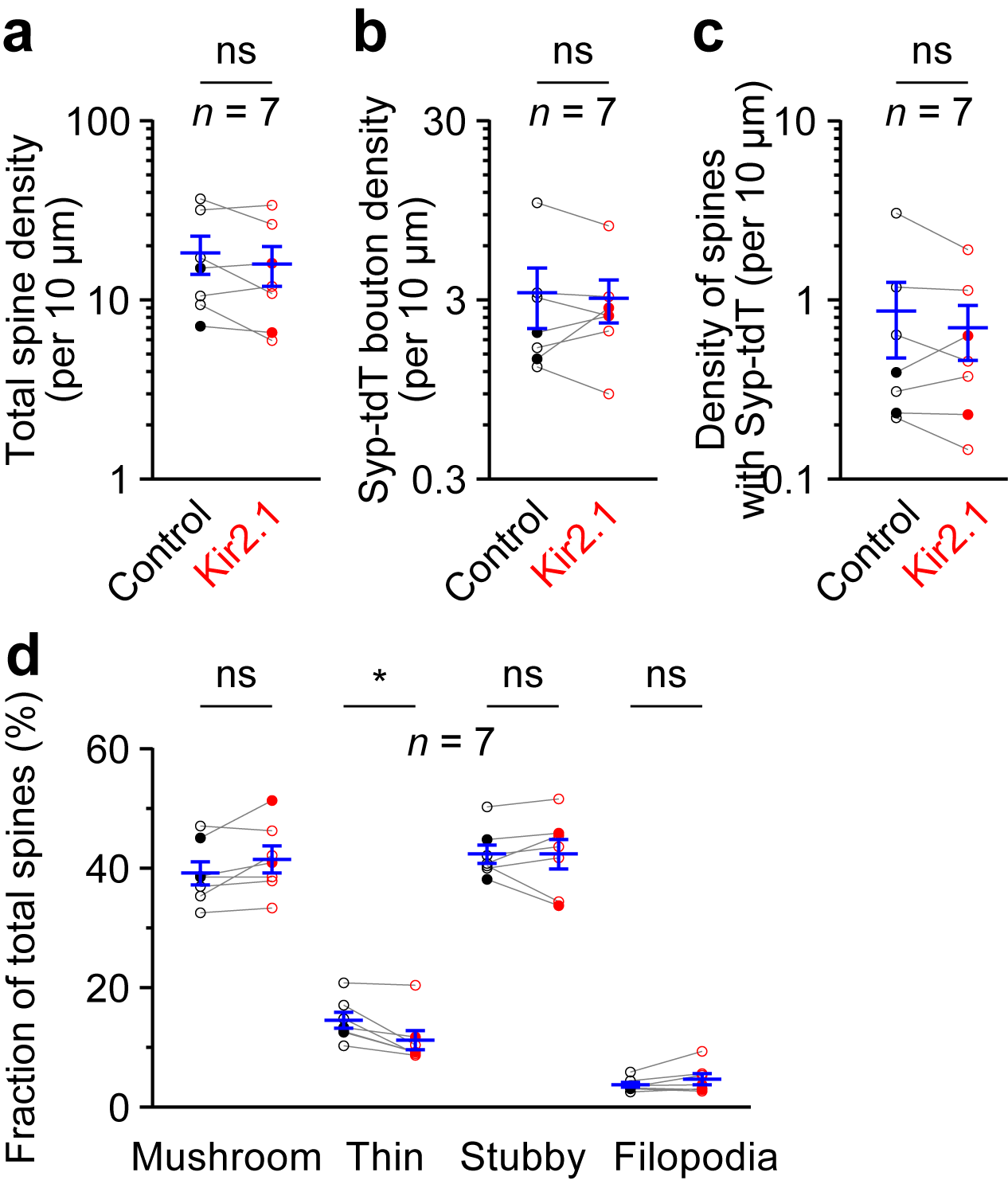

### Extended data F10_Intrinsic excitability of Kir2.1+GluA1CT or A2CT_EPSC of GluA1CT or GluA2CT 20260424.tif

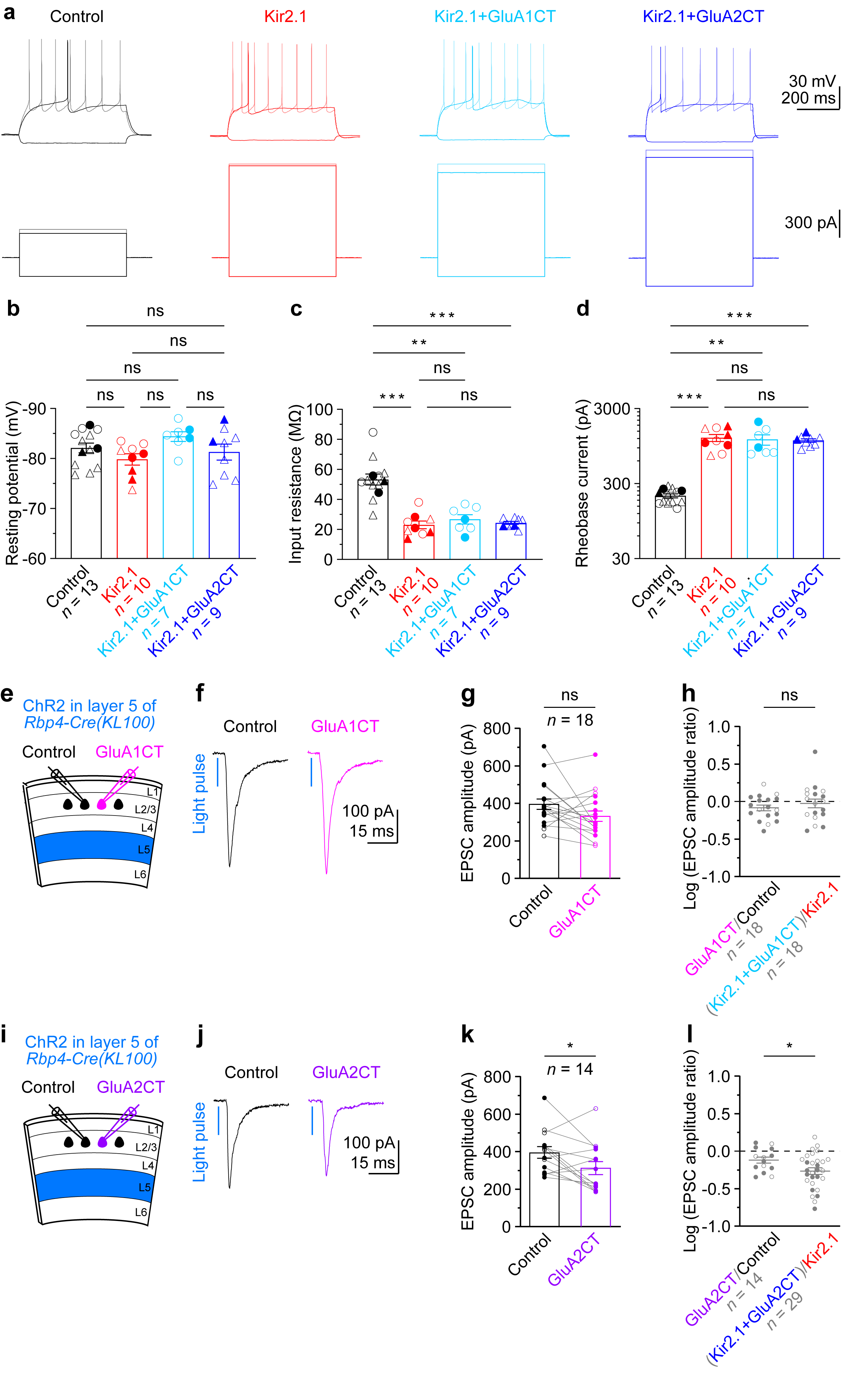
